# Chromosome-level genome sequence of the Genetically Improved Farmed Tilapia (GIFT, *Oreochromis niloticus*) highlights regions of introgression with *O. mossambicus*

**DOI:** 10.1101/2022.05.10.490902

**Authors:** GJ Etherington, W Nash, A Ciezarek, TK Mehta, A Barria, C Peñaloza, MGQ Khan, A Durrant, N Forrester, F Fraser, N Irish, GG Kaithakottil, J Lipscombe, T Trong, C Watkins, D Swarbreck, E Angiolini, A Cnaani, K Gharbi, RD Houston, JAH Benzie, W Haerty

## Abstract

**Background:** The Nile tilapia (*Oreochromis niloticus*) is the third most important freshwater fish for aquaculture. Its success is directly linked to continuous breeding efforts focusing on production traits such as growth rate and weight. Among those elite strains, the Genetically Improved Farmed Tilapia (GIFT) programme initiated by WorldFish is now distributed worldwide. To accelerate the development of the GIFT strain through genomic selection, a high-quality reference genome is necessary.

**Findings:** Using a combination of short (10X Genomics) and long read (PacBio HiFi, PacBio CLR) sequencing and a genetic map for the GIFT strain, we generated a chromosome level genome assembly for the GIFT. Using genomes of two closely related species (*O. mossambicus, O. aureus*), we characterised the extent of introgression between these species and *O. niloticus* that has occurred during the breeding process. Over 11Mb of *O. mossambicus* genomic material could be identified within the GIFT genome, including genes associated with immunity but also with traits of interest such as growth rate.

**Conclusion:** Because of the breeding history of elite strains, current reference genomes might not be the most suitable to support further studies into the GIFT strain. We generated a chromosome level assembly of the GIFT strain, characterising its mixed origins, and the potential contributions of introgressed regions to selected traits.

## Introduction

It is estimated that by 2050 food demand will increase by 36% - 51% [1], with significant challenges such as increased scarcity and reduced quality of land and water sources [2]. This includes ever-increasing demands for animal protein, with a growing importance of aquaculture in this process. Aquaculture production is now nearly on par with wild capture output reaching 46% of the total fish industry yield in 2018 [3]. Inland finfish production represented nearly 47 million tonnes in 2018, with the Nile tilapia (*Oreochromis niloticus*) being the third most important species, representing 8.3% of the global production, [3].

The success of Nile tilapia is partly associated with at least 27 breeding programmes, with efforts that have mainly focused on increasing growth rate to shorten production time. This is exemplified by the Genetically Improved Farmed Tilapia (GIFT) that was initiated in 1987 through a collaboration between the International Center for Living Aquatic Resources Management (ICLARM, now WorldFish Center) and the Institute for Aquaculture Research (AKVAFORSK, Norway). The breeding programme was developed in 1988 from a total of eight strains and populations, four populations from Africa (Egypt, Ghana, Kenya, and Senegal) and four already being farmed in Israel, Singapore, Taiwan, and Thailand [4–6] generating a genetically diverse population used for improved growth rate selection, aiming to increase productivity and decrease production costs [7–9]. This led to an increased body weight of 67%-88% relative to the base population over five generations of selection [10]. The improved GIFT strain is now distributed worldwide, playing a key role in developing countries. To further develop the GIFT strain and to enable future improvements, several studies have been investigating the genetic bases underlying traits of economic interest. As a consequence, Quantitative Trait Loci (QTL) associated with sex determination [11], Tilapia Lake Virus resistance [12, 13], feed efficiency [14], adaptation to oxygen stress [15] in the GIFT strain have now been identified.

All identified QTLs have so far been mapped against the only existing Nile tilapia reference genome from an XX homozygous clonal line [16]. This was first released in 2014 based on short read sequencing leading to relatively low contiguity (Contig N50: 29.3kb [17]). Continuous improvement through long-read sequencing of the same line as in Brawand et al [17] led to a highly contiguous reference genome assembly of the Nile tilapia [16]. This reference assembly is contained within 2,566 contigs with a contig N50 of 3.1 MB anchored to 22 linkage groups. One major assumption made when mapping QTLs from GIFT onto this reference is that there should be little genomic differences between the two genomes. However, this reference was generated from DNA extracted from an inbred clonal line of *O. niloticus* that was first originated from Lake Manzala in Egypt [18], and therefore does not share the same evolutionary history as the GIFT strain, including the inter-population crosses and introgressions from *O. mossambicus* and *O. aureus* [19–21]. The complex breeding history of the GIFT strain implies potential large genomic differences between the reference Nile tilapia genome as exemplified by the different genomic locations associated with sex determination loci between the reference strain (LG1, [22]) and GIFT (LG23, [11]).

The increased efforts in identifying the genetic bases associated with traits of interest in GIFT and the need to apply genomic selection to speed up the improvement process requires the generation of genomic resources specific to the GIFT strain including a high-quality reference genome. Here we present the first genome assembly for the GIFT strain generated using a combination of long and linked reads anchored to linkage groups. We also report *O. mossambicus* and *O. aureus* genomic regions that have introgressed into the GIFT genome.

## Material and Methods

### Tissue collection and HMW DNA extraction

A single male GIFT individual (Pit tag ID: 00075F642B) was euthanized with clove oil (400 mg/litre) at WorldFish (Malaysia). Several tissues were dissected, submerged in absolute ethanol and then flash frozen at −80°C. High molecular weight (HMW) DNA was extracted from 7 mg of testis tissue using the Circulomics HMW Tissue DNA Kit Alpha. This followed a slightly modified version of the Dounce method outlined in the kit handbook v0.16c. The DNA was eluted in a final volume of 500 µL and left at room temperature for 48 hours while mixing twice daily with a pipette using a wide-bore tip. The DNA was quantified using Qubit dsDNA HS (High Sensitivity) Assay Kit (Q32854, Thermo Fisher Scientific).and checked for integrity using the FEMTO Pulse® System (Agilent, P/N M5330AA). From 7 mg of tissue input, we obtained 67 µg of HMW DNA (62% of fragments > 50 kb).

A single male *Oreochromis mossambicus* individual (ID: OmM5) was euthanized using an overdose of MS-222 (tricaine) at the Agriculture Research Organization (Israel). Several tissues were dissected, submerged in absolute ethanol and then flash frozen at −80°C. HMW DNA was extracted from 25 mg of testis tissue using the QIAGEN MagAttract HMW DNA kit, following the standard protocol. Genomic DNA was quantified using the Qubit dsDNA HS (High Sensitivity) Assay Kit (Q32854, Thermo Fisher Scientific). The integrity of the DNA was checked using the FEMTO Pulse System Genomic DNA 165 kb Kit (FP-1002-0275, Agilent Technologies) which determined that 21% of the material was greater than 50kb in size,

All animal procedures were approved by the relevant institution and carried out in accordance with approved guidelines.

### Library preparation

#### 10X Genomics

The Chromium 10x platform (10x Genomics) microfluidic Genome Chip (PN-120216) was used to produce barcoded linked-read libraries from GIFT and O. *mossambicus* HMW DNA using 0.625 ng input into the Chromium™ Genome Library Kit & Gel bead Kit v2 (120258) following the Chromium Genome Reagent Kits Version 2 User Guide (CG00043). Library yield was quantified using the Qubit dsDNA HS Assay and insert size was determined with Agilent 2100 Bioanalyzer High Sensitivity DNA chip (5067-4627, Agilent Technologies). The final libraries were quantified by qPCR (07960204001, Roche Diagnostics Ltd) prior to sequencing. Each library was sequenced on an Illumina HiSeq 4000 using 150 base paired-end reads.

A total of 1.25 ng of the HMW DNA from *O. mossambicus*, at a concentration of 1 ng/uL was used by the Earlham Insitute (EI) Genomics Pipeline to prepare a 10x Genomics Chromium library following the standard protocol. The library was sequenced at EI on one lane of Illumina HiSeq4000, generating 344 million 150-bp paired-end (PE) reads.

#### PacBio HiFi and CLR

A HiFi and CLR library was prepared from 13 µg and 10 µg gDNA respectively. Each sample was manually sheared with the Megaruptor 3 instrument (Diagenode, P/N B06010003) with the parameters appropriate for each library size according to the Megaruptor 3 operations manual. Each sample underwent AMPure® PB bead (PacBio®, P/N 100-265-900) purification and concentration before undergoing library preparation using the SMRTbell® Express Template Prep Kit 2.0 (PacBio®, P/N 100-983-900). The HiFi library was prepared according to the HiFi protocol version 03 (PacBio®, P/N 101-853-100) and the final library was size-fractionated to approximately 18 kb using the SageELF® system (Sage Science®, P/N ELF0001), 0.75% cassette (Sage Science®, P/N ELD7510). The CLR library was constructed according to the instructions in the CLR protocol version 01 (PacBio®, P/N 101-693-800), and size selected to > 30 kb using the BluePippin® system (Sage Science®, P/N BLU0001), 0.75% Agarose cassette high pass program and U1 marker (Sage Science®, P/N PAC30KB). HiFi and CLR libraries were quantified by fluorescence (Invitrogen Qubit™ 3.0, P/N Q33216) and the size of fractions and libraries was estimated from a smear analysis performed on the FEMTO Pulse® System (Agilent, P/N M5330AA).

The loading calculations for sequencing were completed using the PacBio® SMRT®Link Binding Calculator v8.0.0.78867. Sequencing primers v2 and v4 were annealed to the adapter sequence of the HiFi and CLR libraries respectively. The libraries were bound to the sequencing polymerase with the Sequel® II Binding Kit v2.0 (PacBio®, P/N 101-842-900). Calculations for primer and polymerase binding ratios were kept at default values for the respective library types. Sequel® II DNA internal control was spiked into each library at the standard concentration prior to sequencing. The sequencing chemistry used was Sequel® II Sequencing Plate 2.0 (PacBio®, P/N 101-820-200) and the Instrument Control Software v8.0.0.78867.

Each library was sequenced on one Sequel II SMRT®cell 8M. The run parameters for sequencing the HiFi library were the following: diffusion loading, 30-hour movie, 2-hour immobilisation time, 4-hour pre-extension time, 35 pM on plate loading concentration. The run parameters for sequencing for the CLR library: diffusion loading, 15-hour movie, 2-hour immobilisation time, no pre-extension time, 30pM on plate loading concentration. HiFi reads were generated from the output of the HiFi run using SMRT Tools v 8.0.0.80529.

### Genome assemblies

#### GIFT

PacBio HiFi subreads were processed using SMRTLINK (v10.1.0.119588, https://www.pacb.com/support/software-downloads/) to generate HiFi quality reads. First multiple subreads of the same SMRTbell molecule were combined to create highly accurate consensus (HiFi) reads using the ‘pb_ccs’ workflow these were then demultiplexed via the ‘pb_demux_ccs’ workflow, for both steps default parameters were used. The sequencing resulted in 2M reads of PacBio HiFi data with a minimum Q20 and read-length N50 of 15.4 kb, 6M reads of PacBio CLR data with a mean read-length of 18.4 kb and N50 of 29.8 kb, in addition to 468M 10x Genomics linked-read pairs (150-bp). We developed an assembly pipeline to generate assemblies and integrate in a stepwise manner the different data sets generated, assessing assembly quality at each step.

Six different draft assemblies (Figure 1A) were generated and compared to select the best *de novo* assembly.

**Figure 1.**
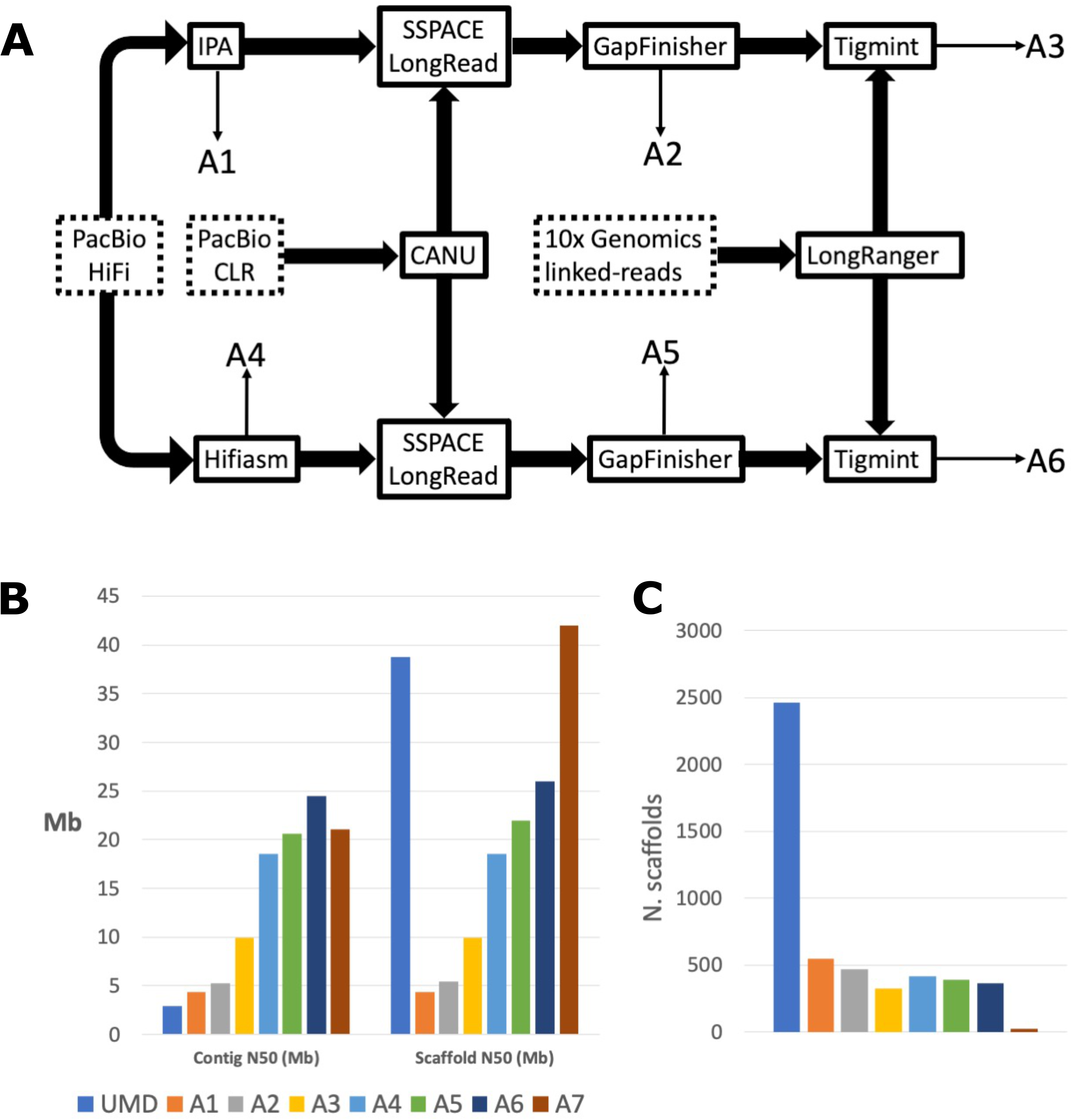
GIFT genome assembly. **A)** Assembly pipeline to combine PacBio HiFi, CLR, and 10X Genomics reads. Input data is denoted by dotted boxes, assembly tools are denoted by solid boxes and output assemblies (A1 – A6) corresponding to those described above are marked by thin arrows. **B)** Comparison of contig sizes between the current *O. niloticus* assembly (UMD, [16]) prior scaffolding and anchoring to the chromosomes, and the successive assemblies produced. **C**) Comparison of scaffold sizes between all assemblies including chromosome anchored assemblies (UMD, A7).

The assemblies were as follows:

A1. PacBio HiFi assembled using PacBio’s Improved Phased Assembly tool (IPA) v1.3.1 (https://github.com/PacificBiosciences/pbipa).
A2. PacBio HiFi assembled using IPA and gap-filled using the PacBio CLR data.
A3. PacBio HiFi assembled using IPA, gap-filled using the PacBio CLR data, and corrected and re-scaffolded using 10x Genomics linked-reads.
A4. PacBio HiFi assembled using HiFiasm [23].
A5. PacBio HiFi assembled using HiFiasm and gap-filled using the PacBio CLR data.
A6. PacBio HiFi assembled using HiFiasm, gap-filled using the PacBio CLR data, and corrected and re-scaffolded using 10x Genomics linked-reads.

Gap-filling with PacBio CLR data was carried out using Canu [24] to correct and trim the reads, followed by SSPACE_LongReads to scaffold the trimmed CLR reads to the IPA/HiFiasm assembly [25], and GapFinisher, which uses the output of SSPACE_LongRead to identify the optimum alignment information to fill gaps [26]. 10x Genomics linked-read scaffolding was carried out by first processing the reads with Longranger [27], and then using Tigmint [28]. Tigmint identifies misassemblies using linked-read information and then reassembles those using ARCS. Each assembly was then soft-masked using RepeatMasker, with ‘cichlidae’ as the reference species [29]. Assemblies were compared based on different quality metrics: Assembly size, contig N50, scaffold N50, and Number of scaffolds. We also used BUSCOv5 to assess the number of single-copy zebrafish orthologs reconstructed in each assembly [30].

### Linkage map scaffolding

We then selected the best *de novo* assembly from A1-A6 and used a linkage map developed from GIFT families to assemble it into linkage groups. To generate the linkage map, 1,325 individuals from 113 full-sibling families were genotyped with a ∼60K SNP array [29]. The fish belonged to a breeding programme managed by WorldFish (Jitra, Malaysia). Fin clips were collected from the fish and preserved in absolute ethanol (Fisher Scientific) until DNA was extracted following [31]. Genotyping with the SNP chip was carried out by Identigen (Dublin, Ireland). Raw intensity files were assessed through the Axiom Analysis Suite Software v4.0.3.3. Genotype calling was performed using default parameter settings for diploid species, except for the sample call rate threshold, which was lowered to 93. Further SNP and sample QC was conducted using Plink v1.09 [32]. Samples with a call rate < 0.95 were removed from the study. Markers with a call rate < 0.99, MAF < 0.05 and HWE p < 1×10-6 were also excluded from the analysis. Additionally, both markers and individuals with an excess of Mendelian errors (> 5 %) were discarded. After QC-filtering, 50,711K SNPs genotyped on 1,133 samples (185 parents and 948 offspring) were available for the construction of a linkage map using Lep-Map2 and Lep-Map3 [33, 34]. The *ParentCall2* module was used to call parental genotypes by taking into account genotype information from parents and offspring. Next, markers exhibiting significant segregation distortion (dataTolerance = 0.001) were filtered out using the *Filtering2* module. SNP markers were assigned to linkage groups using the *SeparateChromosomes2* algorithm with a LOD threshold of 21 and distortionLod = 1. *JoinSingles2All* was used to add unassigned SNPs to existing linkage groups with parameters set as lodLimit = 3 and distortionLod = 1. Markers were ordered three times within each linkage group (*OrderMarkers2* module), and the best order was chosen based on the likelihood of each configuration. The final map comprised 50,707 SNPs anchored to 22 linkage groups, representing the expected karyotype of Nile tilapia.

We used pblat [35] to align the probe sequences used to design the SNP array on the linkage map to the best GIFT *de novo* assembly and examined their distribution across the scaffolds. GIFT scaffolds were then ordered and orientated into their linkage group of origin. Scaffolds without any mapped probes were allocated to LG0. We further refer to this assembly as ‘A7’.

### *O. mossambicus* assembly

The 688M 10x Genomics Chromium link-read pairs were assembled using the 10x Genomics Supernova (v2.1.1) software [36] with default parameters. This resulted in an effective coverage of 64x with a mean molecule length of 38 kb. Supernova initially generates a contig-assembly that is then scaffolded using molecule-specific barcode information [36]. The resulting assembly was 814 Mb in size (scaffolds >=10 kb) and outputted in the ‘pseudohap’ style, randomly creating 1 haplotype per scaffold, with a contig N50 of 38 kb and scaffold N50 of 2 Mb.

### *O. mossambicus* and *O. niloticus* tissue collection, RNA extraction and RNA-seq

Three male *Oreochromis mossambicus* individuals (ID: OmM3, OmM4, and OmM6) from the same line as the genome assembly individual (ID: OmM5), and three male *Oreochromis niloticus* individuals (ID: OnM6, OnM7, and OnM11) from the same Lake Manzala stock used to generate the Nile tilapia reference genome [16], were euthanized using an overdose of MS-222 (tricaine) at the Agriculture Research Organization (Israel). Gill, liver and kidney tissues were dissected, submerged in RNAlater (1:5 ratio) and then flash frozen at −80°C. RNA was purified from each tissue using the RNeasy Plus Mini kit (QIAGEN) according to the manufacturer’s protocol. RNA and DNA content were quantified on the Qubit 4 fluorometer (Invitrogen) and integrity assessed on the Agilent 4200 Tapestation (Agilent Technologies), selecting samples with RIN≥7 and <15% genomic DNA. Stranded RNA-seq libraries were constructed using a combination of poly-A pull down beads from Illumina TruSeq RNA v2 library construction kit (PN: RS-122-2001) and the NEXTflex™ Rapid Directional RNA-Seq Kit (PN: 5138-07) with NEXTflex™ DNA Barcodes – 48 (PN: 514104) diluted to 6 µM. 1 µg of RNA was purified to extract mRNA with a poly-A pull down using biotin beads from the Illumina TruSeq RNA library construction kit. The resulting purified mRNA was processed using the NEXTflex™ Rapid Directional RNA-Seq Kit. Libraries were amplified using 10 cycles of PCR (30 mins at 37°C, 2 mins at 98°C, 10 cycles of 30 sec at 98°C, 30 secs_65°C, and 60 secs at 72°C, followed by a final 4 mins at 72oC). The quality of the resulting libraries was determined using a Perkin Elmer DNA High Sensitivity Reagent Kit (CLS760672) with DNA 1K / 12K / HiSensitivity Assay LabChip (760517) and the concentration measured with a Quant-iT™ dsDNA Assay Kit, high sensitivity assay from ThermoFisher (Q-33120). All stranded RNA-seq libraries were equimolar pooled using qPCR (07960204001, Roche Diagnostics Ltd) and sequenced using 150 base paired-end reads on an Illumina HiSeq 4000 platform, generating an average of 26 million reads per library. These reads were used for genome annotation.

### Genome synteny

We compared the GIFT linkage group assembly to the current *O. niloticus* reference (UMD, [16]) and *O. mossambicus* assemblies by aligning each of them (with the GIFT-derived assembly as the reference genome) using the MUMmer4 nucmer program [37] and visualising with Dot [38].

### Genome Annotation

#### Annotation

To generate the predicted gene models, the GIFT assembly A7 was used as input for Liftoff (v1.5.1),[39]. Liftover was conducted from the reference annotations of the Nile Tilapia [40] and *O. mossambicus* in GFF format with the settings `-s 0.7 -a 0.9 -flank 0.1`. The GFF annotations generated by Liftoff were passed to the `ei-liftover` pipeline (https://github.com/lucventurini/ei-liftover), which was used to calculate F1 scores. Following the calculation of these metrics, gene models in the raw Liftoff output were filtered for results scoring 100% ensuring only models with consistent gene structures between original and transferred models were retained. These models were filtered from the original GFF and the resulting filtered projected annotations were passed, in GFF format, to the final MINOS step (https://github.com/EI-CoreBioinformatics/minos, described below).

The reference annotations of 9 cichlid species (Table 1) in GFF format, and their reference sequences in FASTA format were passed to the `reat homology` pipeline (https://github.com/EI-CoreBioinformatics/reat) to generate a set of gene models from cross-species proteins alignments. The pipeline was run using a custom configuration and scoring file (available at https://github.com/ethering/gift_assembly_paper). The resulting protein homology-based annotation in GFF format, was passed to the final MINOS step.

**Table 1.** Cichlid genomes used for annotation projection with the `reat homology` (https://github.com/EI-CoreBioinformatics/reat) pipeline. All publically available versions were downloaded from Ensembl.

| Species | Genome Version | Citation | Number of<br>PC genes | Number of<br>transcripts |
| --- | --- | --- | --- | --- |
| *Amphilophus*<br>*citrinellus* | Midas_v5 | [63] | 23696 | 32597 |
| *Astatotilapia burtoni* | AstBur1.0 | [17] | 25955 | 52846 |
| *Astatotilapia*<br>*calliptera* | fAstCal1.2 | [64] | 27018 | 42580 |
| *Metriaclima zebra* | M_zebra_UMD2a | [65] | 27187 | 39681 |
| *Neolamprologus*<br>*brichardi* | NeoBri1.0 | [17] | 23596 | 32961 |
| *Oreochromis*<br>*niloticus* | O_niloticus_UMD1 | [16] | 28142 | 75508 |
| *Oreochromis*<br>*mossambicus* | Oreochromis_mossambicus_EIV1 | This study | 37000 | 55099 |
| *Oreochromis aureus* | ASM587006v1 | [66] | 24464 | 60785 |
| *Pundamilia nyererei* | PunNye1.0 | [17] | 23501 | 32924 |

### Expression based annotation liftover

The RNA-seq reads from *O. niloticus* and *O. mossambicus* were submitted to the `reat transcriptome` pipeline to align, assemble and create a set of transcriptome based gene models. The pipeline was run using custom configuration and scoring files, available at https://github.com/EI-CoreBioinformatics/reat. The resulting expression based annotation, in GFF format, was passed to the final MINOS step (https://github.com/EI-CoreBioinformatics/minos).

A *de novo* repeat annotation was created for the A7 assembly using the RepeatModeller(v1.0.11)/RepeatMasker(v4.07) pipeline [29] with defaults settings and the `--gff` output option enabled. The RepeatModeler library was hard masked using the Oreochromis mossambicus EIv1.0 protein coding genes to remove protein coding sequences. The Oreochromis mossambicus EIv1.0 protein coding genes were first filtered to remove any genes with descriptions indicating “transposon” or “helicase”. TransposonPSI (r08222010) was then run to remove any tranposon hits by hard-masking them and using the filtered gene set to mask the RepeatModeler library. Unclassified RepeatModeler repeats were also hard-masked after a BLAST (v2.6.0) search against the organellar genomes (mitochondrial sequences from Actinopterygii [ORGN] in NCBI nucleotide division, downloaded on 27Oct2021). The RepeatMasker GFF output from the above custom RepeatModeler library and the GFF from a RepeatMasker run using the RepBase (release 20170127, Actinopterygii) library were combined to create a final repeat annotation that resulted in 36.73% of the genome being soft-masked. The repeat annotation, in GFF format, was passed to the final MINOS step (https://github.com/EI-CoreBioinformatics/minos).

MINOS was run to aggregate all of the annotation projections described above and create a unified annotation of the A7 genome assembly, models were scored based on congruence with supporting evidence and their intrinsic features (e.g. bonus and penalties relating to size/number of exon, intron, CDS and UTR features). The gene models generated by Liftoff [39] / `ei-liftover`, `reat homology`, and `reat transcriptome` were input as GFF. Evidence used for scoring were the protein fasta files from the reference annotations, the alignment of these protein sequences to the target genome generated by `reat homology`, the RNA-seq mappings generated within the `reat transcriptome` pipeline, the repeat region / interspersed repeat annotations generated with RepeatMasker / RED. Protein fasta files were extracted using gffread-0.12.2 -V -y following samtools-1.11 faidx. In Minos, the final annotation version of GIFT assembly A7 was named ORESP2315963_EIv1.0, corresponding to the ‘Oreochromis sp’ and the NCBI taxon.

### Introgression analysis

To identify regions of the genome supporting different species topologies, we used 250 bp paired-end PCR-free reads for the GIFT*, O. aureus* and *O. mossambicus* specimens. Raw reads from pure wild *O. niloticus* and *O. urolepis* specimens were downloaded from the European Nucleotide Archive (PRJEB36772: ERR4508021, ERR4508023). Adapter sequences and polyG tails were trimmed from raw reads using fastp (v0.20.0;[41]). Trimmed reads were then mapped against both the reference *O. niloticus* UMD assembly and the GIFT genome assembly (same versions as outlined earlier in the manuscript) using bwa mem (v0.7.17;[42]), with mapped reads having mate coordinates and insert size fields added and sorted using samtools (v1.9;[43]). All analyses were carried out on both read mappings separately, but results from the GIFT genome mapping are presented in the main results. SNPs were then called using bcftools (v1.10.2,[44]). The bcftools mpileup function was used, with max-depth per file set to 350. Bcftools call was then used with the multiallelic variant caller. Called SNPs with a total depth of less than 90, more than 900, a minor allele counts of less than 3, more than 1 individual with missing data, indels or SNPs overlapping or within 3bp of a called indels were excluded. SNPs were then phased and imputed using beagle (v4.1;[45]), with a window size of 10,000bp and overlap of 1,000bp. Four-taxon *D* statistics and *f*4 statistics were calculated using Dsuite (v0.3). Data from all linkage groups, except for the repeat rich LG3 were used for this analysis. Following Malinsky et al. [46], correlated *f*4 statistics were used to calculate *f-*branch, to identify the branches of the phylogeny where introgression is most likely to have occurred. To obtain a topology as input for Dsuite, the vcf was pruned for linkage (R2 values > 0.6 over 20kb windows) using bcftools (v1.12). A phylogenetic tree was then inferred using this pruned dataset using iqtree (v2.0) with 1000 rapid bootstrap replicates, 10 independent runs and automated model detection with ascertainment bias correction.

To assess the topological relationships across the genome, the VCF file was converted to a geno format file using genomics_general (https://github.com/simonhmartin/genomics_general), and phylogenetic trees were inferred over sliding 200bp windows, with 40bp overlap, using iqtree (v1.6.12;[47]), with an automated model detection [48] with ascertainment bias correction. This was implemented using a custom version of the genomics_general sliding_windows scripts. We ran TWISST [49] to calculate topology weightings, using the method ‘complete’. Prior to plotting, a loess smoothing parameter was applied (span 0.05). Of the 15 topologies, the frequency of the most frequent topology where GIFT was sister with either *O. urolepis, O. aureus* and *O. mossambicus* were plotted. The average weightings of these three topologies, plus the topology where GIFT was sister to *O. niloticus* were also plotted. Discordance across the highly repetitive mega-chromosome LG3 was investigated further. All 15 topologies were plotted along the length of the chromosome, and the average weighting of each topology across this linkage group was compared to across the other linkage groups combined.

As the previous approach relies on mapping short reads to the *O. niloticus* genome, we applied a complementary approach based on the long reads we generated to identify novel *O. mossambicus* introgressed regions. We investigated our raw sequencing data to identify PacBio HiFi reads that spanned the GIFT linkage group (LG) assembly but were split between the *O. niloticus* UMD reference (GenBank accession GCA_001858045.3, [16]) and *O. mossambicus*. We hypothesise that reads will be split at the boundaries of introgressed blocks, but note that incomplete lineage sorting or mapping difficulties, such as in repeat-rich areas, may generate similar patterns. We used pbmm2 (https://github.com/PacificBiosciences/pbmm2) to map the PacBio HiFi reads to each of the references. We used Samtools (v1.9, [43]) to trim the first and last 10Kb of reads from each scaffold in order to reduce false positives where reads merely span near the end of one reference scaffold and near the start of another. We then identified reads that fully aligned to the GIFT assembly but were split between UMD and *O. mossambicus* and where no part of the read was mapped to both references. We then calculated coverage over 1kb intervals and merged contiguous intervals that contained at least 30x read coverage (roughly the mean read coverage for all three references), identifying the reads within each interval. Starting with GIFT, we took each of these high-coverage intervals in turn and identified which intervals had the highest intersection of read names (and hence shared the most reads that were fully mapped in GIFT Tilapia and split across UMD and *O. mossambicus*). We then calculated the length of the interval and estimated the read-coverage in that region. We also calculated the combined length of all merged contiguous intervals across each LG/scaffold to identify and rank the LGs/scaffolds that contributed most to read splitting.

### Identifying introgressed genes

A total of 11.02Mb of regions of the GIFT genome putatively introgressed from O. *mossambicus* were identified using the TWISST approach. These were defined as regions where the topology most closely resembling the species tree, but with GIFT and *mossambicus* sister to each other had a weighting of 1.0. These regions were intersected with the ORESP2315963_EIv1.0 GIFT annotation using bedtools v2.30.0 [50] (`bedtools intersect -wa -wb`). The sequences of the 423 predicted ORESP2315963_EIv1.0 GIFT genes were extracted (`bedtools getFasta`). A custom blast database was created, using `makeblastdb` (blast+ v2.10.0+-4, [51]), from the ENSEMBL 105 *O.niloticus* CDS fasta file (Oreochromis_niloticus.O_niloticus_UMD_NMBU.cds.all.fa.gz). `blastn` was used to retrieve a single match from the UMD genome for each GIFT gene (`-outfmt 6 -max_target_seqs 1 - max_hsps 1`). The gene IDs from the Ensembl UMD Onil were used to conduct functional enrichment.

### Gene Ontology (GO) enrichment analysis

Since there are few annotated *Oreochromis niloticus* GO terms, we used the closely related *Oryzias latipes* (Japanese medaka HdrR) GO terms for enrichment analysis. The direct orthologs of introgressed *O. niloticus* Ensembl genes in *O. latipes* were obtained using the ‘g:Orth’ module of g:Profiler (https://biit.cs.ut.ee/gprofiler/orth, [52]). GO enrichment analysis was conducted using the ‘g:GOst’ module of g:Profiler (https://biit.cs.ut.ee/gprofiler/gost) version e105_eg52_p16_e84549f (February 2022), using the *O. latipes* database. We use the FDR-corrected hypergeometric *p* value to assess enrichment of GO terms, with a statistical cut-off of FDR < 0.05.

## Results

### Genome assembly of the Genetically Improved Farmed Tilapia (GIFT) strain

We applied a stepwise approach to integrate the different sequencing libraries (10x Genomics, PacBio HiFi, PacBio CLR) and generated six different *de novo* assemblies (A1 - A6, **Figure 1A**). We assessed each assembly using a range of different QC methods (see **Supplementary Table 1**). At each step we compared our assemblies to that of the current *O. niloticus* reference (UMD) both before and after anchoring to chromosomes (**Figure 1B-1C**)[16]. Our stepwise approach, starting with the HiFi data, allowed us to assess the performance of different approaches (IPA vs HiFiasm) and to further increase contiguity through the inclusion of Pacbio CLR reads and 10x Genomics linked reads. From our six *de novo* assemblies (A1-A6), the HiFiasm assembler [23] generated assemblies that were more contiguous than those generated with IPA as the base assembly. Even the most contiguous IPA-based assembly (A3), which used all the available data, was not as contiguous as the base HiFiasm assembly (A4), both in terms of contig N50 and scaffold N50. The UMD assembly had the lowest contig N50 of all assemblies (2.9 Mb), being more than half the size of the next smallest contig N50 (A1, 4.34 Mb). The number of scaffolds in UMD was significantly higher than those of our *de novo* assemblies, having 4.5 times more scaffolds than the assembly with the highest number of scaffolds (A1) and 7.5 times larger than the assembly with the lowest number of scaffolds (A3). UMD’s low contig N50 combined with the high scaffold count and scaffold N50 represents scaffolding of contigs onto a chromosome-scale linkage map, but with 2,437 unplaced contigs representing 9.75% of the total assembly size.

Comparing the initial assemblies created with IPA and HiFiasm, we see contig and scaffold N50 up to 4.34 and 18.5 Mb respectively, which further improved with the addition of PacBio CLR and 10x Genomics linked-read data to contig N50 of 9.88 and 24.5Mb (**Supplementary Table 1**). It is important to note that the improved contiguity of our assemblies prior to anchoring to chromosomes relative to the UMD *O. niloticus* reference reflects the rapid sequencing technology improvements since its release.

Both the UMD reference and all the IPA-based assemblies (A1 – A3) had a span of 1.01 Gb. This increased slightly in the HiFiasm-based assemblies (A4 – A6) and the GIFT LG assembly, which had spans of between 1.06 and 1.07 Gb respectively.

### Linkage group assembly

We used the scaffolds from assembly A6 and a genetic map for the GIFT strain [53] to anchor into linkage groups (LGs, assembly A7). Most scaffolds only had probes mapped from one LG, but 11 scaffolds had probes from two or more LGs, suggesting some misassembled scaffolds in the GIFT assembly. The distribution of the probes across the scaffolds confirmed this, with probes from the same LGs clustering across adjoining regions and not overlapping with probes from other LGs. We therefore broke up these 11 scaffolds, using the locations of the most distantly mapped probes from each LG as break sites. Each GIFT scaffold was ordered and orientated following the genetic location on each LG and then concatenated using 100 Ns to separate each scaffold. To confirm the scaffolds had been joined in the correct order and orientation we re-ran pblat, aligning the probes to the new GIFT LG assembly and created Marley plots of hits for each LG using genetic location on one axis and physical location (in bp) on the other (**Supplementary Figure 1**). Of the 364 scaffolds, we were able to successfully anchor 88 (accounting for 94.3% of the assembly) into 22 linkage groups. The remaining scaffolds were allocated to LG0.

The BUSCO analysis of the resulting assembly shows that 95.5% of the BUSCO genes are found as single copies, and only 1% of the genes are missing, as observed with the *O. niloticus* reference (**Figure 2**).

**Figure 2.**
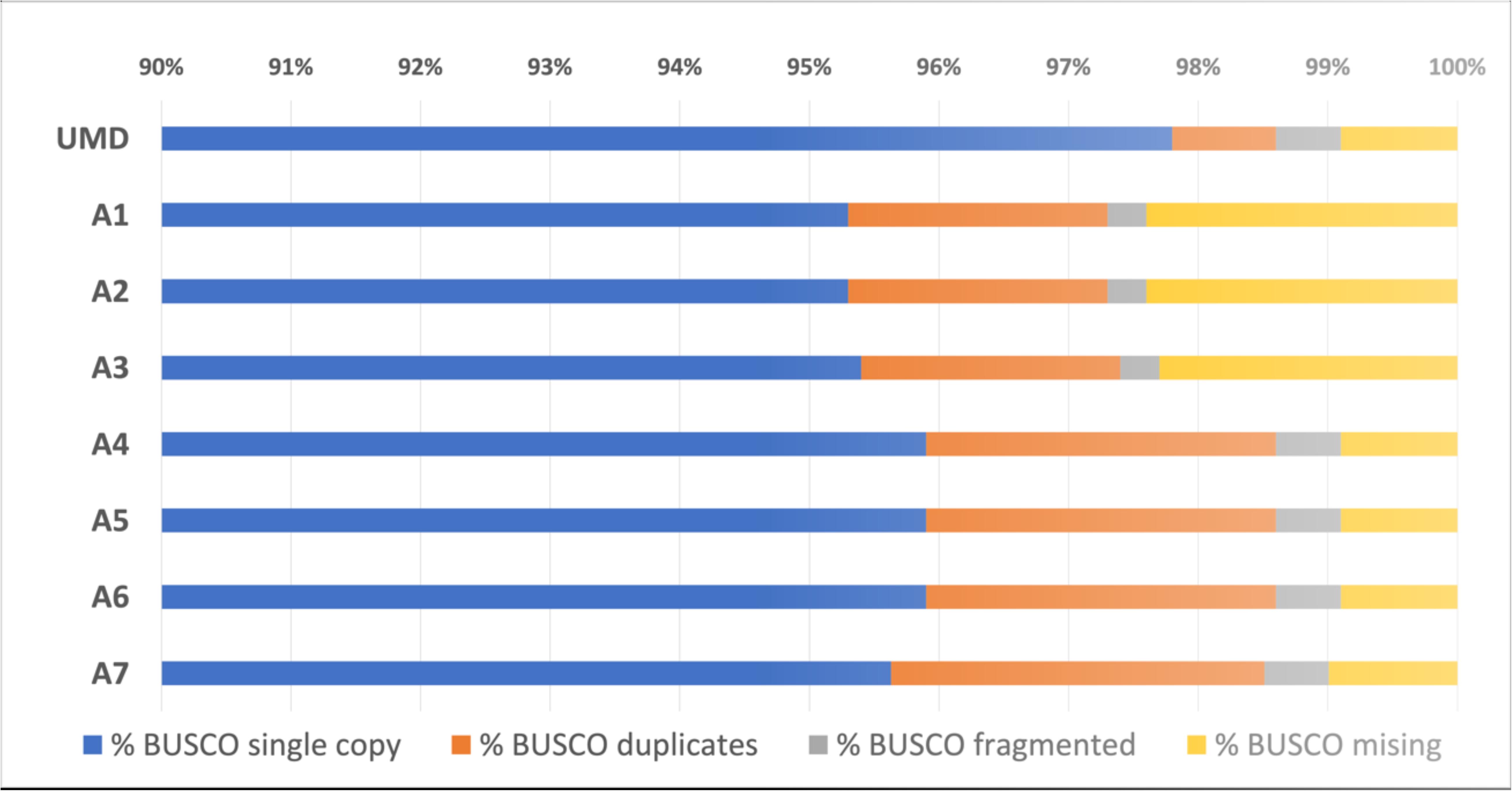
BUSCO hits for eight assemblies – the six assemblies described in the assembly pipeline (A1 – A6), assembly A6 after scaffolding into linkage groups (A7), and the reference Tilapia genome (UMD). The chart starts with single-copy orthologs at 90% as all assemblies recovered at least 95.3%.

Except for LG3 and LG23, the reference *O. niloticus* genome and the GIFT assembly show excellent synteny (**Figure 3**). For LG3 the lack of alignment between the two assemblies is likely the result of the increased repeat content within this LG as previously noted by Conte et al. [16] as well as its origin. LG3 has been suggested to be the result of an ancient chromosome fusion event identified by cytogenetics [54] and genome [55] analyses. The difference for LG23 is of specific interest as the sex determination locus for the GIFT strain was identified on this chromosome [11].

**Figure 3.**
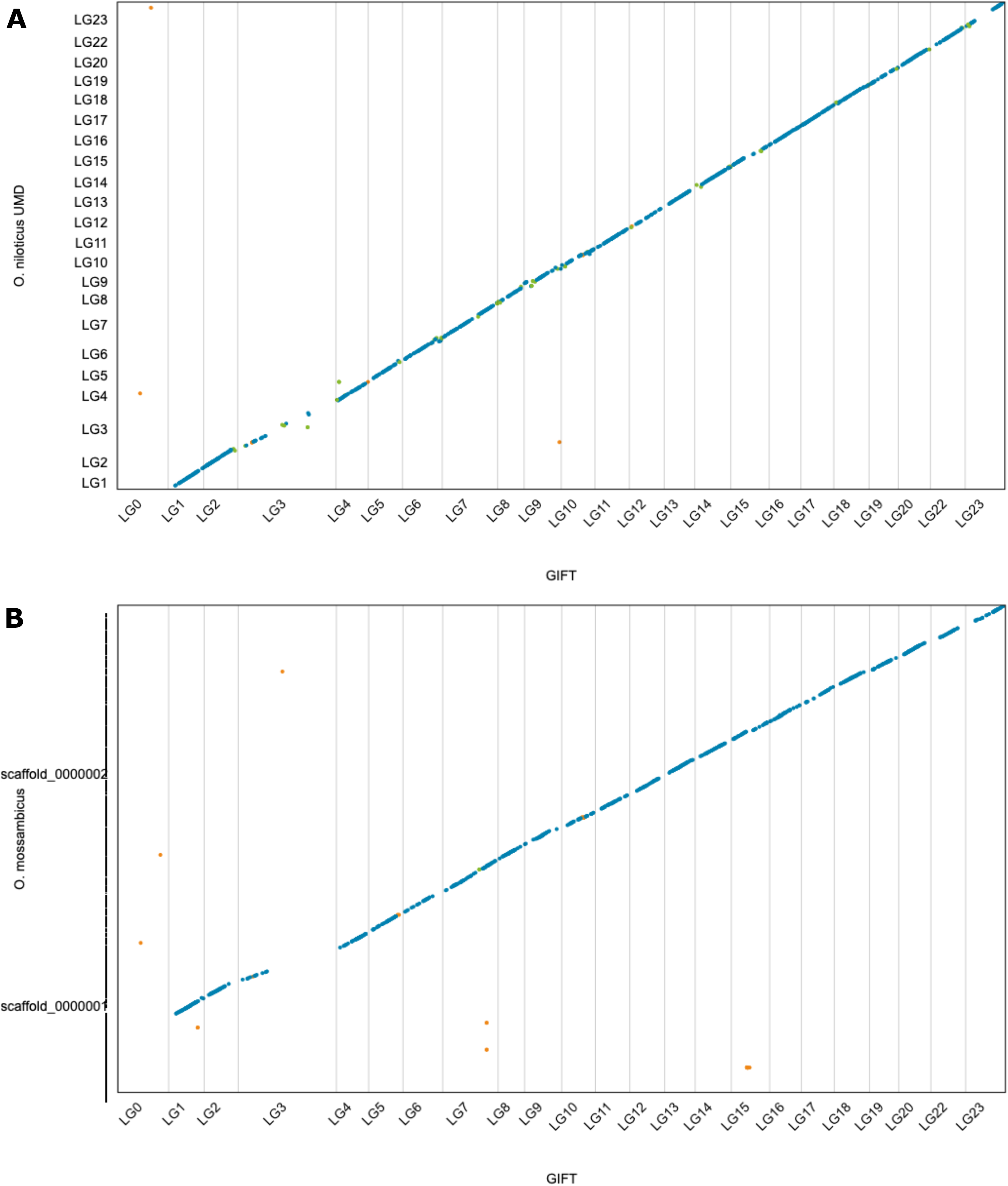
Alignments between *O. niloticus* GIFT and *O. niloticus* UMD (A), and *O. niloticus* GIFT and *O. mossambicus* (B). Blue dots represent unique forward alignments, green dots represent unique reverse alignments, and orange dots represent repetitive alignments.

Using the final assembly, we predicted a total of 31,508 high confidence genes and 59,220 transcripts for the GIFT strain (Table 2).

**Table 2.** The number of genes and transcripts contained in the final ORESP2315963_EIv1.0 annotation of GIFT assembly A7. Confidence level is assigned within the Minos pipeline, with high confidence gene models being supported by multiple sources of evidence, and low confidence gene models being supported by only a single source of evidence.

| Biotype | Confidence | Gene | Transcript |
| --- | --- | --- | --- |
| Protein coding gene | High | **31508** | **59220** |
| Transposable element gene | High | 6281 | 6429 |
| Protein coding gene | Low | 3467 | 3980 |
| Transposable element gene | Low | 2624 | 2723 |
| Predicted gene | Low | 1946 | 1965 |
| Total |  | 45826 | 74317 |

### Introgression

We identified a difference of almost 50Mb between the genome size of GIFT and the *O. niloticus* UMD reference. When comparing the repeat content of the two assemblies, we observed about 2.3Mb extra repeat content in GIFT (27.3Mb vs 25Mb respectively in GIFT and the *O. niloticus* reference). We also assessed the potential impact of the breeding history of the GIFT strain by investigating the genome-wide presence of introgressed regions from *O. mossambicu*s within the GIFT genome.

We calculated intervals in GIFT where reads were fully mapped but split (and not overlapping) between *O. niloticus* UMD and *O. mossambicus* (**Supplementary Table 2**). These are regions that may putatively be introgressed but may also be associated with incomplete lineage sorting or lower mapping quality. We identified 1,114 intervals in GIFT that shared at least one read of this description. Of these, 191 shared at least 10 reads per 1kb interval and were not from LG0 (unplaced scaffolds). To identify the LGs with the most read splitting, we ranked the LGs of these 191 intervals by how often they occurred. LG3 has by far the largest number of reads split between *O. niloticus* UMD and *O. mossambicus*, having more than 8 times more intervals and reads than any other LG. Additionally, the length of regions associated with read splitting (calculated from the combined length of intervals) was more than 11 times greater in LG3 (4.97Mb), than the next largest LG (0.45Mb in LG6), (**Supplementary Table 2**).

We observed significant variation in the tree topologies along different regions of the genome, both in the mapping against the GIFT genome (**Figure 4a**), as well as to the *O. niloticus* genome (**Supplementary Figure 2**), which gave very similar results. GIFT appears to be most closely related to *O. niloticus*, with the average topology weighting of this relationship exceeding 0.7 genome-wide. The topology with *O. aureus* sister to GIFT was the next most common, with an average topology weighting of 0.1. This showed small peaks across the genome, with each linkage group having regions exceeding 0.27, and 13 linkage groups with regions exceeding 0.5 in the smoothed data. The topology with *O. mossambicus* sister to GIFT had an average topology of weighting 0.03, and showed a markedly heterogeneous pattern across the genome, with distinct peaks in LGs 8, 12, 13, 14, 15, 18, 19 and 23, and low weightings apart from these peaks. The topology with *O. urolepis* sister to GIFT had an average topology weighting of 0.008, with low weightings across the genome. There peaks were far smaller (none exceeding 0.15) than the topologies where *O. mossambicus* or *O. aureus* were sister to GIFT, although there is some overlap in where these small peaks are with the larger peaks in the *O. mossambicus* comparison (e.g. LGs 8, 12, 18, 23).

**Figure 4.**
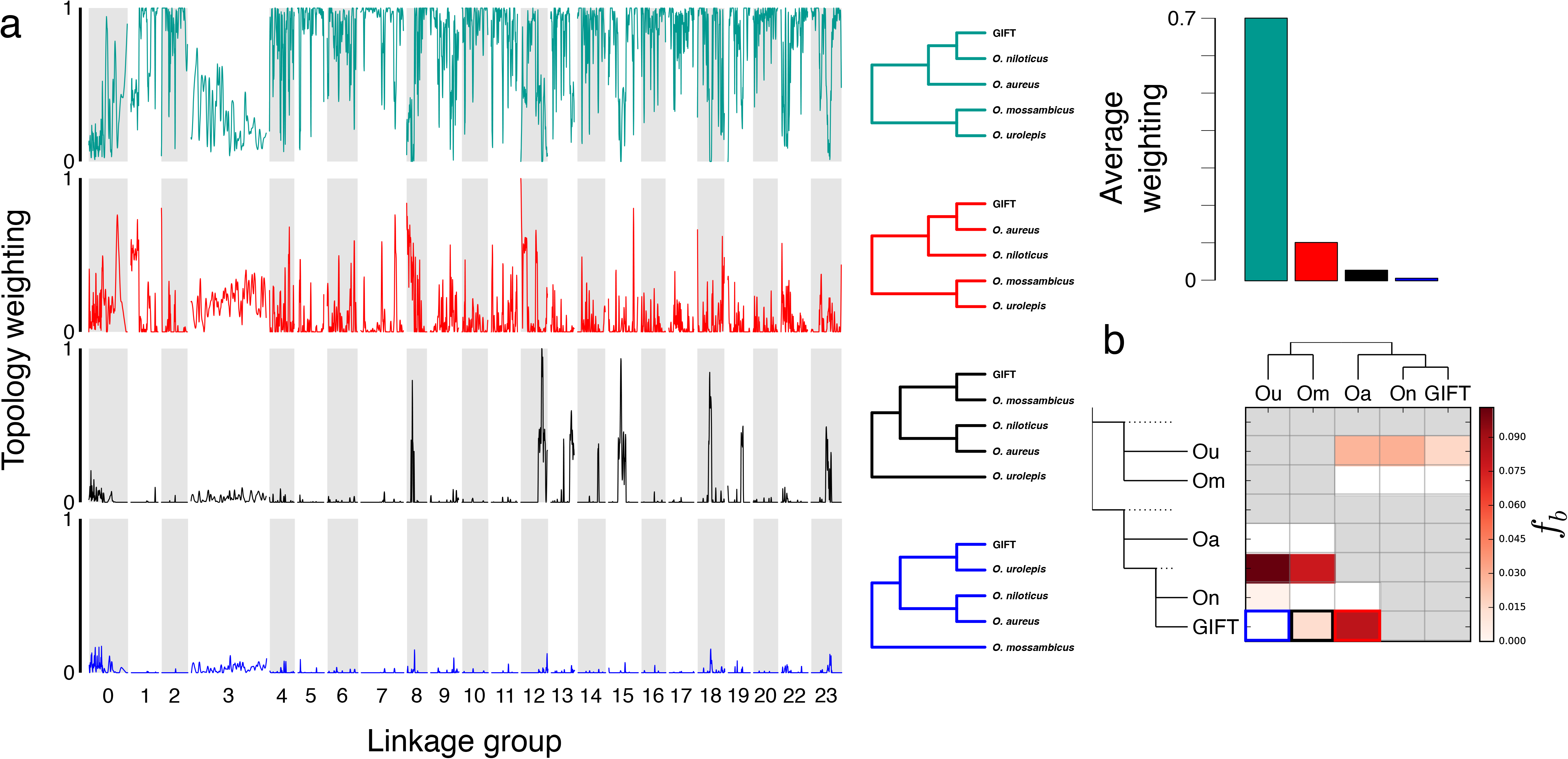
Phylogenetic representation across genomes of species as estimated by TWISST. **A)** Relative weighting of each phylogeny across the GIFT genome assembly. The colours refer to the phylogenies provided in panel B. **B)** Normalized weights across the *O. niloticus* UMD genome assembly for the three different phylogenies.

Overall, the large LG3 appears to be notably discordant, in both TWISST and the read-splitting analysis. However, examination of discordant topologies shows no clear evidence of any specific introgression events in LG3; the species-tree shows a relatively low weighting (0.36 of its average weighting in the rest of the genome) throughout and all discordant topologies have elevated weightings compared to their genome-wide average, without a clear excess of one in particular (**Supplementary Figure 3**). All 14 discordant topologies had average topology weightings higher than the rest of the genome, with 9 of them having the weighting increased between 5-8 fold. The *mossambicus*-GIFT topology was the least elevated of the discordant topologies, with the weighting only 4% higher than in the rest of the genome (**Supplementary Figure 3**). Overall, this is more consistent with the pattern expected from widespread incomplete lineage sorting than introgression.

Genome-wide introgression from *O. mossambicus* and *O. aureus* into the GIFT genome was further supported by *D* and *f-*branch statistics. All trios tested with the *D* statistic were significant, except for the test of introgression from *O. urolepis* into either *O. niloticus* or GIFT in the GIFT mapping dataset. The *f-*branch analyses from the GIFT genome mapping analysis indicated a few different introgression events contributed to this pattern; from both *O. urolepis* and *O. mossambicus* into the ancestor of *O. niloticus* and GIFT, as well as from *O. aureus* into GIFT (*f-*branch = 0.083; indicating around 8.3% admixture proportion) and from *O. mossambicus* into GIFT (*f-*branch = 0.014; around 1.4% admixture proportion). The results from the *O. niloticus* mapping gave similar results (**Supplementary Figure 2**), but with higher *f-*branch estimates (0.153 between *O. aureus* and GIFT; 0.051 between *O. mossambicus* and GIFT)

In total, we identified 283 ORESP2315963_EIv1.0 predicted genes in 11.02 Mb of introgressed regions in the GIFT genome from *O. mossambicus* (**Supplementary Table 3**). GO analyses of the 283 genes against the closely related *O. latipes* (Japanese medaka) database (see Material and Methods), identified enrichment of 71 biological processes (**Figure 5**), 4 molecular functions, 11 cellular components and 5 KEGG pathways (**Supplementary Figure 4, Supplementary Table 4**) terms. The most significant enrichment was that of 41 genes (*adj. p-value* 3.3e-53, FDR < 0.05) associated with phagocytosis (GO:0006910) (**Figure 5**) and concurrently, 8 of those genes in the phagosome KEGG pathway (KEGG:04145) (**Supplementary Figure 4**). This includes tubulin beta chain (*tubb)* and major histocompatibility complex class I genes that play an important role in phagocytosis and immune response to bacterial *Aeromonas hydrophila* infection [56]. Alongside a significant enrichment of genes associated with immune response, we also identified enrichment of 93 genes (*adj. p-value* 1.4e-12, FDR < 0.05) associated with signalling (GO: 0023052) (**Figure 5**), including the pro-opiomelanocortin (*pomc*) gene, that contributes to sexual size dimorphism in tilapia [57].

**Figure 5.**
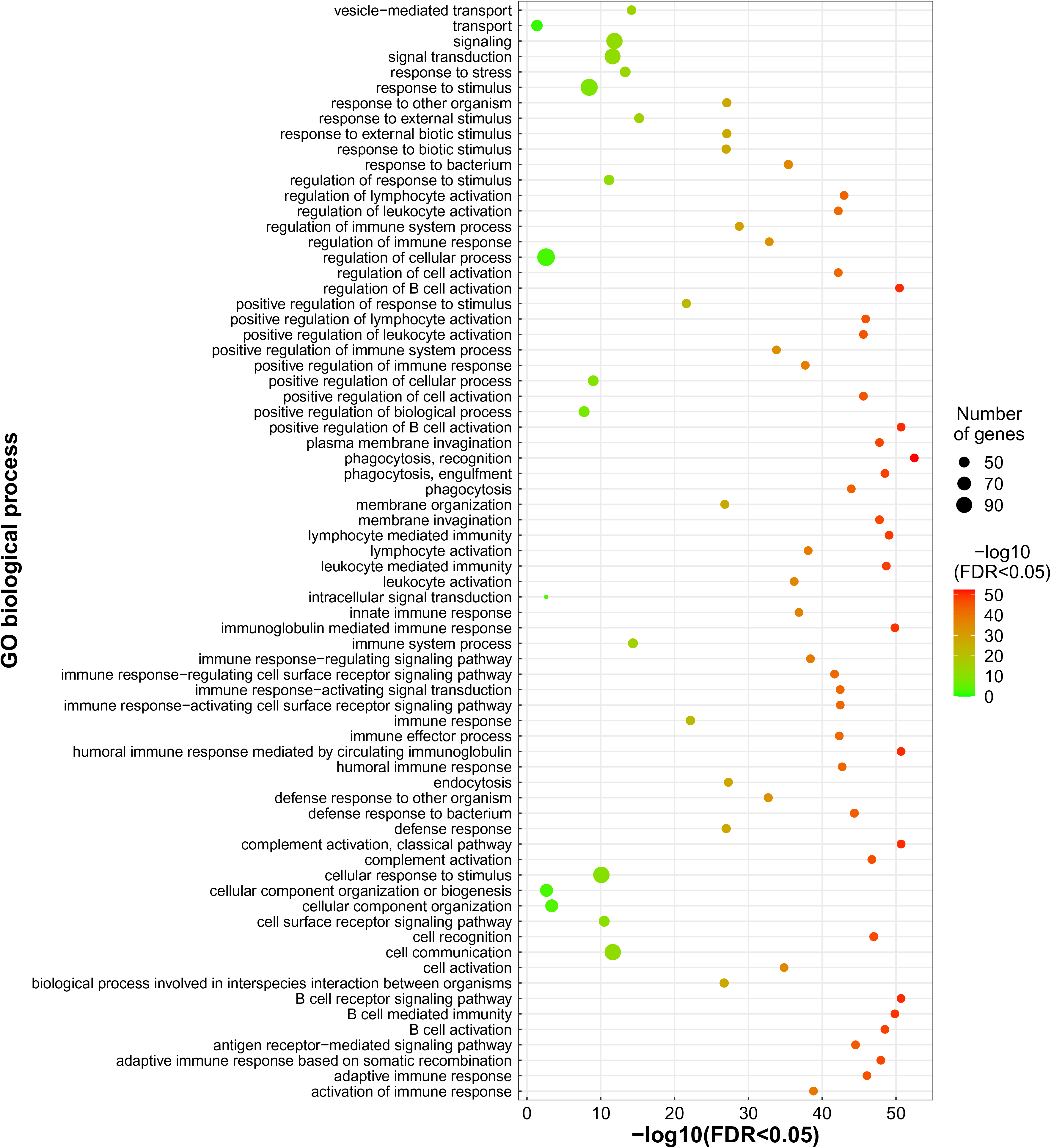
Gene ontology (Biological Processes) enrichment for the genes identified within *O. mossambicus* introgressed regions

## Discussion

The GIFT strain is the first Nile tilapia population to have undergone genetic improvement through selective breeding for survival and increased growth rate. This strain has now been distributed in at least 14 countries across five continents. Because of its distinct breeding history which includes the use of eight *O. niloticus* populations (four from Africa and four already used in South East Asia) and crosses with *O. mossambicus*, the current high-quality reference for the Nile tilapia [16] might not be the best resource to map QTL, identify causative variants and develop novel markers for genomic selection in GIFT. This situation was highlighted by the identification of different chromosomal locations for the sex determination loci in the UMD reference (LG1) and GIFT (LG23) [11, 16].

The combined use of PacBio and 10X Genomics data has generated a high-quality genome assembly for GIFT. We have assembled a highly contiguous assembly (contig N50 – 24.5 Mb, scaffold N50 – 26 Mb, and 364 scaffolds), which is also almost complete (99.1% of BUSCO orthologs, of which 95.9% were present as single-copies). There appears to be a small trade-off between contiguity and haplotype purging and this might explain the slightly larger assembly size (1.07 Gb compared to 1.01 Gb in UMD and IPA-based assemblies), along with the slightly higher number of duplicated BUSCO orthologs (2.7% compared to 2.0% in IPA-based assemblies).

This high-quality assembly has allowed us to examine the genomic impact of introgressions between the selected *O. niloticus* strains and other *Oreochromis* species during the breeding process of GIFT. By leveraging the high-quality PacBio HiFi long reads, we have identified an increase of nearly 50Mb in genome content between our assembly and the *O. niloticus* reference genome. Whilst a significant proportion of the 50Mb difference could be due to technological differences of genome sequencing and assembly approaches, some of these sequences harbour introgressed regions from *O. aureus* or *O. mossambicus*. As such, we have identified a total of 11Mb of genomic space in GIFT that are putatively the result of introgressions from *O. mossambicus* (**Supplementary Table 2**).

While introgressions from *O. mossambicus* have previously been described using mitochondrial or microsatellite markers [19,20,58,59], our work provides a more detailed characterization of the extent of introgressed genomic material. This new assembly and the characterization of these introgressed regions also opens the opportunity to investigate their contributions to the selected traits during breeding.

We identified 283 genes within the introgressed regions, including genes affecting traits potentially relevant for aquaculture, such as sexual size dimorphism [57], defence response to bacterial infection [56] and immune response [60, 61]. More specifically, the pro-opiomelanocortin (*pomc*) gene, has been identified as expressed in the brain and peripheral tissues in cichlid fishes and associated with sexual dimorphism and regulation of social behaviour in *Astatotilapia burtoni* [62]. In *O. mossambicus*, females display a greater expression level of *pomc* in the brain relative to males [57]. The knockout of *pomc* in zebrafish was associated with increased length and body weight in the mutants compared to wild type individuals as well as decreased feed latency in the mutants [57]. Introgression from *O. mossambicus* and *O. aureus* into the GIFT genome may have enabled selection of favourable genes implicated in traits of growth and immunity. Positive selection of growth and immune traits and their associated genes are important for the success of breeding farmed Nile tilapia [61].

This novel genome for the GIFT strain will enable to accurately remap previously identified QTLs for Tilapia Lake Virus resistance [12, 13], feed efficiency [14], and adaptation to oxygen stress [15] that were previously identified using the *O. niloticus* reference genome. This novel genome assembly, including the associated annotations, will enable the identification of causative variants associated with those traits. This novel assembly will enable the generation of novel genomic, epigenomic, and transcriptomic resources leading to a refined annotation of both coding and non-coding functional regions of the genome, enabling variant characterisation and prioritisation. Finally, these novel resources will prove essential in genomic selection programmes aimed at driving the genetic improvement of the GIFT strain and other tilapia species.

## Data availability

All sequencing data generated for the GIFT assembly, along with the final linkage group assembly can be found under ENA study accession PRJEB48957. All scripts are available on the github repository: https://github.com/ethering/gift_assembly_paper.

## Author contributions

WH and JAHB designed the study. GJE generated the GIFT assembly and ran the introgression analysis based on long reads. WN generated the annotation of the GIFT strain and the analysis of the genes within introgressed regions. AC performed the TWISST and short-read based introgression analysis. AC prepared the *O. mossambicus* and *O. niloticus* tissues. TKM extracted HMW DNA for O. mossambicus, generated the *O. mossambicus* assembly and ran GO enrichment analysis. CW and KG designed the sequencing strategy and oversaw data production. TKM, NF and JL extracted RNA from *O. mossambicus* and *O. niloticus.* GK and DS annotated the *O. mossambicus* genome. AD performed HMW extraction from GIFT. FF prepared 10X linked reads libraries. NI prepared HiFi and CLR libraries. AB, CP, MGQK, and RDH developed the genetic map. TT prepared the GIFT tissues. EA project support and management of NF. The manuscript was prepared by WH, GJE, WN, AC, and TKM.

## Acknowledgments

The author(s) acknowledge the support of the Biotechnology and Biological Sciences Research Council (BBSRC), part of UK Research and Innovation; this research was funded by the BBSRC Core Strategic Programme Grant BB/CSP1720/1 and its constituent work packages (BBS/E/T/000PR9818 and BBS/E/T/000PR9819), and the Core Capability Grant BB/CCG1720/1 and the National Capability at the Earlham Institute BBS/E/T/000PR9816 (NC1 - Supporting EI’s ISPs and the UK Community with Genomics and Single Cell Analysis), BBS/E/T/000PR9811 (NC4 - Enabling and Advancing Life Scientists in data-driven research through Advanced Genomics and Computational Training), and BBS/E/T/000PR9814 (NC 3 - Development and deployment of versatile digital platforms for ‘omics-based data sharing and analysis). The authors would like to acknowledge the Scientific Computing group, as well as support for the physical HPC infrastructure and data centre delivered via the NBI Research Computing group.

## Supplementary Tables and Figures

**Supplementary Figure 1.** Comparison of genomic and genetic positions of the SNP markers of the genetic map.

**Supplementary Figure 2.** Phylogenetic representation across genomes of species as estimated by TWISST. **A)** Relative weighting of each phylogeny across the *O. niloticus* UMD genome assembly. The colours refer to the phylogenies provided in panel B. **B)** Normalized weights across the *O. niloticus* UMD genome assembly for the three different phylogenies.

**Supplementary Figure 3. A).** Phylogenetic representation across LG3 of species as estimated by TWISST. **B)** Normalized weights across LG3 for the different phylogenies.

**Supplementary Figure 4.** Gene Ontology analysis (Molecular Function, Cellular Components, KEGG Pathway) for the genes identified within *O. mossambicus* introgressed regions.

**Supplementary Table 1.** Assembly statistics.

**Supplementary Table 2.** Genomics regions in GIFT with evidence of *O. mossambicus* introgression

**Supplementary Table 3**. Genes identified within *O. mossambicus* introgressed regions including corresponding orthologs in medaka

**Supplementary Table 4:** Ontology terms significantly enriched among genes identified within *O. mossambicus* introgressed regions.

