## Supplementary material for "Chromosome-level genome sequence of the Genetically Improved Farmed Tilapia (GIFT, *Oreochromis niloticus*) highlights regions of introgression with *O. mossambicus*": Supplementary_Figure_1.docx

**Supplementary Figure 1:** Comparison of genomic and genetic positions of the SNP markers of the genetic map.

LG1

|  | genetic_pos |
| --- | --- |
| 237466 | 12.46 |
| 350048 | 12.46 |
| 351036 | 12.91 |
| 357853 | 12.46 |
| 359726 | 12.46 |
| 360178 | 12.46 |
| 459443 | 12.46 |
| 459887 | 12.91 |
| 489683 | 12.82 |
| 502949 | 12.91 |
| 518235 | 12.46 |
| 522605 | 12.46 |
| 551186 | 12.91 |
| 552605 | 12.91 |
| 563971 | 12.46 |
| 576742 | 11.31 |
| 579477 | 12.46 |
| 584204 | 13.18 |
| 596448 | 12.91 |
| 623464 | 13.63 |
| 626922 | 12.46 |
| 633166 | 13.09 |
| 645131 | 13.18 |
| 647667 | 13 |
| 674648 | 13.09 |
| 679711 | 13.09 |
| 696500 | 13.09 |
| 710932 | 12.82 |
| 730071 | 13.18 |
| 734633 | 12.91 |
| 735893 | 13.09 |
| 746707 | 12.46 |
| 747194 | 11.61 |
| 749574 | 12.91 |
| 755796 | 13.63 |
| 760662 | 13.09 |
| 764431 | 12.19 |
| 792021 | 12.82 |
| 860927 | 13.09 |
| 863191 | 13.63 |
| 882622 | 12.46 |
| 884423 | 12.55 |
| 884552 | 13.27 |
| 887028 | 13.36 |
| 892507 | 13.09 |
| 905837 | 12.91 |
| 909257 | 12.82 |
| 911564 | 13.09 |
| 914693 | 12.46 |
| 918048 | 13.63 |
| 942919 | 13.09 |
| 948046 | 13.72 |
| 954420 | 12.55 |
| 957466 | 13.09 |
| 960086 | 12.46 |
| 965569 | 12.91 |
| 966773 | 12.46 |
| 985827 | 13.09 |
| 1003492 | 12.46 |
| 1004114 | 9.23 |
| 1005148 | 13.27 |
| 1014647 | 11.91 |
| 1035673 | 11.91 |
| 1040732 | 12.82 |
| 1041361 | 12.46 |
| 1074563 | 13.72 |
| 1092140 | 12.46 |
| 1106010 | 12.46 |
| 1133775 | 12.46 |
| 1134646 | 12.46 |
| 1300837 | 12.46 |
| 1308820 | 13.63 |
| 1318874 | 12.46 |
| 1332705 | 12.46 |
| 1367802 | 12.91 |
| 1391144 | 12.46 |
| 1395755 | 12.82 |
| 1397798 | 12.55 |
| 1398596 | 13.63 |
| 1399975 | 13.63 |
| 1403118 | 13.72 |
| 1404389 | 15.24 |
| 1412537 | 13.63 |
| 1421060 | 12.91 |
| 1426996 | 13.63 |
| 1442341 | 13.63 |
| 1463154 | 13.63 |
| 1503142 | 13.63 |
| 1539632 | 12.46 |
| 1629482 | 13.63 |
| 1636878 | 13.09 |
| 1645300 | 13.63 |
| 1658596 | 13.63 |
| 1673468 | 11.91 |
| 1685389 | 13.27 |
| 1689735 | 13.63 |
| 1694664 | 12.46 |
| 1707424 | 13.36 |
| 1713639 | 9.23 |
| 1716968 | 12.09 |
| 1725618 | 13.36 |
| 1740800 | 12.46 |
| 1752876 | 12.91 |
| 1785080 | 12.46 |
| 1843403 | 10.63 |
| 2201790 | 13.09 |
| 2203249 | 12.46 |
| 2214525 | 11.61 |
| 2216965 | 12.46 |
| 2217400 | 13.63 |
| 2225061 | 12.55 |
| 2227341 | 12.46 |
| 2245235 | 12.46 |
| 2263143 | 11.31 |
| 2267405 | 12.46 |
| 2270988 | 12.46 |
| 2278974 | 12.46 |
| 2286141 | 11.91 |
| 2287062 | 12.46 |
| 2638197 | 12.46 |
| 2640024 | 12.46 |
| 2643099 | 12.46 |
| 2648889 | 12.46 |
| 2663248 | 13.09 |
| 2666822 | 13.63 |
| 2670347 | 13.09 |
| 2670873 | 13.09 |
| 2675795 | 13.09 |
| 2679794 | 12.09 |
| 2683730 | 12.46 |
| 2689969 | 12.46 |
| 2709805 | 13.27 |
| 2714063 | 12.46 |
| 2721448 | 12.46 |
| 2741599 | 12.46 |
| 2744863 | 13.63 |
| 2752099 | 12.46 |
| 2753579 | 13.72 |
| 2756919 | 13.09 |
| 2761082 | 12.46 |
| 2766121 | 12.82 |
| 2794426 | 13.09 |
| 2798514 | 12.91 |
| 2803560 | 13.09 |
| 2811473 | 12.46 |
| 2815988 | 13.09 |
| 2824242 | 12.46 |
| 2835494 | 13.72 |
| 2843623 | 12.91 |
| 2853856 | 12.46 |
| 2871516 | 12.91 |
| 2888544 | 12.46 |
| 2891353 | 12.46 |
| 2893055 | 12.91 |
| 2894926 | 12.46 |
| 2898800 | 13.09 |
| 2900212 | 13.09 |
| 2911248 | 11.91 |
| 2912192 | 12.46 |
| 2921056 | 12.91 |
| 2928866 | 12.46 |
| 2932890 | 12.46 |
| 2935285 | 12.46 |
| 2938811 | 12.82 |
| 2941542 | 12.46 |
| 2943156 | 12.46 |
| 2947376 | 12.91 |
| 2951016 | 12.91 |
| 3006012 | 12.46 |
| 3011998 | 12.82 |
| 3013827 | 12.46 |
| 3022873 | 13.36 |
| 3045630 | 12.46 |
| 3046245 | 12.91 |
| 3067166 | 12.46 |
| 3083468 | 12.55 |
| 3095456 | 12.91 |
| 3098246 | 12.46 |
| 3102238 | 12.46 |
| 3114874 | 12.46 |
| 3135483 | 12.91 |
| 3140337 | 12.46 |
| 3142459 | 12.46 |
| 3172670 | 12.91 |
| 3191025 | 12.46 |
| 3195784 | 12.46 |
| 3205787 | 12.91 |
| 3210168 | 12.46 |
| 3216852 | 12.46 |
| 3219467 | 12.46 |
| 3224591 | 13 |
| 3249647 | 12.46 |
| 3265807 | 12.46 |
| 3273466 | 12.46 |
| 3274669 | 12.46 |
| 3284866 | 12.46 |
| 3292335 | 13.63 |
| 3311135 | 13.09 |
| 3319357 | 12.46 |
| 3323110 | 12.46 |

LG2

|  | genetic_pos |
| --- | --- |
| 254233 | 4.92 |
| 261275 | 6.38 |
| 288995 | 6.38 |
| 290440 | 6.38 |
| 293282 | 6.38 |
| 296940 | 6.38 |
| 303386 | 6.38 |
| 303835 | 6.38 |
| 385312 | 6.38 |
| 423631 | 6.38 |
| 470036 | 6.38 |
| 496458 | 6.38 |
| 504799 | 6.38 |
| 955525 | 6.38 |
| 965143 | 6.38 |
| 1182839 | 6.38 |
| 1189363 | 6.38 |
| 1190997 | 6.38 |
| 1193535 | 6.38 |
| 1205547 | 6.38 |
| 1206582 | 6.03 |
| 1249088 | 6.38 |
| 1267297 | 6.15 |
| 1417462 | 6.38 |
| 1419802 | 3.82 |
| 1420512 | 6.38 |
| 1444459 | 6.38 |
| 1674930 | 6.38 |
| 1797519 | 6.38 |
| 1867648 | 6.49 |
| 1868756 | 6.72 |
| 2192435 | 6.49 |
| 2204647 | 6.49 |
| 2235188 | 6.49 |
| 2241672 | 6.49 |
| 2247303 | 6.49 |
| 2286046 | 6.49 |
| 2315355 | 6.49 |
| 2325228 | 6.49 |
| 2342594 | 6.49 |
| 2348586 | 6.38 |
| 2368553 | 6.49 |
| 2371626 | 6.49 |
| 2380638 | 6.49 |
| 2394020 | 6.49 |
| 2617136 | 6.38 |
| 2627713 | 6.61 |
| 2633752 | 6.61 |
| 2676570 | 6.61 |
| 2716785 | 6.38 |
| 2730299 | 6.72 |
| 2759897 | 6.61 |
| 2777383 | 6.72 |
| 2786003 | 6.61 |
| 2793889 | 6.72 |
| 2809092 | 6.61 |
| 2817839 | 6.61 |
| 2829495 | 6.72 |
| 2836741 | 6.72 |
| 2847108 | 6.72 |
| 2859678 | 6.83 |
| 2871254 | 6.72 |
| 2906027 | 6.72 |
| 2920630 | 6.72 |
| 2923222 | 6.72 |
| 2935773 | 6.83 |
| 2936709 | 6.72 |
| 2957939 | 6.72 |
| 2969699 | 6.72 |
| 2979521 | 6.72 |
| 3009053 | 6.72 |
| 3026484 | 6.72 |
| 3040618 | 6.72 |
| 3061148 | 6.83 |
| 3066612 | 6.72 |
| 3074267 | 6.83 |
| 3086535 | 6.72 |
| 3103036 | 6.61 |
| 3110896 | 6.72 |
| 3135364 | 6.72 |
| 3141667 | 6.72 |
| 3152708 | 6.72 |
| 3163106 | 6.83 |
| 3166951 | 6.72 |
| 3171402 | 6.72 |
| 3183698 | 6.72 |
| 3191909 | 6.72 |
| 3193035 | 6.72 |
| 3194903 | 6.83 |
| 3203526 | 6.72 |
| 3206892 | 6.72 |
| 3245079 | 6.72 |
| 3245479 | 6.72 |
| 3253923 | 6.72 |
| 3278761 | 5.22 |
| 3286003 | 6.72 |
| 3306551 | 6.72 |
| 3334334 | 6.72 |
| 3338134 | 6.83 |
| 3369629 | 6.72 |
| 3373457 | 6.72 |
| 3384561 | 6.83 |
| 3385060 | 6.83 |
| 3406586 | 6.83 |
| 3406929 | 6.72 |
| 3410770 | 6.72 |
| 3416645 | 6.83 |
| 3442627 | 6.72 |
| 3454991 | 6.72 |
| 3477360 | 6.83 |
| 3484243 | 0 |
| 3507547 | 6.83 |
| 3528539 | 6.83 |
| 3555717 | 6.83 |
| 3567117 | 6.72 |
| 3568665 | 6.72 |
| 3573388 | 6.72 |
| 3577795 | 6.95 |
| 3584122 | 6.95 |
| 3592522 | 6.95 |
| 3652332 | 6.95 |
| 3669097 | 6.95 |
| 3712621 | 6.72 |
| 3722869 | 6.38 |
| 3727934 | 6.95 |
| 3733337 | 7.05 |
| 3752764 | 6.95 |
| 3772749 | 6.95 |
| 3778919 | 6.95 |
| 3779313 | 4.99 |
| 3786271 | 7.17 |
| 3793874 | 4.99 |
| 3808145 | 7.05 |
| 3808454 | 7.05 |
| 3813546 | 7.05 |
| 3822449 | 6.95 |
| 3825662 | 7.05 |
| 3834696 | 7.05 |
| 3847217 | 7.05 |
| 3854431 | 7.05 |
| 3863023 | 7.17 |
| 3874675 | 7.17 |
| 3881934 | 6.95 |
| 3940373 | 7.05 |
| 3940804 | 7.17 |
| 3943181 | 7.05 |
| 3954263 | 6.95 |
| 3956852 | 7.05 |
| 3963305 | 6.95 |
| 3977733 | 7.17 |
| 3997469 | 7.05 |
| 4003329 | 4.99 |
| 4024348 | 6.95 |
| 4031122 | 7.05 |
| 4031685 | 7.05 |
| 4044474 | 7.05 |
| 4063904 | 7.05 |
| 4070206 | 7.05 |
| 4083813 | 7.05 |
| 4084889 | 7.05 |
| 4087978 | 4.99 |
| 4114036 | 7.05 |
| 4133950 | 5.91 |
| 4144055 | 7.05 |
| 4160583 | 7.05 |
| 4170043 | 7.05 |
| 4180225 | 7.05 |
| 4198841 | 6.95 |
| 4218955 | 6.72 |
| 4223676 | 6.95 |
| 4251319 | 6.72 |
| 4257754 | 7.05 |
| 4267309 | 7.05 |
| 4330516 | 7.05 |
| 4334276 | 6.83 |
| 4357543 | 7.17 |
| 4383758 | 7.05 |
| 4404140 | 7.28 |
| 4415663 | 6.72 |
| 4416593 | 7.83 |
| 4427327 | 4.99 |
| 4438170 | 2.19 |
| 4440437 | 7.05 |
| 4454272 | 7.83 |
| 4456209 | 7.83 |
| 4471977 | 7.83 |
| 4486245 | 3.23 |
| 4489508 | 7.83 |
| 4494635 | 7.05 |
| 4531709 | 7.05 |
| 4536203 | 7.83 |
| 4555432 | 7.83 |
| 4575222 | 7.05 |
| 4593024 | 7.72 |
| 4607811 | 7.05 |
| 4615526 | 7.83 |
| 4635890 | 8.37 |
| 4640942 | 5.91 |
| 4641157 | 8.26 |
| 4644064 | 7.05 |

LG3

|  | genetic_pos |
| --- | --- |
| 422467 | 5.4 |
| 471868 | 5.4 |
| 493349 | 5.4 |
| 505615 | 5.4 |
| 511848 | 5.4 |
| 512810 | 5.4 |
| 513603 | 5.4 |
| 513894 | 5.4 |
| 515909 | 5.4 |
| 516860 | 5.4 |
| 517054 | 5.4 |
| 517514 | 5.4 |
| 517918 | 5.4 |
| 526247 | 5.4 |
| 528491 | 5.4 |
| 528771 | 4.92 |
| 529028 | 5.4 |
| 530269 | 5.4 |
| 530505 | 4.92 |
| 531669 | 4.92 |
| 532240 | 4.92 |
| 578765 | 5.4 |
| 590453 | 5.4 |
| 2050159 | 6.56 |
| 2064962 | 4.92 |
| 2085740 | 5.4 |
| 2095437 | 5.21 |
| 2096809 | 5.4 |
| 2108318 | 5.4 |
| 2115683 | 4.82 |
| 2139932 | 3.73 |
| 2150136 | 5.4 |
| 2185568 | 5.4 |
| 2193626 | 4.92 |
| 2200856 | 5.4 |
| 2203026 | 5.21 |
| 2466309 | 5.4 |
| 2484408 | 4.82 |
| 2704985 | 4.63 |
| 2733520 | 5.4 |
| 2736772 | 5.4 |
| 2745009 | 5.4 |
| 2778112 | 4.82 |
| 2792044 | 5.01 |
| 2793827 | 6.56 |
| 2798425 | 5.01 |
| 2807032 | 4.92 |
| 2813396 | 5.4 |
| 2815897 | 6.56 |
| 2837778 | 5.4 |
| 2847781 | 5.21 |
| 2862332 | 5.4 |
| 3872597 | 5.4 |
| 3880699 | 6.56 |
| 3926493 | 5.01 |
| 3927100 | 5.4 |
| 3927802 | 4.82 |
| 3931133 | 5.4 |
| 3948458 | 5.4 |
| 3982208 | 6.56 |
| 4043390 | 6.56 |
| 4051933 | 5.4 |
| 4072003 | 6.56 |
| 4114920 | 4.82 |
| 4120424 | 5.79 |
| 4697685 | 5.89 |
| 4704485 | 4.82 |
| 4723962 | 5.6 |
| 4728702 | 5.4 |
| 4772455 | 6.56 |
| 4777455 | 5.21 |
| 4787489 | 5.4 |
| 4796533 | 5.21 |
| 4807388 | 5.6 |
| 4808140 | 5.99 |
| 4826861 | 5.7 |
| 4834273 | 5.4 |
| 4841117 | 4.82 |
| 4865229 | 5.6 |
| 4880335 | 5.6 |
| 4882175 | 5.21 |
| 4887226 | 5.4 |
| 4891388 | 5.6 |
| 4919345 | 5.6 |
| 4922057 | 5.4 |
| 4958330 | 6.56 |
| 4961332 | 5.6 |
| 4990192 | 5.6 |
| 5005374 | 5.6 |
| 5005743 | 5.89 |
| 5010272 | 5.6 |
| 5019540 | 5.6 |
| 5039334 | 5.6 |
| 5072719 | 5.21 |
| 5073048 | 4.82 |
| 5074846 | 5.4 |
| 5077329 | 6.56 |
| 5081706 | 5.6 |
| 5085193 | 5.99 |
| 5089796 | 5.79 |
| 5094586 | 6.56 |
| 5108721 | 6.56 |
| 5145071 | 5.6 |
| 5146914 | 4.92 |
| 5150473 | 5.4 |
| 5153342 | 5.6 |
| 5162942 | 4.82 |
| 5172193 | 5.79 |
| 5182288 | 6.94 |
| 5200641 | 5.6 |
| 5205123 | 6.94 |
| 5213633 | 6.09 |
| 5225918 | 5.79 |
| 5229753 | 5.79 |
| 5233744 | 5.79 |
| 5236347 | 6.56 |
| 5239585 | 6.28 |
| 5244749 | 5.89 |
| 5253897 | 6.28 |
| 5261190 | 8.17 |
| 5278255 | 6.75 |
| 5281711 | 6.56 |
| 5290927 | 4.82 |
| 5296034 | 5.6 |
| 5298164 | 6.56 |
| 5305361 | 5.79 |
| 7888597 | 5.89 |
| 7891436 | 3.73 |
| 7892864 | 5.21 |
| 7894204 | 4.23 |
| 7894396 | 4.43 |
| 7895060 | 6.56 |
| 7912177 | 5.79 |
| 7914497 | 6.56 |
| 7915833 | 6.56 |
| 7923742 | 6.28 |
| 7953513 | 5.79 |
| 7958890 | 6.56 |
| 7963327 | 6.56 |
| 7973943 | 6.56 |
| 8609481 | 6.56 |
| 8671594 | 5.79 |
| 8690612 | 3.35 |
| 8700818 | 6.18 |
| 8718204 | 6.28 |
| 8729420 | 6.56 |
| 8778020 | 6.56 |
| 8789620 | 6.28 |
| 8797255 | 4.14 |
| 8807097 | 4.23 |
| 8819731 | 4.82 |
| 8825875 | 2.97 |
| 8828271 | 6.18 |
| 8837992 | 5.89 |
| 8838318 | 5.79 |
| 8845845 | 5.89 |
| 8847933 | 5.79 |
| 8883257 | 6.56 |
| 8884236 | 6.56 |
| 8919515 | 7.04 |
| 8937904 | 5.6 |
| 8944867 | 4.23 |
| 8967944 | 6.56 |
| 8986886 | 4.82 |
| 8996197 | 4.63 |
| 9027406 | 5.79 |
| 9058525 | 6.56 |
| 9065204 | 5.79 |
| 9078586 | 5.79 |
| 10138516 | 4.14 |
| 10185579 | 4.43 |
| 10193381 | 6.28 |
| 10223580 | 4.23 |
| 10306122 | 5.89 |
| 10325607 | 6.28 |
| 10336295 | 4.53 |
| 10346401 | 0 |
| 10352938 | 7.04 |
| 10358600 | 6.56 |
| 10374649 | 5.89 |
| 10383196 | 6.28 |
| 10400000 | 5.89 |
| 10404751 | 5.79 |
| 10415221 | 5.6 |
| 10425337 | 6.28 |
| 10435077 | 6.09 |
| 10442236 | 7.04 |
| 10455431 | 7.04 |
| 10464378 | 5.89 |
| 10472886 | 6.56 |
| 10493101 | 7.13 |
| 10499165 | 7.13 |
| 10501306 | 7.13 |
| 10501960 | 7.89 |
| 10554560 | 6.94 |
| 10561047 | 5.79 |
| 10595670 | 6.75 |
| 10624883 | 6.09 |
| 10626000 | 6.09 |
| 10674707 | 6.28 |

LG4

|  | genetic_pos |
| --- | --- |
| 523966 | 4.92 |
| 535750 | 6.47 |
| 559574 | 6.47 |
| 611572 | 6.47 |
| 611978 | 6.38 |
| 614624 | 6.47 |
| 619199 | 5.92 |
| 656917 | 6.47 |
| 661561 | 6.38 |
| 664732 | 6.47 |
| 678686 | 6.47 |
| 685309 | 5.74 |
| 706596 | 6.47 |
| 763606 | 2.63 |
| 774632 | 6.29 |
| 780962 | 6.47 |
| 786593 | 4.92 |
| 823460 | 6.29 |
| 999044 | 5.92 |
| 1000296 | 7.11 |
| 1000605 | 6.47 |
| 1002124 | 5.74 |
| 1002274 | 6.47 |
| 1004307 | 5.37 |
| 1005587 | 6.47 |
| 1011896 | 6.47 |
| 1018343 | 6.29 |
| 1034593 | 6.47 |
| 1044803 | 2.63 |
| 1060572 | 6.47 |
| 1064662 | 6.57 |
| 1071142 | 3.27 |
| 1075508 | 6.38 |
| 1076063 | 7.11 |
| 1077283 | 7.11 |
| 1086528 | 6.38 |
| 1088027 | 4.92 |
| 1093983 | 7.2 |
| 1102434 | 6.38 |
| 1108504 | 6.47 |
| 1237961 | 4.92 |
| 1242132 | 6.38 |
| 1256256 | 6.47 |
| 1280148 | 6.38 |
| 1281268 | 6.47 |
| 1290601 | 6.47 |
| 1291251 | 6.29 |
| 1302900 | 4.36 |
| 1339322 | 6.29 |
| 1357212 | 6.47 |
| 1400535 | 6.29 |
| 1785996 | 7.11 |
| 1790892 | 6.47 |
| 1820002 | 6.38 |
| 1869820 | 6.75 |
| 1877398 | 6.47 |
| 1882251 | 7.47 |
| 1883307 | 6.47 |
| 2172634 | 6.47 |
| 2202076 | 3.17 |
| 2206551 | 5.74 |
| 2209812 | 7.2 |
| 2434317 | 6.47 |
| 2449155 | 6.47 |
| 2451178 | 6.29 |
| 2460700 | 6.29 |
| 2472475 | 7.11 |
| 2473425 | 7.11 |
| 2477815 | 6.29 |
| 2478730 | 6.29 |
| 2481405 | 6.47 |
| 2482648 | 5.92 |
| 2487634 | 7.11 |
| 2489256 | 6.47 |
| 2489754 | 6.47 |
| 2491768 | 6.38 |
| 2498677 | 0.65 |
| 2500230 | 4.16 |
| 2519809 | 7.11 |
| 2521241 | 6.47 |
| 2523293 | 5.74 |
| 2526887 | 6.47 |
| 2527959 | 4.92 |
| 2539936 | 7.11 |
| 2596831 | 5.37 |
| 2628869 | 5.74 |
| 2644057 | 6.75 |
| 2646032 | 5.74 |
| 2648983 | 6.38 |
| 2649860 | 6.11 |
| 2653961 | 6.47 |
| 2657792 | 6.11 |
| 2658859 | 6.29 |
| 2667772 | 6.47 |
| 2673643 | 7.2 |
| 2677405 | 6.47 |
| 2679068 | 7.2 |
| 2688248 | 7.11 |
| 2689042 | 6.38 |
| 2689917 | 6.47 |
| 2692430 | 6.38 |
| 2709770 | 4.92 |
| 2715814 | 6.11 |
| 2716478 | 6.47 |
| 2760945 | 5.74 |
| 2764266 | 6.11 |
| 2787173 | 7.11 |
| 2788607 | 6.11 |
| 2794133 | 7.2 |
| 2796277 | 7.2 |
| 2800489 | 7.11 |
| 2807369 | 6.29 |
| 2819830 | 5.92 |
| 2860045 | 6.75 |
| 2875487 | 5.37 |
| 2877533 | 7.11 |
| 2888386 | 6.29 |
| 2891531 | 7.11 |
| 2893213 | 7.11 |
| 2895834 | 6.47 |
| 2902917 | 6.47 |
| 3033838 | 7.11 |
| 3121050 | 7.2 |
| 3271413 | 8.18 |
| 3476366 | 5.39 |
| 3504331 | 61.96 |
| 3566202 | 61.86 |
| 3599039 | 61.96 |
| 3657321 | 61.96 |
| 3685563 | 64.34 |
| 3685665 | 61.96 |
| 3697050 | 61.96 |
| 3708996 | 61.96 |
| 3724117 | 62.64 |
| 3767847 | 61.17 |
| 3771308 | 61.96 |
| 3775569 | 61.96 |
| 4233775 | 5.74 |
| 4313024 | 4.92 |
| 4314405 | 6.75 |
| 4327145 | 6.47 |
| 4344069 | 7.11 |
| 4527369 | 7.11 |
| 4658257 | 6.47 |
| 4674915 | 6.47 |
| 4684281 | 6.38 |
| 4706458 | 6.47 |
| 4714237 | 7.11 |
| 4780724 | 7.2 |
| 4791396 | 5.74 |
| 4803520 | 4.92 |
| 4851061 | 6.47 |
| 4852430 | 7.11 |
| 4861813 | 5.92 |
| 4871918 | 6.47 |
| 4880359 | 6.47 |
| 4901587 | 6.47 |
| 4918284 | 6.47 |
| 4919863 | 5.74 |
| 4934741 | 6.47 |
| 4940651 | 6.47 |
| 4962741 | 4.27 |
| 5035595 | 6.47 |
| 5041165 | 6.75 |
| 5055929 | 5.37 |
| 5061830 | 6.57 |
| 5112705 | 6.47 |
| 5124244 | 6.47 |
| 5129900 | 4.27 |
| 5139425 | 6.47 |
| 5142387 | 4.92 |
| 5150776 | 5.74 |
| 5158655 | 6.47 |
| 5180796 | 7.11 |
| 5191259 | 7.29 |
| 5219376 | 6.75 |
| 5234826 | 7.11 |
| 5247155 | 6.57 |
| 5271294 | 5.74 |
| 5281375 | 6.57 |
| 5289195 | 6.75 |
| 5331703 | 6.57 |
| 5370776 | 5.74 |
| 5372148 | 6.57 |
| 5382319 | 6.75 |
| 5385058 | 6.57 |
| 5415287 | 5.37 |
| 5419780 | 6.57 |
| 5420482 | 6.57 |
| 5420745 | 6.57 |
| 5428742 | 6.57 |
| 5458266 | 6.57 |
| 5459142 | 7.2 |
| 5496622 | 7.11 |
| 5507223 | 5.92 |
| 5768371 | 6.47 |
| 5777241 | 6.47 |
| 5783765 | 4.82 |
| 5788498 | 6.57 |
| 5818757 | 7.11 |

LG5

|  | genetic_pos |
| --- | --- |
| 233468 | 62.04 |
| 243584 | 4.93 |
| 245401 | 5.39 |
| 254054 | 6.14 |
| 277925 | 4.84 |
| 280325 | 5.95 |
| 292936 | 5.3 |
| 294558 | 5.3 |
| 294842 | 5.39 |
| 295997 | 5.3 |
| 297472 | 5.3 |
| 338151 | 5.02 |
| 339010 | 5.3 |
| 339589 | 5.3 |
| 339806 | 5.3 |
| 346615 | 5.3 |
| 347104 | 5.3 |
| 350918 | 5.3 |
| 353163 | 5.3 |
| 353526 | 5.3 |
| 355003 | 5.3 |
| 366154 | 5.3 |
| 367945 | 5.39 |
| 371037 | 5.39 |
| 392231 | 5.3 |
| 404818 | 5.3 |
| 420115 | 5.58 |
| 486377 | 5.02 |
| 513482 | 5.3 |
| 530383 | 5.3 |
| 540762 | 5.95 |
| 557990 | 5.58 |
| 590609 | 5.02 |
| 656544 | 5.02 |
| 668844 | 5.3 |
| 683382 | 5.3 |
| 692779 | 5.3 |
| 702086 | 5.02 |
| 732893 | 5.95 |
| 737205 | 5.3 |
| 748524 | 5.3 |
| 760639 | 5.02 |
| 784807 | 5.02 |
| 820147 | 5.86 |
| 829195 | 5.02 |
| 838382 | 5.86 |
| 873068 | 5.3 |
| 889425 | 5.3 |
| 935320 | 5.3 |
| 970799 | 5.86 |
| 1135466 | 5.02 |
| 1158360 | 5.02 |
| 1335521 | 5.3 |
| 1349017 | 5.02 |
| 1405501 | 5.3 |
| 1427797 | 5.02 |
| 1460206 | 5.3 |
| 1477263 | 5.95 |
| 1489620 | 5.02 |
| 1548609 | 5.3 |
| 1549928 | 5.86 |
| 1557163 | 5.3 |
| 1566468 | 4.2 |
| 1585577 | 5.3 |
| 1687495 | 5.3 |
| 1697406 | 5.3 |
| 1717274 | 5.3 |
| 1725620 | 4.65 |
| 1731220 | 5.86 |
| 1737734 | 5.02 |
| 1989989 | 5.86 |
| 1995667 | 5.02 |
| 2005687 | 5.02 |
| 2013720 | 6.14 |
| 2016793 | 5.3 |
| 2029181 | 5.02 |
| 2058804 | 5.76 |
| 2069175 | 5.95 |
| 2078174 | 5.3 |
| 2093351 | 5.02 |
| 2124900 | 5.02 |
| 2134240 | 5.3 |
| 2139907 | 5.3 |
| 2140767 | 5.3 |
| 2150972 | 5.3 |
| 2183360 | 5.95 |
| 2197272 | 5.86 |
| 2216584 | 5.3 |
| 2230928 | 5.3 |
| 2239377 | 5.02 |
| 2248074 | 5.3 |
| 2279372 | 5.3 |
| 2282545 | 5.3 |
| 2287191 | 5.58 |
| 2387324 | 5.39 |
| 2394456 | 5.3 |
| 2402517 | 5.02 |
| 2427625 | 5.3 |
| 2432048 | 5.3 |
| 2433233 | 5.86 |
| 2448201 | 5.39 |
| 2463214 | 5.86 |
| 2469769 | 5.76 |
| 2470802 | 5.02 |
| 2473579 | 5.39 |
| 2479842 | 5.3 |
| 2493377 | 5.3 |
| 2502386 | 5.02 |
| 2513571 | 5.95 |
| 2569753 | 5.02 |
| 2588832 | 5.39 |
| 2599220 | 6.14 |
| 2609137 | 5.02 |
| 2621235 | 5.95 |
| 2621446 | 5.39 |
| 2626229 | 5.95 |
| 2647734 | 5.02 |
| 2670962 | 5.02 |
| 2673632 | 5.3 |
| 2677496 | 5.02 |
| 2682970 | 5.02 |
| 2687613 | 5.3 |
| 2690363 | 5.58 |
| 2693778 | 5.02 |
| 2701998 | 5.02 |
| 2721722 | 5.3 |
| 2732792 | 5.86 |
| 2734699 | 5.3 |
| 2736317 | 5.39 |
| 2750898 | 5.76 |
| 2755519 | 5.95 |
| 2790691 | 5.3 |
| 2793391 | 5.86 |
| 2800251 | 5.39 |
| 2808800 | 5.3 |
| 2819446 | 5.76 |
| 2827419 | 4.74 |
| 2833099 | 5.95 |
| 2844092 | 5.3 |
| 2864127 | 5.3 |
| 2871358 | 6.14 |
| 2900029 | 5.02 |
| 2939412 | 5.95 |
| 2954966 | 5.86 |
| 2975530 | 3.31 |
| 3845177 | 5.02 |
| 3879993 | 5.86 |
| 3895370 | 5.02 |
| 3923374 | 5.3 |
| 3934971 | 5.02 |
| 3939813 | 5.3 |
| 3968927 | 5.3 |
| 3978295 | 6.23 |
| 3985315 | 5.21 |
| 3992673 | 5.3 |
| 3995916 | 6.32 |
| 4012826 | 5.86 |
| 4031677 | 5.39 |
| 4032843 | 5.02 |
| 4033399 | 5.39 |
| 4188632 | 5.02 |
| 4613807 | 5.3 |
| 4621382 | 5.02 |
| 4626884 | 4.65 |
| 4627560 | 5.02 |
| 4632764 | 5.39 |
| 4681817 | 5.86 |
| 4767960 | 5.02 |
| 4783095 | 5.3 |
| 4788846 | 6.23 |
| 4822486 | 5.21 |
| 4823138 | 5.86 |
| 4835883 | 5.02 |
| 4845976 | 5.76 |
| 4851897 | 5.86 |
| 4853438 | 5.76 |
| 4860637 | 5.58 |
| 4866110 | 5.76 |
| 4882859 | 6.23 |
| 4886243 | 5.3 |
| 4888545 | 5.02 |
| 4902565 | 5.02 |
| 4911683 | 5.86 |
| 4917067 | 5.49 |
| 4917656 | 5.76 |
| 4919639 | 5.02 |
| 4926268 | 4.07 |
| 4928537 | 6.41 |
| 4978506 | 5.02 |
| 4998992 | 5.02 |
| 5021566 | 5.12 |
| 5030928 | 5.3 |
| 5035359 | 4.65 |
| 5048231 | 5.39 |
| 5051575 | 5.58 |
| 5059764 | 6.23 |
| 5071149 | 5.58 |
| 5079565 | 5.02 |
| 5090998 | 5.02 |
| 5103870 | 5.3 |

LG6

|  | genetic_pos |
| --- | --- |
| 209637 | 5.33 |
| 249055 | 5.22 |
| 265469 | 5.33 |
| 302935 | 5.33 |
| 316026 | 5.33 |
| 359119 | 4.66 |
| 367507 | 5.33 |
| 403869 | 4.44 |
| 409532 | 5.33 |
| 454663 | 5.33 |
| 595622 | 1.52 |
| 612013 | 5.33 |
| 626147 | 5.33 |
| 636644 | 5.33 |
| 656436 | 3.53 |
| 698297 | 2.37 |
| 729571 | 5.33 |
| 746178 | 4.66 |
| 767313 | 4.66 |
| 806772 | 4.44 |
| 833882 | 5.33 |
| 874908 | 5.33 |
| 904418 | 5.22 |
| 905578 | 5.22 |
| 920168 | 5.22 |
| 925775 | 5.22 |
| 943672 | 4.66 |
| 953858 | 5.22 |
| 993872 | 5.33 |
| 1013241 | 5.11 |
| 1040326 | 5.33 |
| 1062716 | 5.33 |
| 1063042 | 5.22 |
| 1064602 | 5.22 |
| 1070831 | 4.66 |
| 1076713 | 4.44 |
| 1079065 | 5.54 |
| 1079997 | 5.33 |
| 1094004 | 5.11 |
| 1100425 | 4.44 |
| 1111194 | 4.44 |
| 1123656 | 5.33 |
| 1152815 | 4.44 |
| 1187969 | 5.33 |
| 1197065 | 4.66 |
| 1210581 | 4.66 |
| 1213575 | 4.66 |
| 1220543 | 4.66 |
| 1221717 | 4.44 |
| 1229693 | 5.33 |
| 1229816 | 5.33 |
| 1285020 | 4.44 |
| 1293105 | 4.66 |
| 1331305 | 2.81 |
| 1360954 | 2.81 |
| 1365257 | 4.44 |
| 1409888 | 4.44 |
| 1415133 | 5.33 |
| 1417274 | 4.66 |
| 1448672 | 4.44 |
| 1456236 | 4.66 |
| 1466638 | 4.44 |
| 1478160 | 5.11 |
| 1508311 | 4.66 |
| 1517880 | 5.33 |
| 1548205 | 5.22 |
| 1555487 | 4.66 |
| 1562896 | 5.11 |
| 1583332 | 4.44 |
| 1593571 | 4.66 |
| 1596904 | 4.66 |
| 1605842 | 4.66 |
| 1624214 | 4.66 |
| 1626517 | 4.66 |
| 1779896 | 6.18 |
| 1817211 | 5.22 |
| 2241671 | 4.66 |
| 2242263 | 4.66 |
| 2244315 | 4.66 |
| 2252162 | 5.22 |
| 2255301 | 4.44 |
| 2255860 | 4.22 |
| 2267200 | 4.22 |
| 2269775 | 5.33 |
| 2271716 | 4.22 |
| 2277709 | 5.33 |
| 2283080 | 4.22 |
| 2286540 | 5.22 |
| 2295596 | 4.44 |
| 2301497 | 4.66 |
| 2313863 | 4.22 |
| 2314143 | 5.22 |
| 2314606 | 5.33 |
| 2319868 | 4.66 |
| 2323557 | 5.33 |
| 2327281 | 5.33 |
| 2334967 | 4.44 |
| 2339081 | 5.33 |
| 2350044 | 5.33 |
| 2376475 | 5.33 |
| 2381343 | 5.22 |
| 2387791 | 4.66 |
| 2391800 | 5.22 |
| 2395822 | 4.44 |
| 2397008 | 4.66 |
| 2403700 | 5.33 |
| 2408625 | 5.22 |
| 2409732 | 5.33 |
| 2425973 | 4.66 |
| 2436262 | 5.33 |
| 2449528 | 5.22 |
| 2457328 | 5.22 |
| 2470292 | 5.33 |
| 2479000 | 4.44 |
| 2497428 | 4.44 |
| 2533277 | 5.22 |
| 2541446 | 4.44 |
| 2553373 | 5.33 |
| 2563078 | 5.33 |
| 2565799 | 4.66 |
| 2574117 | 4.66 |
| 2576951 | 4.44 |
| 2592443 | 4.66 |
| 2620229 | 4.66 |
| 2622521 | 5.22 |
| 2636236 | 4.22 |
| 2666874 | 4.66 |
| 2679639 | 5.33 |
| 2686298 | 5.22 |
| 2706272 | 4.66 |
| 2727376 | 3.99 |
| 2734881 | 5.33 |
| 2756395 | 4.66 |
| 2777132 | 4.66 |
| 2785066 | 2.81 |
| 2804653 | 5.33 |
| 2838648 | 5.22 |
| 2858099 | 4.44 |
| 2864632 | 4.66 |
| 2867443 | 5.54 |
| 2874957 | 4.66 |
| 2877789 | 5.44 |
| 2896190 | 4.66 |
| 2969077 | 5.54 |
| 2980442 | 5.54 |
| 3000349 | 5.44 |
| 3022301 | 4.44 |
| 3023132 | 5.44 |
| 3034390 | 5.44 |
| 3066459 | 4.66 |
| 3087581 | 5.54 |
| 3094478 | 5.54 |
| 3103759 | 5.44 |
| 3115956 | 5.54 |
| 3132745 | 5.54 |
| 3154268 | 5.44 |
| 3184125 | 4.33 |
| 3392082 | 5.54 |
| 3402107 | 5.76 |
| 3411256 | 5.44 |
| 3414557 | 5.76 |
| 3429794 | 7.19 |
| 3438157 | 5.87 |
| 3441425 | 2.22 |
| 3441950 | 5.87 |
| 3452184 | 5.87 |
| 3452885 | 5.97 |
| 3460362 | 5.76 |
| 3462813 | 5.87 |
| 3481425 | 4.22 |
| 3484567 | 5.76 |
| 3490922 | 5.76 |
| 3501875 | 1.56 |
| 3506349 | 7.19 |
| 3533321 | 7.19 |
| 3542878 | 6.08 |
| 3559520 | 5.76 |
| 3573808 | 5.44 |
| 3576909 | 7.19 |
| 3580746 | 7.19 |
| 3588602 | 7.19 |
| 3589615 | 4.66 |
| 3609283 | 7.19 |
| 3621440 | 5.76 |
| 3630967 | 5.76 |
| 3646421 | 5.44 |
| 3654161 | 4.66 |
| 3675650 | 5.44 |
| 3684852 | 5.76 |
| 3692834 | 4.66 |
| 3704987 | 5.76 |
| 3715298 | 4.66 |
| 3743805 | 4.66 |
| 3773938 | 5.76 |
| 3779217 | 4.44 |
| 3779805 | 4.66 |
| 3782931 | 4.66 |
| 3849638 | 7.19 |
| 3856991 | 7.19 |
| 3868869 | 5.76 |

LG7

|  | genetic_pos |
| --- | --- |
| 32783 | 2.3 |
| 58725 | 2.06 |
| 77847 | 1.52 |
| 92775 | 1.52 |
| 118499 | 1.52 |
| 334142 | 2.55 |
| 513196 | 1.52 |
| 563554 | 2.55 |
| 604351 | 2.3 |
| 681999 | 2.55 |
| 691240 | 2.3 |
| 705031 | 1.52 |
| 709623 | 2.06 |
| 848031 | 1.52 |
| 862536 | 2.55 |
| 868303 | 2.3 |
| 917484 | 2.55 |
| 920667 | 2.55 |
| 930373 | 2.55 |
| 934695 | 1.52 |
| 937884 | 2.55 |
| 949187 | 2.55 |
| 960811 | 2.55 |
| 967258 | 2.55 |
| 999959 | 2.06 |
| 1009587 | 2.55 |
| 1015037 | 2.3 |
| 1028702 | 2.55 |
| 1049168 | 2.55 |
| 1077622 | 2.55 |
| 1080521 | 2.55 |
| 1115119 | 2.55 |
| 1120858 | 2.8 |
| 1171957 | 2.06 |
| 1178238 | 2.55 |
| 1193099 | 2.55 |
| 1198308 | 1.52 |
| 1199952 | 2.55 |
| 1202955 | 1.52 |
| 1253694 | 2.55 |
| 1285451 | 2.3 |
| 1296412 | 2.3 |
| 1307413 | 1.52 |
| 1321819 | 1.52 |
| 1327039 | 1.52 |
| 1340398 | 2.55 |
| 1347433 | 2.55 |
| 1390373 | 2.55 |
| 1536243 | 1.52 |
| 1538205 | 2.55 |
| 1574038 | 2.55 |
| 1643762 | 2.3 |
| 1665785 | 2.55 |
| 1775710 | 1.52 |
| 1786766 | 1.52 |
| 1818391 | 1.52 |
| 1828468 | 2.55 |
| 1849650 | 2.55 |
| 1863579 | 1.52 |
| 1875395 | 1.52 |
| 1884006 | 1.52 |
| 1903829 | 2.55 |
| 1932037 | 2.3 |
| 1951141 | 2.55 |
| 1990277 | 2.55 |
| 2001398 | 2.55 |
| 2054359 | 2.55 |
| 2079640 | 2.55 |
| 2081080 | 1.52 |
| 2089065 | 2.55 |
| 2143644 | 2.55 |
| 2147129 | 2.55 |
| 2168522 | 1.52 |
| 2185842 | 1.52 |
| 2191415 | 1.52 |
| 2196428 | 2.55 |
| 2198092 | 2.3 |
| 2205137 | 1.52 |
| 2337879 | 2.55 |
| 2367713 | 2.55 |
| 2379940 | 2.55 |
| 2438883 | 2.3 |
| 2446280 | 2.55 |
| 2451271 | 2.06 |
| 2610833 | 2.55 |
| 2624183 | 2.55 |
| 2668240 | 2.55 |
| 2678040 | 1.52 |
| 2698914 | 1.52 |
| 2708896 | 2.55 |
| 2719763 | 2.55 |
| 2730059 | 2.06 |
| 2744573 | 1.52 |
| 2752194 | 2.55 |
| 2805045 | 2.55 |
| 2812923 | 2.06 |
| 2850643 | 2.55 |
| 2851988 | 1.52 |
| 2856734 | 1.52 |
| 2857941 | 2.55 |
| 2895508 | 2.55 |
| 2905060 | 2.3 |
| 2920676 | 2.55 |
| 2922896 | 2.55 |
| 2929358 | 2.55 |
| 2931160 | 2.55 |
| 2933720 | 2.55 |
| 2955920 | 2.55 |
| 2974230 | 2.3 |
| 3024567 | 2.55 |
| 3035970 | 2.55 |
| 3040121 | 2.99 |
| 3046909 | 2.99 |
| 3054411 | 2.8 |
| 3073934 | 2.99 |
| 3081889 | 2.99 |
| 3145174 | 2.55 |
| 3156603 | 2.99 |
| 3162361 | 2.99 |
| 3175199 | 2.99 |
| 3347814 | 2.99 |
| 3385177 | 2.99 |
| 3412487 | 2.99 |
| 3434730 | 3.24 |
| 3440605 | 2.99 |
| 3451749 | 2.99 |
| 3484862 | 2.99 |
| 3569075 | 2.99 |
| 3581195 | 2.99 |
| 3598183 | 2.55 |
| 3608622 | 2.99 |
| 3667328 | 3.24 |
| 3793783 | 3.24 |
| 3802749 | 2.99 |
| 3862175 | 2.99 |
| 3873597 | 2.99 |
| 3891296 | 2.8 |
| 3916417 | 2.99 |
| 3926180 | 3.24 |
| 3938590 | 2.8 |
| 3969290 | 2.99 |
| 3993803 | 2.99 |
| 4001199 | 2.99 |
| 4004641 | 3.24 |
| 4014447 | 2.99 |
| 4186946 | 2.55 |
| 4189898 | 3.24 |
| 4196068 | 3.86 |
| 4257130 | 2.99 |
| 4258613 | 2.99 |
| 4308599 | 3.24 |
| 4321921 | 2.99 |
| 4331706 | 2.99 |
| 4338289 | 3.24 |
| 4396043 | 2.99 |
| 4404734 | 2.99 |
| 4414605 | 2.99 |
| 4424587 | 2.99 |
| 4436899 | 3.48 |
| 4444513 | 3.24 |
| 4472579 | 4.03 |
| 4491088 | 2.99 |
| 4492330 | 3.24 |
| 4584932 | 2.99 |
| 4604357 | 4.03 |
| 4604884 | 2.99 |
| 4615051 | 2.99 |
| 4646121 | 3.86 |
| 4704223 | 3.24 |
| 4719945 | 3.67 |
| 4733970 | 3.67 |
| 4769183 | 3.24 |
| 4827048 | 3.24 |
| 4833511 | 2.99 |
| 4852413 | 2.99 |
| 4913851 | 3.24 |
| 4922280 | 3.24 |
| 4932590 | 4.03 |
| 4945550 | 3.24 |
| 4982634 | 2.99 |
| 5027042 | 2.99 |
| 5047840 | 2.99 |
| 5283150 | 5.99 |
| 5294131 | 3.24 |
| 5313786 | 3.24 |
| 5333199 | 3.24 |
| 5341285 | 2.99 |
| 5372577 | 3.48 |
| 5383200 | 2.55 |
| 5459004 | 2.99 |
| 5504640 | 2.99 |
| 5518737 | 3.24 |
| 5529559 | 3.24 |
| 5530834 | 3.24 |
| 5535764 | 2.99 |
| 5560422 | 2.99 |
| 5588889 | 3.24 |
| 5613760 | 2.99 |
| 5677280 | 2.99 |
| 5696988 | 2.99 |

LG8

|  | genetic_pos |
| --- | --- |
| 96361 | 0 |
| 111455 | 6.56 |
| 114636 | 5.86 |
| 118312 | 6.26 |
| 126370 | 6.56 |
| 136389 | 5.65 |
| 151522 | 6.56 |
| 157672 | 5.44 |
| 203569 | 6.26 |
| 243948 | 5.34 |
| 275660 | 5.44 |
| 283791 | 6.56 |
| 283939 | 5.65 |
| 307714 | 6.26 |
| 317470 | 6.26 |
| 493597 | 6.56 |
| 495599 | 6.56 |
| 501271 | 6.66 |
| 506589 | 5.96 |
| 514231 | 6.56 |
| 520332 | 6.56 |
| 525157 | 6.26 |
| 533560 | 6.26 |
| 545046 | 6.56 |
| 558019 | 6.66 |
| 583373 | 6.56 |
| 586532 | 6.26 |
| 592874 | 6.86 |
| 642129 | 5.65 |
| 658041 | 5.65 |
| 681905 | 6.56 |
| 692981 | 5.55 |
| 746286 | 6.56 |
| 765122 | 6.26 |
| 771997 | 5.65 |
| 772915 | 6.26 |
| 783598 | 6.26 |
| 809556 | 6.26 |
| 859928 | 6.26 |
| 869253 | 6.56 |
| 870408 | 5.34 |
| 875304 | 6.56 |
| 878373 | 6.66 |
| 1057601 | 6.26 |
| 1063297 | 6.56 |
| 1063716 | 6.26 |
| 1072619 | 6.56 |
| 1083756 | 6.56 |
| 1096359 | 6.56 |
| 1105234 | 6.26 |
| 1120955 | 6.56 |
| 1140017 | 5.86 |
| 1147560 | 5.44 |
| 1149088 | 5.96 |
| 1167571 | 6.66 |
| 1172216 | 6.26 |
| 1180207 | 6.56 |
| 1341799 | 6.56 |
| 1343859 | 5.34 |
| 1345306 | 6.56 |
| 1348433 | 6.36 |
| 1349743 | 5.44 |
| 1350202 | 6.56 |
| 1358102 | 6.26 |
| 1372850 | 6.56 |
| 1380912 | 6.26 |
| 1751686 | 5.24 |
| 1761304 | 6.46 |
| 1770983 | 4.17 |
| 1787871 | 5.34 |
| 1806292 | 6.46 |
| 1813577 | 5.65 |
| 1860214 | 4.9 |
| 1867242 | 6.26 |
| 1924961 | 6.56 |
| 1946586 | 6.56 |
| 1963896 | 5.65 |
| 1971031 | 6.26 |
| 1973141 | 5.96 |
| 2205749 | 6.26 |
| 2216018 | 5.96 |
| 2230531 | 6.56 |
| 2241492 | 6.26 |
| 2251427 | 6.56 |
| 2256041 | 5.13 |
| 2258754 | 5.96 |
| 2266265 | 6.26 |
| 2283361 | 6.46 |
| 2292934 | 6.66 |
| 2348895 | 6.56 |
| 2385333 | 5.24 |
| 2453658 | 6.26 |
| 2456783 | 6.56 |
| 2465495 | 6.56 |
| 2475834 | 6.46 |
| 2482326 | 6.56 |
| 2486172 | 6.46 |
| 2495668 | 6.26 |
| 2500021 | 6.26 |
| 2502705 | 6.26 |
| 2507636 | 6.26 |
| 2521268 | 6.56 |
| 2530050 | 6.56 |
| 2532381 | 5.96 |
| 2547170 | 6.66 |
| 2552025 | 6.26 |
| 2572446 | 6.46 |
| 2590750 | 6.56 |
| 2599688 | 6.56 |
| 2602480 | 5.96 |
| 2612255 | 5.96 |
| 2626380 | 6.26 |
| 2659511 | 6.56 |
| 2671578 | 6.26 |
| 2690168 | 6.56 |
| 2699483 | 6.26 |
| 2709153 | 5.96 |
| 2717515 | 6.56 |
| 2742652 | 6.26 |
| 2752346 | 6.56 |
| 2758531 | 6.66 |
| 2776906 | 7.06 |
| 2779546 | 5.65 |
| 2807829 | 6.56 |
| 2826563 | 6.66 |
| 2840727 | 5.34 |
| 2861989 | 5.86 |
| 2864068 | 5.96 |
| 2903158 | 5.02 |
| 2908526 | 6.56 |
| 2932858 | 5.34 |
| 2950261 | 6.66 |
| 2957700 | 5.02 |
| 2963773 | 5.65 |
| 2982131 | 5.34 |
| 3011178 | 5.65 |
| 3022456 | 6.26 |
| 3033489 | 6.56 |
| 3043639 | 5.44 |
| 3117312 | 5.86 |
| 3120823 | 6.26 |
| 3124813 | 5.86 |
| 3143048 | 6.56 |
| 3164646 | 6.56 |
| 3166527 | 6.56 |
| 3174566 | 6.26 |
| 3197371 | 5.65 |
| 3217463 | 6.56 |
| 3252213 | 6.56 |
| 3268160 | 6.26 |
| 3280331 | 5.86 |
| 3291923 | 6.56 |
| 3298399 | 6.26 |
| 3305487 | 6.26 |
| 3312843 | 5.65 |
| 3315309 | 6.56 |
| 3319828 | 6.66 |
| 3328976 | 5.96 |
| 3336740 | 6.66 |
| 3337973 | 5.44 |
| 3349820 | 5.34 |
| 3350650 | 6.56 |
| 3351114 | 6.66 |
| 3353397 | 6.56 |
| 3356730 | 6.56 |
| 3359730 | 6.26 |
| 3419490 | 6.56 |
| 3419649 | 6.56 |
| 3419952 | 6.26 |
| 3420786 | 6.56 |
| 3446021 | 6.26 |
| 3446545 | 2.2 |
| 3450247 | 6.46 |
| 3450711 | 6.26 |
| 3451118 | 5.24 |
| 3456377 | 5.34 |
| 3456475 | 6.56 |
| 3475771 | 6.26 |
| 3491670 | 6.26 |
| 3511750 | 5.65 |
| 3533780 | 6.26 |
| 3542853 | 5.65 |
| 3554320 | 5.65 |
| 3568707 | 6.56 |
| 3575278 | 6.56 |
| 3586078 | 6.76 |
| 3591364 | 6.26 |
| 3602299 | 6.56 |
| 3612929 | 6.66 |
| 3613946 | 6.56 |
| 3623006 | 6.56 |
| 3629538 | 5.96 |
| 3665126 | 5.96 |
| 3665780 | 6.26 |
| 3666737 | 6.26 |
| 3671806 | 6.26 |
| 3677421 | 6.56 |
| 3689728 | 6.56 |
| 3692967 | 6.56 |
| 3708506 | 6.66 |

LG9

|  | genetic_pos |
| --- | --- |
| 150083 | 8.37 |
| 203246 | 8.19 |
| 295083 | 8.19 |
| 318519 | 7.81 |
| 326267 | 6.59 |
| 355948 | 8.19 |
| 364596 | 7.81 |
| 376013 | 8.19 |
| 412839 | 7.72 |
| 433999 | 8.82 |
| 435525 | 9.18 |
| 493061 | 9.18 |
| 576503 | 8.19 |
| 578684 | 8.19 |
| 581652 | 8.54 |
| 594334 | 8.54 |
| 617694 | 9.18 |
| 626704 | 9.09 |
| 634291 | 8.82 |
| 645946 | 8.91 |
| 647580 | 8.82 |
| 648984 | 9.09 |
| 649528 | 8.91 |
| 654420 | 8.91 |
| 787251 | 7.81 |
| 790304 | 7.62 |
| 797330 | 8.64 |
| 807637 | 7.12 |
| 829597 | 9.26 |
| 833093 | 7.81 |
| 871063 | 7.81 |
| 872029 | 9.18 |
| 879935 | 9.18 |
| 928794 | 8.91 |
| 1094391 | 9.18 |
| 1098042 | 8.91 |
| 1113233 | 9 |
| 1129965 | 9.18 |
| 1139370 | 8.82 |
| 1147455 | 8.82 |
| 1152715 | 8.91 |
| 1166334 | 8.82 |
| 1186112 | 8.82 |
| 1246402 | 9.18 |
| 1280722 | 9.35 |
| 1289120 | 9.26 |
| 1329048 | 9.35 |
| 1340495 | 7.81 |
| 1357290 | 9.18 |
| 1378926 | 9.35 |
| 1386628 | 8.82 |
| 1398889 | 9.44 |
| 1439915 | 8.82 |
| 1448638 | 8.91 |
| 1457223 | 9.44 |
| 1486763 | 9.35 |
| 1487980 | 9.44 |
| 1495055 | 8.91 |
| 1499663 | 9 |
| 1509582 | 8.19 |
| 1514967 | 9.35 |
| 1526839 | 6.92 |
| 1538496 | 8.19 |
| 1564650 | 9.35 |
| 1573301 | 8.91 |
| 1582681 | 9 |
| 1587216 | 9.18 |
| 1745478 | 8.54 |
| 1745873 | 9.44 |
| 1766031 | 9.53 |
| 1784498 | 9.62 |
| 1795104 | 9.53 |
| 1802552 | 9.53 |
| 1814132 | 9.53 |
| 1829928 | 9.8 |
| 1837216 | 9.62 |
| 1842772 | 9.62 |
| 1856803 | 9.62 |
| 1865329 | 9.62 |
| 1874298 | 9.71 |
| 1883689 | 9.62 |
| 1893981 | 9.44 |
| 1895584 | 9.53 |
| 1917586 | 9.71 |
| 1932291 | 7.81 |
| 1937393 | 9.8 |
| 1943909 | 9.71 |
| 1970310 | 9.62 |
| 1975825 | 8.82 |
| 1987320 | 7.81 |
| 1998699 | 9.53 |
| 2007627 | 10.07 |
| 2013466 | 10.25 |
| 2017455 | 10.07 |
| 2023963 | 9.89 |
| 2035877 | 9.53 |
| 2045160 | 10.07 |
| 2066412 | 9.8 |
| 2079262 | 10.07 |
| 2087111 | 10.33 |
| 2096409 | 10.42 |
| 2117287 | 10.07 |
| 2120130 | 9.44 |
| 2125869 | 10.42 |
| 2148278 | 10.69 |
| 2155665 | 10.51 |
| 2165481 | 10.42 |
| 2179202 | 10.42 |
| 2471624 | 10.42 |
| 2526745 | 10.69 |
| 2560578 | 10.25 |
| 2573662 | 10.69 |
| 2681138 | 10.87 |
| 2714014 | 10.78 |
| 2717510 | 11.49 |
| 2723785 | 10.42 |
| 2731580 | 10.69 |
| 2741689 | 10.96 |
| 2759887 | 10.69 |
| 2773698 | 10.78 |
| 2780605 | 10.87 |
| 2789793 | 11.22 |
| 2811773 | 10.96 |
| 2838817 | 10.96 |
| 2841607 | 10.25 |
| 2846840 | 10.96 |
| 2852319 | 10.96 |
| 2863298 | 10.96 |
| 2872703 | 10.96 |
| 2902893 | 11.31 |
| 2904450 | 11.31 |
| 2912569 | 10.96 |
| 2930989 | 11.58 |
| 2937963 | 11.31 |
| 2951467 | 10.96 |
| 2962637 | 11.76 |
| 2963136 | 11.76 |
| 2999670 | 10.78 |
| 3003410 | 11.31 |
| 3012066 | 11.31 |
| 3039180 | 11.94 |
| 3043018 | 11.31 |
| 3047019 | 11.22 |
| 3274129 | 8.19 |
| 3492286 | 8.82 |
| 3746801 | 8.37 |
| 3755548 | 9.18 |
| 3774998 | 7.81 |
| 3819300 | 7.81 |
| 3840981 | 9.18 |
| 3852408 | 8.54 |
| 3857142 | 9.18 |
| 3869519 | 7.81 |
| 3873196 | 7.81 |
| 3877576 | 8.37 |
| 3900305 | 8.54 |
| 3940932 | 8.37 |
| 3956134 | 8.64 |
| 3984757 | 8.37 |
| 4014959 | 8.19 |
| 4021529 | 8.91 |
| 4025353 | 9.18 |
| 4038869 | 8.64 |
| 4048055 | 6.33 |
| 4054812 | 8.37 |
| 4060950 | 8.37 |
| 4090760 | 8.37 |
| 4099331 | 9.18 |
| 4108916 | 8.19 |
| 4144992 | 7.33 |
| 4168051 | 9.18 |
| 4179349 | 5.61 |
| 4191826 | 7.81 |
| 4213056 | 9.18 |
| 4215484 | 7.22 |
| 4222383 | 8.54 |
| 4238274 | 8.54 |
| 4239877 | 8.54 |
| 4245335 | 8.54 |
| 4256329 | 7.81 |
| 4265792 | 9 |
| 4279031 | 7.22 |
| 4428224 | 9.18 |
| 4586541 | 8.37 |
| 5097676 | 8.28 |
| 5108542 | 8.82 |
| 5116281 | 9.18 |
| 5124193 | 8.64 |
| 5127287 | 8.19 |
| 5134115 | 8.19 |
| 5137082 | 8.54 |
| 5143433 | 8.19 |
| 5153129 | 8.82 |
| 5156887 | 9.18 |
| 5166199 | 8.64 |
| 5174104 | 8.82 |
| 5175401 | 8.82 |
| 5270183 | 9.26 |
| 5578236 | 8.37 |
| 5826818 | 7.81 |

LG10

|  | genetic_pos |
| --- | --- |
| 77909 | 3.37 |
| 79012 | 3.09 |
| 88335 | 3.09 |
| 95804 | 2.53 |
| 114816 | 3.09 |
| 264349 | 2.73 |
| 378960 | 2.73 |
| 384664 | 2.73 |
| 405256 | 2.73 |
| 428339 | 2.53 |
| 467676 | 2.82 |
| 479636 | 3.09 |
| 489674 | 3.09 |
| 498857 | 0 |
| 509978 | 3.09 |
| 516037 | 2.73 |
| 531243 | 2.82 |
| 541537 | 3.09 |
| 552028 | 2.53 |
| 556041 | 3.09 |
| 575431 | 2.53 |
| 585006 | 3.09 |
| 594862 | 3.83 |
| 596335 | 3.09 |
| 615092 | 3.09 |
| 632262 | 2.73 |
| 638533 | 3.09 |
| 655105 | 3.09 |
| 676413 | 2.73 |
| 684308 | 2.53 |
| 695888 | 2.73 |
| 699416 | 2.02 |
| 702677 | 2.53 |
| 715194 | 1.58 |
| 735390 | 2.43 |
| 748769 | 3.83 |
| 764538 | 2.53 |
| 774829 | 2.43 |
| 775486 | 2.53 |
| 825345 | 2.73 |
| 832265 | 3.56 |
| 919773 | 2.73 |
| 929784 | 2.82 |
| 936552 | 2.91 |
| 947528 | 2.43 |
| 966635 | 3.09 |
| 978597 | 2.43 |
| 986790 | 2.53 |
| 1013298 | 2.82 |
| 1018985 | 2.73 |
| 1028990 | 2.53 |
| 1039077 | 1.31 |
| 1050131 | 2.53 |
| 1055468 | 2.82 |
| 1063375 | 1.92 |
| 1065472 | 2.82 |
| 1066090 | 1.17 |
| 1076367 | 2.73 |
| 1076776 | 2.73 |
| 1086738 | 2.43 |
| 1111079 | 3.83 |
| 1113256 | 2.43 |
| 1131089 | 2.02 |
| 1151162 | 2.53 |
| 1157440 | 2.43 |
| 1190862 | 3.83 |
| 1201739 | 2.43 |
| 1208324 | 2.53 |
| 1225237 | 2.53 |
| 1232551 | 2.73 |
| 1244565 | 2.91 |
| 1296954 | 2.73 |
| 1343778 | 2.53 |
| 1352939 | 2.73 |
| 1356276 | 3 |
| 1364260 | 2.82 |
| 1374371 | 2.73 |
| 1384597 | 2.53 |
| 1401359 | 2.73 |
| 1412210 | 2.73 |
| 1422083 | 3.92 |
| 1433326 | 2.73 |
| 1441764 | 2.91 |
| 1453489 | 2.53 |
| 1461619 | 2.82 |
| 1499239 | 2.82 |
| 1502139 | 3 |
| 1511254 | 3.83 |
| 1524127 | 3.83 |
| 1787022 | 5.29 |
| 1788863 | 3.56 |
| 1791860 | 3.83 |
| 1805873 | 2.43 |
| 1812894 | 2.73 |
| 1872044 | 2.73 |
| 1873501 | 2.73 |
| 1892672 | 2.73 |
| 1894905 | 2.53 |
| 2014035 | 3.09 |
| 2027133 | 3.37 |
| 2031754 | 2.53 |
| 2037112 | 1.81 |
| 2055949 | 2.73 |
| 2056322 | 2.73 |
| 2064638 | 2.91 |
| 2073199 | 3.09 |
| 2073998 | 3.83 |
| 2095207 | 2.73 |
| 2097958 | 2.73 |
| 2104556 | 1.31 |
| 2112452 | 3.92 |
| 2142267 | 2.73 |
| 2170627 | 3.56 |
| 2380759 | 2.73 |
| 2416673 | 2.43 |
| 2425696 | 2.73 |
| 2456786 | 3.09 |
| 2488099 | 2.53 |
| 2496224 | 2.02 |
| 2497802 | 2.43 |
| 2499053 | 2.43 |
| 2506350 | 2.43 |
| 2519339 | 3.09 |
| 2521969 | 3.47 |
| 2529649 | 2.73 |
| 2566280 | 2.73 |
| 2575496 | 2.73 |
| 2578539 | 2.73 |
| 2715842 | 2.43 |
| 2732811 | 2.82 |
| 2758490 | 2.53 |
| 2765133 | 2.43 |
| 2784511 | 2.73 |
| 2801954 | 2.73 |
| 2846052 | 2.43 |
| 2852679 | 2.73 |
| 2857464 | 2.82 |
| 2876817 | 3.09 |
| 2919133 | 2.73 |
| 2928312 | 2.73 |
| 2938854 | 3.09 |
| 2942955 | 2.43 |
| 2947583 | 3.09 |
| 3001882 | 2.53 |
| 3013787 | 3.09 |
| 3027112 | 2.02 |
| 3039985 | 2.43 |
| 3080976 | 2.73 |
| 3086165 | 2.53 |
| 3098446 | 2.82 |
| 3105408 | 2.53 |
| 3117081 | 2.53 |
| 3130571 | 2.73 |
| 3137676 | 2.73 |
| 3142572 | 2.53 |
| 3340874 | 2.82 |
| 3343917 | 2.73 |
| 3345344 | 2.73 |
| 3346576 | 2.73 |
| 3347679 | 2.82 |
| 3351103 | 2.73 |
| 3359348 | 1.31 |
| 3635318 | 0.86 |
| 3880288 | 2.73 |
| 3885488 | 1.31 |
| 3889971 | 2.73 |
| 3890221 | 3.56 |
| 3908203 | 2.73 |
| 3929225 | 2.53 |
| 3939808 | 2.82 |
| 3965483 | 2.73 |
| 3965928 | 2.73 |
| 3970392 | 2.82 |
| 3990021 | 2.73 |
| 4027889 | 2.02 |
| 4032049 | 2.73 |
| 4039348 | 1.81 |
| 4048901 | 2.02 |
| 4073969 | 3.92 |
| 4084340 | 2.82 |
| 4087959 | 2.53 |
| 4111063 | 2.53 |
| 4135515 | 2.53 |
| 4156273 | 3.83 |
| 4194887 | 2.73 |
| 4215015 | 3.09 |
| 4246114 | 3.09 |
| 4269187 | 3.09 |
| 4341226 | 2.02 |
| 4480974 | 3.09 |
| 4501299 | 3.09 |
| 4503693 | 3.09 |
| 4513777 | 3.09 |
| 4517522 | 3.09 |
| 4517762 | 1.17 |
| 4557948 | 2.53 |
| 4574602 | 2.53 |
| 4599623 | 2.73 |
| 4609866 | 3.09 |
| 4619162 | 2.82 |

LG11

|  | genetic_pos |
| --- | --- |
| 479984 | 4.41 |
| 513149 | 5.29 |
| 516246 | 4.41 |
| 655191 | 4.41 |
| 715154 | 4.9 |
| 721121 | 5.29 |
| 732277 | 6.77 |
| 733673 | 4.41 |
| 771976 | 4.5 |
| 786181 | 4.7 |
| 813647 | 4.99 |
| 854009 | 4.5 |
| 964523 | 4.7 |
| 979102 | 5.76 |
| 984930 | 5.29 |
| 995651 | 6.4 |
| 1059246 | 4.9 |
| 1114483 | 4.5 |
| 1114691 | 4.41 |
| 1128397 | 4.41 |
| 1147209 | 4.41 |
| 1166701 | 4.7 |
| 1174976 | 5.29 |
| 1204788 | 5.76 |
| 1216451 | 5.29 |
| 1222564 | 3.34 |
| 1232194 | 6.5 |
| 1240685 | 4.41 |
| 1260297 | 4.7 |
| 1408826 | 4.41 |
| 1771295 | 5.48 |
| 1780720 | 5.29 |
| 1792579 | 5.18 |
| 1835367 | 4.99 |
| 1879290 | 4.99 |
| 1887030 | 4.99 |
| 1900347 | 5.29 |
| 1908955 | 6.4 |
| 1917661 | 5.29 |
| 1935964 | 5.38 |
| 1951777 | 5.38 |
| 1965170 | 5.29 |
| 1973700 | 5.48 |
| 1982766 | 4.99 |
| 1983318 | 6.4 |
| 2050579 | 4.99 |
| 2065379 | 5.29 |
| 2084717 | 4.41 |
| 2157306 | 5.48 |
| 2172898 | 4.99 |
| 2209285 | 4.9 |
| 2213861 | 6.4 |
| 2251916 | 5.18 |
| 2273041 | 3.12 |
| 2281901 | 4.9 |
| 2306634 | 5.29 |
| 2325956 | 5.18 |
| 2357731 | 5.29 |
| 2365768 | 4.9 |
| 2367720 | 5.85 |
| 2376862 | 5.38 |
| 2386313 | 4.9 |
| 2395666 | 6.4 |
| 2404025 | 5.29 |
| 2405739 | 4.9 |
| 2407170 | 5.48 |
| 2423412 | 4.9 |
| 2432561 | 6.4 |
| 2444416 | 4.02 |
| 2456229 | 5.76 |
| 2466632 | 4.5 |
| 2476147 | 4.9 |
| 2504628 | 5.29 |
| 2512022 | 5.38 |
| 2523867 | 4.7 |
| 2532846 | 5.29 |
| 2540350 | 4.5 |
| 2557278 | 6.4 |
| 2599152 | 5.29 |
| 2606644 | 5.29 |
| 2609502 | 6.4 |
| 2614312 | 5.29 |
| 2614502 | 5.85 |
| 2614905 | 4.9 |
| 2652545 | 5.29 |
| 2670111 | 5.18 |
| 2694582 | 5.29 |
| 2711666 | 5.29 |
| 2716653 | 5.76 |
| 2728245 | 5.29 |
| 2734964 | 5.29 |
| 2862311 | 5.29 |
| 2869351 | 5.29 |
| 2881358 | 4.9 |
| 2884120 | 4.9 |
| 2891075 | 4.5 |
| 2904341 | 6.4 |
| 2914911 | 4.5 |
| 2921210 | 5.29 |
| 2922102 | 4.9 |
| 2934391 | 5.29 |
| 2959489 | 5.29 |
| 2966639 | 4.7 |
| 2978694 | 5.29 |
| 2991614 | 4.99 |
| 2997202 | 5.29 |
| 3009393 | 5.29 |
| 3027058 | 5.48 |
| 3065850 | 5.76 |
| 3078474 | 4.5 |
| 3098167 | 5.48 |
| 3109130 | 4.02 |
| 3470468 | 4.5 |
| 3476480 | 4.41 |
| 3485262 | 4.5 |
| 3536909 | 4.5 |
| 3547395 | 4.5 |
| 3626859 | 5.38 |
| 3651940 | 4.5 |
| 3802221 | 4.5 |
| 3821481 | 5.48 |
| 3853547 | 4.9 |
| 3880708 | 4.81 |
| 3907671 | 4.99 |
| 3930747 | 4.5 |
| 3939303 | 5.48 |
| 4142998 | 5.29 |
| 4179104 | 5.48 |
| 4190887 | 5.38 |
| 4199534 | 5.48 |
| 4331597 | 5.48 |
| 4515221 | 5.48 |
| 4578330 | 5.48 |
| 5414324 | 5.48 |
| 5432352 | 5.38 |
| 5438577 | 5.48 |
| 5547467 | 5.38 |
| 5577825 | 2.65 |
| 5587004 | 5.38 |
| 5596676 | 5.48 |
| 5723479 | 4.9 |
| 5731963 | 5.29 |
| 5732343 | 5.48 |
| 5753433 | 3.46 |
| 5806528 | 4.9 |
| 5820675 | 4.9 |
| 5828406 | 4.9 |
| 5925779 | 6.4 |
| 5977615 | 4.5 |
| 5983283 | 5.38 |
| 6002806 | 5.38 |
| 6097171 | 5.29 |
| 6135888 | 5.76 |
| 6179036 | 5.48 |
| 6201201 | 5.38 |
| 6211106 | 4.41 |
| 6231183 | 5.58 |
| 6240095 | 4.5 |
| 6253052 | 4.41 |
| 6263188 | 5.48 |
| 6312383 | 5.48 |
| 6318944 | 5.48 |
| 6379528 | 4.41 |
| 6392820 | 5.76 |
| 6428385 | 5.48 |
| 6456733 | 5.48 |
| 6467761 | 5.58 |
| 6478456 | 5.38 |
| 6490340 | 5.48 |
| 6499156 | 4.5 |
| 6508257 | 5.76 |
| 6540084 | 6.23 |
| 6559472 | 4.5 |
| 6570621 | 5.38 |
| 6577978 | 5.38 |
| 6588402 | 6.77 |
| 6601812 | 6.4 |
| 6609875 | 4.32 |
| 6621610 | 5.48 |
| 6631414 | 5.76 |
| 6639816 | 5.38 |
| 6678724 | 2.49 |
| 6679642 | 4.5 |
| 6680471 | 4.5 |
| 6686757 | 4.5 |
| 6695402 | 4.5 |
| 6706336 | 5.38 |
| 6891626 | 5.76 |
| 6948912 | 4.5 |
| 6959922 | 0 |
| 6976526 | 5.76 |
| 6994286 | 5.38 |
| 7007136 | 4.5 |
| 7015218 | 5.48 |
| 7023744 | 4.5 |
| 7026577 | 4.5 |
| 7029822 | 5.76 |
| 7038374 | 5.48 |
| 7047291 | 5.48 |
| 7065779 | 4.5 |

LG12

|  | genetic_pos |
| --- | --- |
| 639087 | 5.54 |
| 659570 | 5.54 |
| 666130 | 6.04 |
| 689980 | 5.54 |
| 710779 | 5.24 |
| 721980 | 5.24 |
| 751275 | 6.14 |
| 752560 | 5.54 |
| 773415 | 5.64 |
| 780391 | 6.04 |
| 820267 | 6.04 |
| 882262 | 6.04 |
| 922974 | 5.54 |
| 934613 | 5.54 |
| 951619 | 6.14 |
| 964657 | 5.54 |
| 966670 | 5.34 |
| 997528 | 5.54 |
| 1006186 | 5.64 |
| 1015171 | 5.64 |
| 1016272 | 5.94 |
| 1220490 | 5.64 |
| 1230788 | 6.14 |
| 1233420 | 4.82 |
| 1424179 | 6.04 |
| 1427830 | 5.54 |
| 1461780 | 4.92 |
| 1481091 | 5.44 |
| 1532067 | 6.04 |
| 1543580 | 4.92 |
| 1573538 | 5.44 |
| 1609794 | 4.82 |
| 1621494 | 6.04 |
| 1661292 | 5.64 |
| 1746466 | 5.64 |
| 1757916 | 6.04 |
| 1762009 | 4.82 |
| 1781541 | 4.82 |
| 1785869 | 5.64 |
| 1798438 | 5.54 |
| 1930745 | 4.82 |
| 1938258 | 6.04 |
| 1947972 | 5.44 |
| 2054393 | 6.34 |
| 2065537 | 6.14 |
| 2085775 | 6.04 |
| 2086888 | 5.54 |
| 2093856 | 4.82 |
| 2103823 | 6.14 |
| 2137701 | 5.54 |
| 2159343 | 6.04 |
| 2176769 | 5.64 |
| 2185964 | 6.04 |
| 2193573 | 5.54 |
| 3069804 | 6.04 |
| 3323675 | 5.64 |
| 3329344 | 6.04 |
| 3330536 | 4.92 |
| 3336406 | 5.44 |
| 3348626 | 6.04 |
| 3354524 | 5.84 |
| 3418904 | 4.82 |
| 3421579 | 4.82 |
| 3437732 | 5.44 |
| 3443413 | 6.04 |
| 3474228 | 5.44 |
| 3689659 | 0 |
| 3706814 | 5.44 |
| 3882345 | 6.14 |
| 3886299 | 5.44 |
| 3886740 | 4.82 |
| 3890481 | 6.04 |
| 3898975 | 5.54 |
| 3954468 | 4.82 |
| 4006734 | 5.24 |
| 4063662 | 5.54 |
| 4069544 | 6.04 |
| 4101194 | 6.14 |
| 4106515 | 5.54 |
| 4131936 | 6.04 |
| 4192195 | 6.04 |
| 4206087 | 4.71 |
| 4272087 | 5.54 |
| 4527167 | 6.04 |
| 4550306 | 5.44 |
| 4576030 | 6.14 |
| 4587128 | 6.04 |
| 4646544 | 6.14 |
| 4669135 | 5.54 |
| 4707144 | 6.04 |
| 4729149 | 6.04 |
| 4749829 | 6.04 |
| 4753560 | 6.04 |
| 4817423 | 4.92 |
| 5336534 | 6.14 |
| 5347776 | 6.04 |
| 5476162 | 6.04 |
| 5485720 | 6.04 |
| 5510027 | 5.14 |
| 5522301 | 6.04 |
| 5529206 | 5.24 |
| 5553095 | 5.54 |
| 5561620 | 5.44 |
| 5621401 | 5.44 |
| 5623513 | 5.64 |
| 5659521 | 5.54 |
| 5666935 | 6.14 |
| 5691456 | 6.14 |
| 5701269 | 6.04 |
| 5721412 | 5.54 |
| 5742766 | 6.14 |
| 5750753 | 5.44 |
| 5784925 | 6.04 |
| 5875094 | 6.04 |
| 5889350 | 5.94 |
| 5890852 | 5.54 |
| 5893925 | 6.04 |
| 5895985 | 6.04 |
| 5902274 | 6.04 |
| 5910174 | 2.19 |
| 5917358 | 4.82 |
| 5922180 | 4.82 |
| 5925300 | 6.04 |
| 5927595 | 6.04 |
| 5930144 | 5.54 |
| 5943894 | 6.04 |
| 5952607 | 5.44 |
| 5955637 | 4.82 |
| 5956454 | 5.64 |
| 5966869 | 5.24 |
| 5969687 | 5.44 |
| 5980387 | 5.24 |
| 5984775 | 6.04 |
| 6034926 | 5.44 |
| 6054371 | 6.04 |
| 6063836 | 6.14 |
| 6069816 | 6.04 |
| 6108687 | 5.44 |
| 6109730 | 4.82 |
| 6122625 | 4.82 |
| 6131673 | 6.04 |
| 6159539 | 4.82 |
| 6164118 | 6.04 |
| 6165587 | 4.82 |
| 6211402 | 6.04 |
| 6211983 | 6.14 |
| 6222875 | 6.14 |
| 6223408 | 4.82 |
| 6225058 | 6.34 |
| 6322489 | 5.64 |
| 6331318 | 4.82 |
| 6731000 | 6.04 |
| 6762172 | 6.04 |
| 6802485 | 6.04 |
| 6807443 | 5.64 |
| 6813695 | 6.04 |
| 6815930 | 6.04 |
| 6822683 | 6.04 |
| 6836296 | 4.82 |
| 6881347 | 6.34 |
| 6902082 | 5.64 |
| 6917430 | 4.82 |
| 6925621 | 5.54 |
| 6935956 | 6.84 |
| 6945531 | 6.04 |
| 6954501 | 5.84 |
| 6960479 | 6.04 |
| 6968135 | 6.44 |
| 6978824 | 3.92 |
| 6992149 | 5.64 |
| 7008681 | 5.64 |
| 7016000 | 4.82 |
| 7021257 | 4.92 |
| 7039191 | 6.04 |
| 7043405 | 4.49 |
| 7070148 | 6.04 |
| 7077707 | 6.04 |
| 7081473 | 5.44 |
| 7091731 | 4.82 |
| 7094293 | 6.04 |
| 7097381 | 5.14 |
| 7098293 | 6.14 |
| 7100139 | 5.64 |
| 7102777 | 5.64 |
| 7108273 | 6.04 |
| 7112527 | 6.84 |
| 7122310 | 6.14 |
| 7133162 | 4.82 |
| 7137516 | 5.24 |
| 7243884 | 6.04 |
| 7248637 | 6.94 |
| 7249120 | 6.14 |
| 7262386 | 5.94 |
| 7265887 | 3.15 |
| 7275454 | 4.82 |
| 7278427 | 5.34 |
| 7282436 | 5.34 |
| 7292968 | 5.03 |
| 7304562 | 6.04 |
| 7315622 | 6.04 |

LG13

|  | genetic_pos |
| --- | --- |
| 110954 | 3.66 |
| 168700 | 4.15 |
| 188855 | 3.05 |
| 218403 | 3.29 |
| 395181 | 3.66 |
| 401283 | 4.27 |
| 465586 | 3.66 |
| 471416 | 3.66 |
| 521985 | 3.66 |
| 524777 | 3.29 |
| 540974 | 2.08 |
| 542866 | 3.29 |
| 583476 | 0.51 |
| 607910 | 3.29 |
| 615150 | 3.66 |
| 638177 | 3.29 |
| 645622 | 3.29 |
| 679780 | 3.05 |
| 690468 | 4.27 |
| 727078 | 4.27 |
| 727734 | 4.15 |
| 728973 | 3.29 |
| 749582 | 3.29 |
| 755401 | 3.05 |
| 757837 | 4.15 |
| 763783 | 3.66 |
| 844223 | 3.66 |
| 853813 | 3.29 |
| 874400 | 3.05 |
| 887132 | 3.05 |
| 936328 | 3.29 |
| 946808 | 3.29 |
| 975551 | 3.05 |
| 985930 | 3.05 |
| 1004573 | 3.66 |
| 1028804 | 3.29 |
| 1030398 | 3.05 |
| 1081826 | 2.92 |
| 1083448 | 3.05 |
| 1100743 | 3.66 |
| 1146423 | 4.15 |
| 1166825 | 3.05 |
| 1176160 | 2.92 |
| 1186989 | 3.05 |
| 1220843 | 3.66 |
| 1234271 | 3.29 |
| 1247937 | 3.05 |
| 1258371 | 4.15 |
| 1270082 | 3.66 |
| 1293435 | 2.5 |
| 1303263 | 3.05 |
| 1305837 | 0 |
| 1388583 | 3.29 |
| 1401616 | 3.66 |
| 1441292 | 3.29 |
| 1444532 | 1.63 |
| 1444924 | 3.29 |
| 1460188 | 3.05 |
| 1469501 | 3.66 |
| 1516978 | 3.66 |
| 1532196 | 4.15 |
| 1541525 | 4.27 |
| 1542473 | 3.66 |
| 1652707 | 4.15 |
| 1675601 | 3.29 |
| 1726907 | 4.15 |
| 1742215 | 4.15 |
| 1747515 | 3.66 |
| 1754701 | 4.15 |
| 1785877 | 3.05 |
| 1786389 | 4.15 |
| 1787017 | 4.15 |
| 1796927 | 3.54 |
| 1803755 | 3.05 |
| 1813913 | 3.66 |
| 1848878 | 0.85 |
| 1905106 | 3.29 |
| 1917201 | 3.05 |
| 1928373 | 3.66 |
| 1945242 | 4.15 |
| 1951417 | 3.05 |
| 1956408 | 3.29 |
| 1977502 | 2.78 |
| 1994623 | 3.66 |
| 2005316 | 3.29 |
| 2024824 | 3.66 |
| 2067659 | 3.05 |
| 2076622 | 3.29 |
| 2120883 | 3.29 |
| 2128577 | 3.66 |
| 2131285 | 4.27 |
| 2210724 | 4.39 |
| 2219228 | 2.08 |
| 2220996 | 4.15 |
| 2262117 | 3.66 |
| 2262637 | 3.05 |
| 2262996 | 4.15 |
| 2264386 | 4.27 |
| 2283789 | 3.66 |
| 2284290 | 3.05 |
| 2336982 | 3.05 |
| 2346161 | 3.05 |
| 2355796 | 3.29 |
| 2367242 | 3.05 |
| 2370038 | 3.66 |
| 2378368 | 3.05 |
| 2531883 | 3.66 |
| 2539528 | 4.15 |
| 2557692 | 3.05 |
| 2561451 | 3.66 |
| 2574793 | 4.15 |
| 2593642 | 4.15 |
| 2631538 | 3.66 |
| 2642818 | 3.05 |
| 2658158 | 2.92 |
| 2692912 | 3.66 |
| 2718864 | 3.05 |
| 2719478 | 3.66 |
| 2741455 | 3.66 |
| 2752717 | 4.15 |
| 2753619 | 4.15 |
| 2761389 | 3.66 |
| 2769685 | 3.05 |
| 2770432 | 3.66 |
| 2780525 | 4.15 |
| 2801191 | 4.15 |
| 2803448 | 3.05 |
| 2928049 | 4.89 |
| 2999230 | 3.66 |
| 3011549 | 3.66 |
| 3020229 | 3.29 |
| 3036068 | 3.29 |
| 3038121 | 4.15 |
| 3039594 | 3.66 |
| 3040692 | 3.05 |
| 3373021 | 3.29 |
| 4091504 | 4.15 |
| 4119048 | 3.54 |
| 4130986 | 3.05 |
| 4144182 | 4.15 |
| 4176269 | 3.29 |
| 4199673 | 2.92 |
| 4210113 | 3.29 |
| 4210317 | 3.66 |
| 4219092 | 3.66 |
| 4223462 | 4.15 |
| 4224905 | 3.66 |
| 4347920 | 4.15 |
| 4411709 | 4.89 |
| 4433213 | 4.76 |
| 4439740 | 4.76 |
| 4450072 | 3.05 |
| 4489525 | 4.76 |
| 4526918 | 4.76 |
| 4535856 | 4.76 |
| 4556774 | 4.76 |
| 4564622 | 4.76 |
| 4573847 | 4.89 |
| 4593994 | 4.89 |
| 4616872 | 3.29 |
| 4623681 | 4.76 |
| 4636469 | 4.76 |
| 4649129 | 4.76 |
| 4655085 | 2.78 |
| 4667682 | 4.76 |
| 4679175 | 6.45 |
| 4781552 | 6.45 |
| 4976016 | 6.57 |
| 5029581 | 6.45 |
| 5052320 | 6.57 |
| 5065994 | 6.45 |
| 5084019 | 6.57 |
| 5094746 | 6.57 |
| 5103906 | 6.45 |
| 5110790 | 6.45 |
| 5131042 | 6.45 |
| 5137996 | 6.45 |
| 5149010 | 6.45 |
| 5159408 | 6.45 |
| 5168697 | 6.57 |
| 5179029 | 6.45 |
| 5190317 | 4.89 |
| 5210483 | 6.57 |
| 5221839 | 6.57 |
| 5235534 | 6.32 |
| 5244077 | 6.57 |
| 5262116 | 6.57 |
| 5284401 | 4.89 |
| 5302711 | 6.45 |
| 5304787 | 6.45 |
| 5315473 | 6.32 |
| 5325679 | 7.05 |
| 5328202 | 6.57 |
| 5338498 | 4.89 |
| 5360140 | 6.45 |
| 5361739 | 6.45 |
| 5362168 | 6.32 |
| 5362629 | 6.45 |
| 5363280 | 6.57 |
| 5376142 | 6.45 |

LG14

|  | genetic_pos |
| --- | --- |
| 204208 | 8.18 |
| 210976 | 8.57 |
| 214248 | 8.38 |
| 218657 | 8.18 |
| 229182 | 8.08 |
| 230660 | 9.13 |
| 235834 | 8.18 |
| 239365 | 9.04 |
| 254412 | 8.18 |
| 261740 | 9.04 |
| 271237 | 8.47 |
| 452749 | 8.38 |
| 465773 | 9.04 |
| 475058 | 8.08 |
| 487011 | 8.08 |
| 569057 | 8.08 |
| 613112 | 9.23 |
| 617651 | 9.04 |
| 620488 | 8.08 |
| 671506 | 8.66 |
| 741272 | 8.75 |
| 754430 | 9.04 |
| 760233 | 9.13 |
| 763892 | 8.38 |
| 778423 | 8.08 |
| 796142 | 9.04 |
| 796854 | 9.13 |
| 810652 | 9.04 |
| 815468 | 8.18 |
| 826244 | 8.08 |
| 834984 | 9.13 |
| 837605 | 8.38 |
| 837866 | 8.18 |
| 841811 | 8.66 |
| 854557 | 9.13 |
| 864474 | 9.13 |
| 868109 | 8.08 |
| 877676 | 9.04 |
| 883705 | 9.23 |
| 894485 | 9.13 |
| 929102 | 9.13 |
| 936704 | 9.04 |
| 955933 | 8.08 |
| 974871 | 8.18 |
| 983006 | 9.13 |
| 986205 | 8.57 |
| 987425 | 8.18 |
| 1188155 | 9.04 |
| 1247003 | 8.18 |
| 1256854 | 8.38 |
| 1601568 | 8.85 |
| 1610038 | 9.52 |
| 1626013 | 9.52 |
| 1627379 | 9.52 |
| 1651750 | 9.52 |
| 1658715 | 9.61 |
| 1663573 | 9.61 |
| 1679455 | 8.66 |
| 1679854 | 8.85 |
| 1691113 | 9.52 |
| 1710206 | 9.52 |
| 1713388 | 9.52 |
| 1731981 | 8.85 |
| 1752374 | 9.52 |
| 1777650 | 9.52 |
| 1779406 | 9.52 |
| 1783431 | 9.52 |
| 1803893 | 9.42 |
| 1811922 | 8.85 |
| 1822072 | 9.23 |
| 1842939 | 9.42 |
| 1868774 | 9.23 |
| 1985014 | 4.61 |
| 1992466 | 9.42 |
| 2005403 | 9.52 |
| 2007748 | 9.42 |
| 2023059 | 9.42 |
| 2034266 | 9.04 |
| 2043927 | 8.57 |
| 2090825 | 9.52 |
| 2112322 | 9.52 |
| 2112906 | 9.52 |
| 2225886 | 8.75 |
| 2345905 | 9.13 |
| 2356184 | 8.85 |
| 2371526 | 9.04 |
| 2383697 | 9.23 |
| 2396563 | 9.52 |
| 2401273 | 9.52 |
| 2403381 | 8.57 |
| 2425259 | 9.52 |
| 2451742 | 8.38 |
| 2462039 | 9.23 |
| 2483856 | 8.38 |
| 2502026 | 9.52 |
| 2506838 | 9.42 |
| 2519359 | 9.13 |
| 2530092 | 9.52 |
| 2549071 | 8.85 |
| 2566315 | 9.04 |
| 2586701 | 9.13 |
| 2589069 | 9.04 |
| 2598045 | 9.04 |
| 2610552 | 9.13 |
| 2620383 | 9.04 |
| 2651353 | 9.23 |
| 2688592 | 9.13 |
| 2702566 | 9.04 |
| 2711028 | 9.13 |
| 2727483 | 9.04 |
| 2734068 | 9.04 |
| 2750941 | 9.04 |
| 2792539 | 8.85 |
| 2881880 | 9.04 |
| 2924766 | 9.04 |
| 2973957 | 9.04 |
| 2977342 | 9.04 |
| 2990230 | 9.04 |
| 2997690 | 9.13 |
| 3015555 | 9.13 |
| 3255059 | 8.75 |
| 3262775 | 9.13 |
| 3338983 | 9.04 |
| 3428756 | 9.23 |
| 3588428 | 8.85 |
| 3831286 | 8.85 |
| 3840070 | 8.85 |
| 3901885 | 9.13 |
| 3915875 | 9.13 |
| 4015491 | 7.21 |
| 4044898 | 9.04 |
| 4073196 | 8.47 |
| 4221613 | 9.13 |
| 4225643 | 9.04 |
| 4268931 | 9.13 |
| 4274430 | 9.04 |
| 4276045 | 8.85 |
| 4296409 | 9.04 |
| 4304831 | 9.04 |
| 4314509 | 7.72 |
| 4324612 | 9.13 |
| 4334167 | 9.13 |
| 4350162 | 9.13 |
| 4351855 | 8.38 |
| 4377154 | 9.13 |
| 4484424 | 9.13 |
| 4486650 | 9.04 |
| 4508842 | 9.04 |
| 4608826 | 9.13 |
| 4624470 | 9.04 |
| 4701272 | 9.04 |
| 4711852 | 8.75 |
| 4781659 | 9.13 |
| 4809881 | 9.04 |
| 4820456 | 9.04 |
| 4833493 | 9.13 |
| 4839084 | 9.04 |
| 4853299 | 9.23 |
| 4889070 | 5.2 |
| 4897813 | 9.04 |
| 4902468 | 9.04 |
| 4904832 | 9.13 |
| 4919704 | 9.04 |
| 4929822 | 8.85 |
| 5193827 | 9.04 |
| 5250739 | 9.23 |
| 5297172 | 9.13 |
| 5306212 | 9.04 |
| 5318276 | 9.04 |
| 5326986 | 9.04 |
| 5493330 | 9.04 |
| 5529463 | 9.04 |
| 5541343 | 9.13 |
| 5594760 | 9.13 |
| 5619417 | 9.13 |
| 5772212 | 9.04 |
| 5818580 | 9.04 |
| 5831175 | 9.13 |
| 5899597 | 9.04 |
| 5900653 | 9.04 |
| 5946304 | 9.13 |
| 5951743 | 9.13 |
| 5977743 | 9.13 |
| 5991789 | 9.13 |
| 6238038 | 9.13 |
| 6243885 | 9.13 |
| 6245768 | 9.23 |
| 6274952 | 9.04 |
| 6278451 | 9.13 |
| 6296326 | 9.13 |
| 6307956 | 9.13 |
| 6317297 | 9.13 |
| 6318706 | 9.04 |
| 6329845 | 9.13 |
| 6337275 | 8.57 |
| 6345086 | 9.04 |
| 6368523 | 9.13 |
| 6369135 | 9.04 |
| 6376225 | 8.18 |
| 6400158 | 9.13 |

LG15

|  | genetic_pos |
| --- | --- |
| 22981 | 3.55 |
| 23145 | 3.55 |
| 31726 | 1.98 |
| 37954 | 3.55 |
| 68742 | 3.45 |
| 71682 | 3.45 |
| 79355 | 3.45 |
| 92078 | 2.1 |
| 217613 | 3.45 |
| 223376 | 3.45 |
| 235145 | 3.55 |
| 244963 | 3.45 |
| 245304 | 3.55 |
| 245500 | 3.45 |
| 246126 | 3.45 |
| 250366 | 3.55 |
| 270143 | 3.55 |
| 280805 | 3.45 |
| 318035 | 3.45 |
| 328935 | 1.71 |
| 336760 | 3.66 |
| 348959 | 3.45 |
| 359887 | 2.1 |
| 392337 | 3.45 |
| 415763 | 3.45 |
| 444108 | 3.45 |
| 445037 | 3.45 |
| 454766 | 3.45 |
| 460144 | 2.71 |
| 471703 | 3.45 |
| 486502 | 3.45 |
| 495904 | 3.45 |
| 515563 | 1.98 |
| 527425 | 3.45 |
| 545219 | 3.45 |
| 552133 | 3.45 |
| 560925 | 3.45 |
| 571774 | 3.45 |
| 584097 | 2.1 |
| 593688 | 3.45 |
| 613209 | 3.45 |
| 638607 | 3.45 |
| 658602 | 2.71 |
| 662927 | 3.45 |
| 673627 | 3.45 |
| 678840 | 3.45 |
| 720723 | 3.55 |
| 728399 | 2.21 |
| 733494 | 3.45 |
| 735669 | 3.45 |
| 764651 | 3.25 |
| 765315 | 3.55 |
| 771586 | 3.55 |
| 772563 | 2.1 |
| 778207 | 3.25 |
| 788717 | 3.45 |
| 792075 | 3.55 |
| 792243 | 3.55 |
| 793012 | 3.45 |
| 825678 | 3.16 |
| 859391 | 3.45 |
| 864435 | 3.45 |
| 867396 | 3.45 |
| 874107 | 1.43 |
| 888447 | 3.45 |
| 896829 | 3.45 |
| 905829 | 3.55 |
| 906093 | 3.55 |
| 907281 | 3.45 |
| 919920 | 2.45 |
| 939707 | 2.1 |
| 947841 | 2.1 |
| 958928 | 3.55 |
| 968284 | 2.45 |
| 1015237 | 1.71 |
| 1018518 | 3.45 |
| 1033951 | 3.45 |
| 1065817 | 3.45 |
| 1070840 | 3.45 |
| 1072813 | 3.45 |
| 1072944 | 3.45 |
| 1073119 | 3.45 |
| 1073551 | 1.98 |
| 1081263 | 3.45 |
| 1093385 | 2.21 |
| 1110765 | 0 |
| 1139898 | 3.75 |
| 1156419 | 2.1 |
| 1157380 | 3.45 |
| 1158380 | 3.45 |
| 1158730 | 3.45 |
| 1158866 | 3.45 |
| 1159667 | 3.45 |
| 1160105 | 3.45 |
| 1160605 | 1.98 |
| 1160971 | 3.45 |
| 1167364 | 3.45 |
| 1168545 | 3.55 |
| 1174434 | 3.04 |
| 1180860 | 2.45 |
| 1192811 | 3.45 |
| 1193958 | 3.45 |
| 1203218 | 3.25 |
| 1203649 | 3.45 |
| 1208846 | 3.45 |
| 1215752 | 3.45 |
| 1222302 | 2.21 |
| 1245263 | 3.45 |
| 1326552 | 3.45 |
| 1334640 | 3.45 |
| 1563457 | 3.45 |
| 1600429 | 3.45 |
| 1608956 | 3.45 |
| 1803677 | 2.21 |
| 1921952 | 3.45 |
| 1928356 | 1.71 |
| 1944019 | 3.45 |
| 1952601 | 3.45 |
| 1969823 | 3.45 |
| 1985043 | 2.93 |
| 1994076 | 3.45 |
| 2004770 | 3.45 |
| 2036070 | 3.16 |
| 2147550 | 1.98 |
| 2149639 | 3.45 |
| 2157862 | 3.45 |
| 2168267 | 3.04 |
| 2177098 | 2.93 |
| 2188226 | 3.45 |
| 2196795 | 3.36 |
| 2212637 | 3.45 |
| 2216982 | 3.45 |
| 2233013 | 2.71 |
| 2235748 | 2.71 |
| 2245009 | 3.45 |
| 2258012 | 3.36 |
| 2263728 | 2.1 |
| 2270060 | 3.45 |
| 2276333 | 1.98 |
| 2288410 | 1.86 |
| 2299053 | 3.45 |
| 2302808 | 1.98 |
| 2320365 | 3.16 |
| 2328439 | 3.45 |
| 2348609 | 3.16 |
| 2356669 | 3.45 |
| 2390286 | 3.45 |
| 2410436 | 3.45 |
| 2419632 | 3.45 |
| 2427873 | 3.45 |
| 2449720 | 3.55 |
| 2450870 | 3.45 |
| 2456473 | 3.45 |
| 2469494 | 3.45 |
| 2470136 | 3.45 |
| 2480003 | 3.45 |
| 2493170 | 3.45 |
| 2497590 | 3.55 |
| 2500916 | 1.12 |
| 2523711 | 3.45 |
| 2533758 | 3.16 |
| 2540565 | 3.36 |
| 2550751 | 3.45 |
| 2577749 | 3.55 |
| 2582011 | 3.16 |
| 2609319 | 3.45 |
| 2623584 | 3.55 |
| 2733681 | 3.25 |
| 2801297 | 3.25 |
| 2802040 | 3.16 |
| 2817810 | 3.25 |
| 2829145 | 3.86 |
| 2834331 | 2.93 |
| 2899924 | 3.16 |
| 2901858 | 3.86 |
| 2910668 | 3.86 |
| 2948258 | 3.86 |
| 2953419 | 4.19 |
| 2956089 | 4.19 |
| 2967352 | 3.86 |
| 2977576 | 3.86 |
| 2981089 | 3.16 |
| 2994273 | 3.86 |
| 2994717 | 3.86 |
| 2996937 | 3.16 |
| 3006587 | 3.86 |
| 3013919 | 3.86 |
| 3134641 | 3.86 |
| 3140002 | 3.86 |
| 3144587 | 3.25 |
| 3174385 | 3.16 |
| 3185818 | 3.25 |
| 3185896 | 3.86 |
| 3226747 | 4.3 |
| 3292398 | 4.3 |
| 3303866 | 4.3 |
| 3316915 | 4.3 |
| 3328653 | 4.4 |
| 3335262 | 4.3 |
| 3344007 | 4.3 |

LG16

|  | genetic_pos |
| --- | --- |
| 41003 | 4.46 |
| 46577 | 4.46 |
| 94193 | 4.46 |
| 139147 | 4.46 |
| 143035 | 4.46 |
| 234905 | 4.27 |
| 252774 | 4.27 |
| 262945 | 4.46 |
| 298634 | 4.46 |
| 305849 | 4.18 |
| 321924 | 4.27 |
| 395578 | 4.46 |
| 436949 | 4.46 |
| 442610 | 4.18 |
| 446551 | 4.46 |
| 446812 | 4.46 |
| 450740 | 4.46 |
| 454407 | 4.46 |
| 456933 | 4.46 |
| 459181 | 4.27 |
| 459612 | 3.88 |
| 460459 | 4.27 |
| 460832 | 4.46 |
| 462074 | 4.27 |
| 475420 | 4.46 |
| 480943 | 4.46 |
| 482781 | 4.27 |
| 483180 | 4.46 |
| 487187 | 4.81 |
| 496942 | 4.18 |
| 499660 | 4.46 |
| 506498 | 4.81 |
| 508112 | 4.46 |
| 509240 | 4.46 |
| 509704 | 4.27 |
| 515373 | 4.46 |
| 520948 | 4.46 |
| 521661 | 4.46 |
| 524143 | 4.46 |
| 528106 | 4.46 |
| 530579 | 4.46 |
| 536405 | 4.27 |
| 537549 | 4.46 |
| 538778 | 4.9 |
| 539446 | 4.46 |
| 540610 | 4.46 |
| 543984 | 4.46 |
| 545353 | 4.46 |
| 546358 | 4.46 |
| 546825 | 4.46 |
| 549001 | 4.81 |
| 550533 | 4.27 |
| 553155 | 4.18 |
| 553999 | 4.46 |
| 556560 | 4.46 |
| 556878 | 4.27 |
| 557575 | 4.46 |
| 558723 | 4.27 |
| 565502 | 4.46 |
| 568022 | 4.46 |
| 570352 | 4.81 |
| 571309 | 4.27 |
| 573899 | 4.46 |
| 574401 | 4.46 |
| 576517 | 4.46 |
| 577962 | 4.27 |
| 583266 | 4.18 |
| 588056 | 4.18 |
| 594575 | 4.46 |
| 601044 | 4.18 |
| 605524 | 4.27 |
| 608717 | 4.46 |
| 613118 | 4.46 |
| 668499 | 4.27 |
| 689892 | 4.46 |
| 706885 | 4.27 |
| 711499 | 4.72 |
| 791321 | 4.27 |
| 792983 | 5 |
| 803484 | 3.88 |
| 810079 | 4.46 |
| 810923 | 4.46 |
| 844440 | 4.46 |
| 847755 | 4.46 |
| 863497 | 4.9 |
| 889558 | 4.27 |
| 901988 | 4.46 |
| 929962 | 4.46 |
| 940567 | 4.46 |
| 959285 | 4.18 |
| 1051316 | 4.46 |
| 1067151 | 4.46 |
| 1114976 | 3.28 |
| 1169412 | 4.46 |
| 1177753 | 4.27 |
| 1179812 | 4.18 |
| 1243819 | 4.46 |
| 1244395 | 4.27 |
| 1255517 | 4.46 |
| 1262534 | 4.46 |
| 1262767 | 4.46 |
| 1273254 | 4.46 |
| 1281664 | 4.46 |
| 1322583 | 4.46 |
| 1336072 | 4.46 |
| 1346386 | 4.27 |
| 1349781 | 4.46 |
| 1385042 | 4.46 |
| 1388194 | 4.46 |
| 1388783 | 4.27 |
| 1413428 | 4.27 |
| 1417266 | 4.18 |
| 1422532 | 4.27 |
| 1429333 | 4.46 |
| 1432875 | 4.27 |
| 1441166 | 4.46 |
| 1457752 | 4.46 |
| 1458063 | 4.9 |
| 1471171 | 4.27 |
| 1478487 | 4.27 |
| 1479532 | 4.46 |
| 1531744 | 4.27 |
| 1556761 | 4.46 |
| 1560457 | 4.27 |
| 1574331 | 4.27 |
| 1583258 | 4.46 |
| 1590357 | 4.27 |
| 1599374 | 4.18 |
| 1605607 | 4.27 |
| 1606484 | 4.27 |
| 1647208 | 4.46 |
| 1656889 | 4.46 |
| 1664710 | 4.27 |
| 1672485 | 4.46 |
| 1675926 | 4.27 |
| 1677321 | 4.72 |
| 1683885 | 4.46 |
| 1684925 | 4.27 |
| 1692173 | 4.27 |
| 1696637 | 4.46 |
| 1716729 | 4.46 |
| 1729841 | 4.46 |
| 1752343 | 4.27 |
| 1759345 | 4.46 |
| 1773236 | 4.27 |
| 1778213 | 4.18 |
| 1785129 | 3.88 |
| 1820530 | 4.27 |
| 1843824 | 4.46 |
| 1852861 | 4.46 |
| 1854341 | 4.46 |
| 1854579 | 4.27 |
| 1857403 | 4.27 |
| 1870410 | 4.46 |
| 1874794 | 4.27 |
| 1888290 | 4.27 |
| 1898058 | 4.27 |
| 1953769 | 4.27 |
| 1999129 | 4.27 |
| 2007415 | 4.46 |
| 2010543 | 4.27 |
| 2012097 | 4.27 |
| 2026157 | 4.27 |
| 2035956 | 4.27 |
| 2046354 | 4.18 |
| 2055261 | 4.18 |
| 2060618 | 4.46 |
| 2086036 | 4.9 |
| 2088264 | 4.46 |
| 2091470 | 4.27 |
| 2094075 | 4.46 |
| 2097833 | 4.46 |
| 2099863 | 4.46 |
| 2116020 | 4.27 |
| 2127731 | 4.08 |
| 2148053 | 4.27 |
| 2163480 | 4.46 |
| 2166385 | 4.27 |
| 2220313 | 4.27 |
| 2237882 | 4.46 |
| 2268733 | 4.27 |
| 2270856 | 4.36 |
| 2277209 | 4.46 |
| 2278845 | 4.46 |
| 2285311 | 4.18 |
| 2327132 | 4.46 |
| 2337566 | 4.46 |
| 2347932 | 2 |
| 2356090 | 4.46 |
| 2400712 | 5.09 |
| 2404459 | 5 |
| 2412199 | 5 |
| 2418532 | 5 |
| 2423214 | 5.09 |
| 2428972 | 3.88 |
| 2431159 | 4.46 |
| 2450981 | 4.46 |
| 2466621 | 4.72 |
| 2496738 | 4.46 |
| 2507794 | 4.46 |

LG17

|  | genetic_pos |
| --- | --- |
| 109369 | 4.82 |
| 125353 | 5.11 |
| 417026 | 5.11 |
| 423346 | 5.11 |
| 483976 | 4.72 |
| 526966 | 5.11 |
| 669233 | 4.72 |
| 928903 | 4.72 |
| 964284 | 5.11 |
| 969694 | 4.62 |
| 978369 | 5.11 |
| 1014286 | 5.11 |
| 1017479 | 5.11 |
| 1083700 | 5.11 |
| 1086239 | 4.72 |
| 1089813 | 4.62 |
| 1099542 | 5.11 |
| 1111300 | 4.72 |
| 1182894 | 5.11 |
| 1205618 | 5.21 |
| 1217073 | 5.11 |
| 1222615 | 4.62 |
| 1243878 | 5.11 |
| 1265025 | 5.11 |
| 1273114 | 4.72 |
| 1291565 | 5.11 |
| 1304983 | 5.21 |
| 1307656 | 5.11 |
| 1338391 | 5.11 |
| 1341354 | 5.11 |
| 1438368 | 5.21 |
| 1446633 | 5.21 |
| 1464038 | 5.11 |
| 1476481 | 4.72 |
| 1495718 | 5.11 |
| 1509935 | 4.72 |
| 1513314 | 5.11 |
| 1519060 | 4.62 |
| 1569665 | 5.11 |
| 1578548 | 5.11 |
| 1588015 | 5.21 |
| 1600135 | 5.11 |
| 1613467 | 5.21 |
| 1624378 | 5.21 |
| 1633243 | 5.11 |
| 1641514 | 5.11 |
| 1652002 | 5.11 |
| 1662506 | 5.11 |
| 1689680 | 4.62 |
| 1702167 | 5.11 |
| 1702836 | 4.72 |
| 1711504 | 5.11 |
| 1720715 | 5.21 |
| 1733470 | 5.11 |
| 1741945 | 5.11 |
| 1753987 | 5.11 |
| 1764361 | 5.11 |
| 1783713 | 5.11 |
| 1795431 | 4.62 |
| 1809868 | 4.72 |
| 1813527 | 4.72 |
| 1824653 | 5.11 |
| 1836101 | 5.11 |
| 1968974 | 5.11 |
| 1973731 | 5.11 |
| 2003768 | 5.11 |
| 2017132 | 4.62 |
| 2066701 | 5.11 |
| 2087259 | 5.11 |
| 2098171 | 5.11 |
| 2107728 | 4.91 |
| 2117296 | 5.11 |
| 2139956 | 5.21 |
| 2140406 | 4.72 |
| 2140781 | 5.11 |
| 2147118 | 4.72 |
| 2157977 | 4.72 |
| 2160224 | 5.11 |
| 2179465 | 4.62 |
| 2193681 | 5.11 |
| 2199008 | 5.11 |
| 2209278 | 5.21 |
| 2216585 | 5.21 |
| 2221300 | 5.21 |
| 2224044 | 4.62 |
| 2230909 | 5.11 |
| 2244351 | 5.21 |
| 2269539 | 5.11 |
| 2445256 | 5.21 |
| 2477565 | 5.21 |
| 2480483 | 5.21 |
| 2498066 | 5.11 |
| 2536504 | 5.11 |
| 2554800 | 5.21 |
| 2581808 | 5.21 |
| 2620136 | 5.21 |
| 2697372 | 5.21 |
| 2705259 | 4.72 |
| 2716223 | 4.72 |
| 2717496 | 4.72 |
| 2720810 | 4.72 |
| 2725142 | 4.62 |
| 2742713 | 4.72 |
| 3014324 | 5.21 |
| 3042382 | 5.41 |
| 3044894 | 5.41 |
| 3050915 | 5.41 |
| 3063617 | 4.32 |
| 3125581 | 4.62 |
| 3133691 | 5.41 |
| 3145066 | 5.41 |
| 3151894 | 5.3 |
| 3206848 | 5.3 |
| 3218405 | 5.3 |
| 3224574 | 4.62 |
| 3239008 | 5.41 |
| 3254104 | 5.41 |
| 3263652 | 5.41 |
| 3285833 | 5.41 |
| 3699733 | 5.3 |
| 3723253 | 5.41 |
| 3727199 | 5.3 |
| 3735465 | 4.62 |
| 3746340 | 5.41 |
| 3747236 | 5.41 |
| 3750744 | 5.41 |
| 3751854 | 5.3 |
| 3759291 | 5.41 |
| 3763400 | 5.41 |
| 3769638 | 3.82 |
| 3771042 | 4.62 |
| 3780600 | 4.62 |
| 3787942 | 5.41 |
| 3803039 | 5.21 |
| 3813573 | 4.62 |
| 3833712 | 2.24 |
| 3852864 | 5.3 |
| 3864853 | 5.3 |
| 3889355 | 5.3 |
| 3894444 | 5.41 |
| 3921325 | 5.41 |
| 3929063 | 5.41 |
| 3952401 | 5.41 |
| 3999816 | 5.41 |
| 4016812 | 5.3 |
| 4062892 | 5.41 |
| 4080293 | 4.62 |
| 4097189 | 4.62 |
| 4225430 | 4.62 |
| 4252738 | 4.62 |
| 4285456 | 5.3 |
| 4315215 | 4.62 |
| 4345967 | 5.41 |
| 4394973 | 5.41 |
| 4416473 | 4.62 |
| 4422980 | 5.41 |
| 4434971 | 5.21 |
| 4443129 | 5.41 |
| 4474014 | 5.41 |
| 4516377 | 5.3 |
| 4525010 | 5.41 |
| 4534032 | 5.21 |
| 4547490 | 4.62 |
| 4558257 | 4.62 |
| 4579123 | 5.41 |
| 4588414 | 4.72 |
| 4597645 | 4.62 |
| 4604800 | 4.51 |
| 4625638 | 4.41 |
| 4648219 | 5.41 |
| 4669796 | 5.3 |
| 4680281 | 5.3 |
| 4686955 | 5.41 |
| 4697170 | 4.62 |
| 4736216 | 5.41 |
| 4741592 | 5.41 |
| 4755948 | 5.3 |
| 4826591 | 4.62 |
| 4841300 | 4.62 |
| 4857721 | 4.62 |
| 4869180 | 5.21 |
| 4878592 | 5.3 |
| 4901188 | 5.21 |
| 4936653 | 4.62 |
| 5004891 | 5.3 |
| 5025749 | 5.11 |
| 5044517 | 4.72 |
| 5097502 | 4.62 |
| 5101576 | 5.61 |
| 5141781 | 5.71 |
| 5147706 | 5.61 |
| 5151525 | 4.62 |
| 5195600 | 5.51 |
| 5227224 | 5.71 |
| 5244778 | 5.3 |
| 5267469 | 5.61 |
| 5274837 | 5.41 |
| 5276146 | 5.71 |
| 5294730 | 4.62 |
| 5295619 | 5.61 |

LG18

|  | genetic_pos |
| --- | --- |
| 970200 | 5.82 |
| 990298 | 4.73 |
| 1007260 | 4.55 |
| 1044370 | 5.64 |
| 1088947 | 5.82 |
| 1109414 | 5.82 |
| 1240692 | 5.55 |
| 1518561 | 5.82 |
| 1541061 | 5.64 |
| 1606504 | 5.27 |
| 1631807 | 5.82 |
| 1705951 | 5.55 |
| 1758484 | 5.55 |
| 1760393 | 5.82 |
| 1789531 | 4.73 |
| 1826137 | 5.55 |
| 1887004 | 5.18 |
| 1908343 | 5.46 |
| 1917952 | 5.82 |
| 1937015 | 5.82 |
| 1960453 | 5.82 |
| 1970531 | 5.55 |
| 1976872 | 5.82 |
| 1980396 | 5.82 |
| 1988730 | 5.55 |
| 2002196 | 5.82 |
| 2007631 | 4.73 |
| 2013863 | 5.82 |
| 2028883 | 5.82 |
| 2031683 | 5.82 |
| 2044901 | 5.18 |
| 2050842 | 5.55 |
| 2062850 | 4.82 |
| 2064416 | 4.21 |
| 2073123 | 5.82 |
| 2082504 | 5.27 |
| 2100785 | 5.55 |
| 2125025 | 5.55 |
| 2129607 | 6.18 |
| 2168784 | 5.55 |
| 2243851 | 5.82 |
| 2250350 | 5.55 |
| 2470517 | 5.82 |
| 2480205 | 5.55 |
| 2488670 | 5.55 |
| 2767938 | 5.55 |
| 2777247 | 5.82 |
| 2787381 | 5.55 |
| 2810651 | 4.73 |
| 2854839 | 5.55 |
| 2864926 | 4.73 |
| 2873470 | 5.55 |
| 2894378 | 5.55 |
| 2904478 | 3.08 |
| 2977256 | 4.73 |
| 2979203 | 5.55 |
| 2982860 | 5.55 |
| 2986276 | 5.46 |
| 2996319 | 5.55 |
| 2998353 | 5.82 |
| 3006192 | 5.55 |
| 3013801 | 5.27 |
| 3019148 | 5.82 |
| 3032155 | 5.55 |
| 3036456 | 5.82 |
| 3050741 | 4.73 |
| 3068721 | 5.18 |
| 3080079 | 5.55 |
| 3097546 | 5.55 |
| 3124016 | 5.55 |
| 3136668 | 5.55 |
| 3152221 | 4.73 |
| 3171884 | 5.55 |
| 3180379 | 4.73 |
| 3200503 | 5.55 |
| 3210433 | 5.82 |
| 3227033 | 5.82 |
| 3238626 | 5.55 |
| 3248369 | 5.55 |
| 3256500 | 5.82 |
| 3385938 | 6.45 |
| 3396575 | 5.55 |
| 3406786 | 5.55 |
| 3424627 | 5.55 |
| 3448334 | 5.82 |
| 3475507 | 5.55 |
| 3522376 | 5 |
| 3524590 | 5.55 |
| 3754791 | 5.55 |
| 3787249 | 5.55 |
| 3798012 | 5.55 |
| 3837495 | 5.82 |
| 3884059 | 5.82 |
| 3919994 | 5.55 |
| 4396436 | 4.73 |
| 4403188 | 4.73 |
| 4448646 | 4.73 |
| 4455641 | 4.21 |
| 4478009 | 5.18 |
| 4489268 | 5.55 |
| 4519494 | 5.82 |
| 4527246 | 5.55 |
| 4556234 | 5.82 |
| 4560298 | 5.55 |
| 4628668 | 5.55 |
| 4630323 | 5.55 |
| 4639207 | 5.55 |
| 4652948 | 4.73 |
| 4660404 | 5.55 |
| 4663126 | 5.55 |
| 4688687 | 5.55 |
| 4699527 | 5.55 |
| 4709437 | 5.55 |
| 4742582 | 5.55 |
| 4801686 | 4.73 |
| 4814151 | 5.82 |
| 4829844 | 5.55 |
| 4840009 | 5.55 |
| 4850057 | 5.82 |
| 4864761 | 4.73 |
| 4919666 | 5.55 |
| 4929048 | 5.64 |
| 4933520 | 4.82 |
| 4987784 | 5.18 |
| 5000765 | 4.82 |
| 5048241 | 4.73 |
| 5055587 | 4.73 |
| 5067307 | 5.46 |
| 5070515 | 5.55 |
| 5080017 | 5.09 |
| 5108362 | 5.82 |
| 5111867 | 4.82 |
| 5113600 | 5.82 |
| 5126344 | 5.82 |
| 5144704 | 5.55 |
| 5153782 | 5.55 |
| 5204694 | 5.82 |
| 5229360 | 5.46 |
| 5652018 | 5.46 |
| 5662748 | 5.82 |
| 5672291 | 5.18 |
| 5702237 | 4.82 |
| 5710077 | 5.82 |
| 5718357 | 4.73 |
| 5727645 | 5.55 |
| 5738508 | 6.27 |
| 5743606 | 6.18 |
| 5744530 | 5.55 |
| 5755695 | 5.55 |
| 5764428 | 5.55 |
| 5772846 | 5.82 |
| 5779843 | 5.82 |
| 5787654 | 6.18 |
| 5827723 | 5.09 |
| 5837432 | 5.55 |
| 5871504 | 5.91 |
| 5912111 | 5.91 |
| 5912819 | 6 |
| 5926589 | 6 |
| 5931355 | 5.91 |
| 5941923 | 6 |
| 5980321 | 6.18 |
| 5981466 | 6 |
| 5993013 | 4.82 |
| 6001443 | 5.55 |
| 6043667 | 4.73 |
| 6060505 | 6 |
| 6076321 | 5.55 |
| 6085854 | 5.91 |
| 6092857 | 5.55 |
| 6101856 | 5.55 |
| 6113441 | 6 |
| 6145998 | 5.55 |
| 6164759 | 5.55 |
| 6173171 | 6.09 |
| 6183464 | 6 |
| 6193567 | 5.55 |
| 6204633 | 5.55 |
| 6232934 | 6.27 |
| 6263838 | 6.09 |
| 6291454 | 6.27 |
| 6336987 | 6.27 |
| 6346540 | 6.27 |
| 6352101 | 6.27 |
| 6372785 | 6.45 |
| 6390615 | 6.36 |
| 6401945 | 6.18 |
| 6414523 | 6.27 |
| 6425885 | 6.27 |
| 6468207 | 6.27 |
| 6471671 | 6.27 |
| 6475127 | 6.36 |
| 6492076 | 6.27 |
| 6503977 | 4.73 |
| 6511393 | 6.27 |
| 6540995 | 4.82 |
| 6550602 | 6.27 |
| 6553920 | 6.27 |
| 6554650 | 6.36 |
| 6561427 | 6.27 |

LG19

|  | genetic_pos |
| --- | --- |
| 1817812 | 5.7 |
| 1826693 | 2.6 |
| 1855987 | 5.8 |
| 1857675 | 5.8 |
| 1877772 | 3.97 |
| 1898596 | 5.3 |
| 1907393 | 5.8 |
| 1916920 | 4.18 |
| 1955872 | 5.8 |
| 1966408 | 6.1 |
| 1977456 | 6.1 |
| 1984089 | 6.1 |
| 2000507 | 5.8 |
| 2002022 | 5.8 |
| 2070727 | 5.7 |
| 2098628 | 5.8 |
| 2111824 | 4.18 |
| 2114222 | 3.97 |
| 2119063 | 2.2 |
| 2136916 | 5.8 |
| 2152931 | 6.1 |
| 2155196 | 3.66 |
| 2157770 | 3.97 |
| 2160089 | 3.97 |
| 2183626 | 3.97 |
| 2190401 | 5.01 |
| 2193816 | 5.4 |
| 2222455 | 5.3 |
| 2235348 | 4.18 |
| 2243732 | 0 |
| 2252596 | 5.01 |
| 2276549 | 5.7 |
| 2323337 | 4.18 |
| 2336837 | 5.7 |
| 2342373 | 4.18 |
| 2355336 | 6.1 |
| 2414428 | 4.18 |
| 2428993 | 5.8 |
| 2430252 | 5.01 |
| 2440932 | 5.7 |
| 2459238 | 5.8 |
| 2477770 | 3.66 |
| 2500625 | 5.7 |
| 2518334 | 3.66 |
| 2521919 | 5.3 |
| 2538116 | 6.9 |
| 2545399 | 4.91 |
| 2646264 | 5.01 |
| 2654782 | 6.1 |
| 2699258 | 2.6 |
| 2700971 | 5.4 |
| 2705067 | 5.7 |
| 2740796 | 5.4 |
| 2756434 | 5.4 |
| 2826000 | 4.18 |
| 2843429 | 5.7 |
| 2845730 | 5.7 |
| 2958835 | 6.43 |
| 2964950 | 5.7 |
| 2966665 | 5.7 |
| 2976910 | 5.8 |
| 3403418 | 6.1 |
| 3414509 | 4.18 |
| 3458999 | 5.4 |
| 3474945 | 5.01 |
| 3503580 | 5.8 |
| 3517580 | 5.7 |
| 3526224 | 5.4 |
| 3535010 | 5.8 |
| 3550092 | 5.01 |
| 3556510 | 5.7 |
| 3576846 | 3.97 |
| 3606824 | 5.01 |
| 3611736 | 5.4 |
| 3622022 | 5.7 |
| 3627232 | 3.97 |
| 3635109 | 6.1 |
| 3638885 | 5.8 |
| 3656679 | 4.18 |
| 3657075 | 5.7 |
| 3715185 | 5.7 |
| 3718096 | 5.8 |
| 3724989 | 5.7 |
| 3735786 | 5.01 |
| 3737925 | 3.66 |
| 3739310 | 3.66 |
| 3745937 | 3.66 |
| 3747391 | 3.97 |
| 3754709 | 5.9 |
| 3778172 | 6.1 |
| 3790038 | 3.66 |
| 3790372 | 5.4 |
| 3790467 | 5.4 |
| 3797595 | 5.9 |
| 3809153 | 6.1 |
| 3819602 | 5.9 |
| 3828845 | 5.7 |
| 3836261 | 5.8 |
| 3842396 | 5.7 |
| 3846949 | 5.9 |
| 3886127 | 5.8 |
| 3905763 | 3.66 |
| 3912746 | 3.97 |
| 3915052 | 5.7 |
| 3915766 | 5.8 |
| 3930631 | 3.97 |
| 3947725 | 5.9 |
| 3966022 | 5.7 |
| 3973188 | 5.3 |
| 3983257 | 5.9 |
| 3984874 | 5.7 |
| 3985804 | 5.3 |
| 4034908 | 6.1 |
| 4056983 | 6.1 |
| 4057981 | 5.01 |
| 4066501 | 5.8 |
| 4068525 | 6.1 |
| 4071276 | 2.6 |
| 4074429 | 5.7 |
| 4081590 | 4.18 |
| 4085617 | 5.9 |
| 4096702 | 6.3 |
| 4109723 | 5.4 |
| 4116941 | 5.01 |
| 4118103 | 6.9 |
| 4124641 | 3.66 |
| 4132723 | 6.1 |
| 4147001 | 6.9 |
| 4166193 | 5.9 |
| 4173178 | 5.9 |
| 4182621 | 5.9 |
| 4198161 | 6 |
| 4199303 | 3.66 |
| 4204958 | 6.4 |
| 4227185 | 4.18 |
| 4246827 | 6 |
| 4250274 | 6 |
| 4254992 | 6 |
| 4257995 | 6 |
| 4263014 | 5.01 |
| 4292302 | 6.9 |
| 4316518 | 4.18 |
| 4333193 | 6.5 |
| 4339120 | 6.5 |
| 4342833 | 2.76 |
| 4353340 | 6.5 |
| 4370972 | 6.5 |
| 4381035 | 3.66 |
| 4390221 | 6.5 |
| 4401277 | 5.01 |
| 4447624 | 4.18 |
| 4479833 | 5.7 |
| 4490171 | 3.97 |
| 4519721 | 5.7 |
| 4565921 | 7.2 |
| 4578711 | 3.97 |
| 4591518 | 6.9 |
| 4598986 | 4.18 |
| 4603085 | 6.7 |
| 4606956 | 6.9 |
| 4613062 | 6.9 |
| 4619498 | 6.9 |
| 4632034 | 8.81 |
| 4642352 | 4.18 |
| 4649683 | 5.3 |
| 4678911 | 3.97 |
| 4687825 | 7 |
| 4701261 | 6.8 |
| 4704426 | 6.9 |
| 4711162 | 7 |
| 4720985 | 5.3 |
| 4729795 | 6.9 |
| 4738427 | 3.66 |
| 4750642 | 5.3 |
| 4757860 | 3.66 |
| 4769796 | 3.66 |
| 4793549 | 6.9 |
| 4801582 | 6.9 |
| 4802544 | 9.82 |
| 4806845 | 4.49 |
| 4827310 | 7 |
| 4831433 | 7 |
| 4840031 | 7 |
| 4842069 | 7 |
| 4842261 | 6.5 |
| 4859726 | 7 |
| 4883708 | 6.8 |
| 4894161 | 9.82 |
| 4902186 | 6.5 |
| 4904328 | 3.66 |
| 4919271 | 3.66 |
| 4950919 | 6.9 |
| 4962172 | 7 |
| 4963548 | 7 |
| 4964987 | 6.9 |
| 4967650 | 6.9 |
| 4968719 | 4.18 |
| 4970559 | 7 |
| 4975813 | 3.66 |
| 4993766 | 7 |

LG20

|  | genetic_pos |
| --- | --- |
| 23430 | 7.08 |
| 121160 | 7.17 |
| 131238 | 6.71 |
| 131727 | 6.43 |
| 149880 | 6.71 |
| 164475 | 6.8 |
| 175680 | 6.43 |
| 351603 | 6.99 |
| 367076 | 6.99 |
| 377415 | 6.43 |
| 400654 | 6.43 |
| 407855 | 6.71 |
| 418557 | 6.43 |
| 419662 | 6.43 |
| 427060 | 6.71 |
| 513841 | 6.43 |
| 515802 | 6.43 |
| 636805 | 7.17 |
| 645972 | 7.08 |
| 661276 | 7.08 |
| 662942 | 6.89 |
| 663899 | 6.99 |
| 677120 | 6.71 |
| 677992 | 6.99 |
| 683857 | 6.71 |
| 686422 | 6.43 |
| 689508 | 6.8 |
| 705200 | 6.43 |
| 706517 | 6.8 |
| 715174 | 6.71 |
| 716480 | 7.08 |
| 732019 | 6.99 |
| 736660 | 6.43 |
| 738984 | 6.8 |
| 801052 | 6.71 |
| 813669 | 6.99 |
| 821891 | 6.99 |
| 831871 | 6.43 |
| 843836 | 6.71 |
| 845216 | 6.43 |
| 845769 | 7.08 |
| 855173 | 6.99 |
| 874958 | 6.99 |
| 878644 | 6.43 |
| 881987 | 6.71 |
| 896117 | 6.71 |
| 918218 | 6.71 |
| 922316 | 6.99 |
| 922614 | 6.43 |
| 930910 | 6.8 |
| 931384 | 6.43 |
| 941626 | 3.85 |
| 950511 | 6.43 |
| 973515 | 6.99 |
| 984971 | 7.08 |
| 990392 | 6.99 |
| 995886 | 6.8 |
| 1020403 | 6.99 |
| 1030876 | 6.99 |
| 1031354 | 6.99 |
| 1049114 | 6.8 |
| 1056627 | 6.99 |
| 1062652 | 6.99 |
| 1076718 | 6.99 |
| 1085484 | 6.99 |
| 1086739 | 6.8 |
| 1115718 | 6.8 |
| 1151281 | 6.99 |
| 1157013 | 7.17 |
| 1162073 | 6.8 |
| 1367898 | 6.8 |
| 1387616 | 7.26 |
| 1404875 | 6.71 |
| 1408112 | 6.8 |
| 1410148 | 6.8 |
| 1417209 | 6.8 |
| 1426878 | 6.99 |
| 1435868 | 7.17 |
| 1443404 | 6.8 |
| 1446446 | 6.99 |
| 1457386 | 6.8 |
| 1460514 | 6.8 |
| 1463989 | 6.8 |
| 1482395 | 6.99 |
| 1566061 | 6.99 |
| 1599255 | 6.99 |
| 1611607 | 6.43 |
| 1666835 | 6.99 |
| 1675371 | 6.43 |
| 1699382 | 6.43 |
| 1839112 | 7.26 |
| 1847622 | 7.17 |
| 1853913 | 7.08 |
| 1859161 | 7.08 |
| 1860088 | 7.17 |
| 1860268 | 6.99 |
| 1863040 | 6.99 |
| 1872783 | 6.71 |
| 1873003 | 6.71 |
| 1875608 | 6.99 |
| 1888520 | 6.71 |
| 1896306 | 6.43 |
| 1901764 | 6.43 |
| 1908113 | 6.8 |
| 1913066 | 6.99 |
| 1918224 | 6.71 |
| 2153472 | 6.33 |
| 2161977 | 6.8 |
| 2256729 | 6.99 |
| 2275857 | 7.08 |
| 2279357 | 6.99 |
| 2283473 | 7.08 |
| 2291935 | 6.99 |
| 2297531 | 6.43 |
| 2332221 | 6.43 |
| 2374781 | 6.43 |
| 2384343 | 6.99 |
| 2394565 | 6.43 |
| 2423479 | 6.99 |
| 2437770 | 6.43 |
| 2444091 | 6.99 |
| 2450447 | 6.8 |
| 2464187 | 6.8 |
| 2471941 | 6.43 |
| 2482333 | 5.41 |
| 2488534 | 7.08 |
| 2524666 | 6.43 |
| 2528020 | 6.99 |
| 2529291 | 6.99 |
| 2533919 | 6.99 |
| 2541582 | 6.71 |
| 2562518 | 6.71 |
| 2573525 | 7.17 |
| 2593495 | 6.71 |
| 2604430 | 6.99 |
| 2612871 | 6.71 |
| 2625010 | 6.99 |
| 2650471 | 6.99 |
| 2651646 | 6.99 |
| 2652257 | 6.99 |
| 2659016 | 7.08 |
| 2661926 | 6.43 |
| 2669627 | 7.17 |
| 2672587 | 7.17 |
| 2673716 | 6.8 |
| 2711348 | 6.43 |
| 2743240 | 6.43 |
| 2753310 | 6.43 |
| 2807210 | 6.43 |
| 2842823 | 6.43 |
| 2852520 | 6.8 |
| 2866141 | 6.99 |
| 2881852 | 6.99 |
| 2885557 | 6.71 |
| 2895931 | 6.99 |
| 2904985 | 6.71 |
| 2917736 | 6.99 |
| 2923180 | 6.71 |
| 2968606 | 6.71 |
| 3053716 | 6.99 |
| 3063132 | 6.99 |
| 3067309 | 6.71 |
| 3074266 | 6.99 |
| 3075797 | 6.43 |
| 3080667 | 3.42 |
| 3089965 | 6.71 |
| 3110829 | 6.71 |
| 3122176 | 6.99 |
| 3130129 | 6.99 |
| 3137518 | 7.08 |
| 3139999 | 6.43 |
| 3166010 | 6.71 |
| 3168638 | 6.43 |
| 3192689 | 6.71 |
| 3195428 | 7.17 |
| 3204171 | 6.71 |
| 3213407 | 7.08 |
| 3229054 | 6.8 |
| 3237182 | 6.71 |
| 3244931 | 6.71 |
| 3252013 | 6.99 |
| 3255147 | 7.17 |
| 3260951 | 7.17 |
| 3263497 | 6.43 |
| 3274744 | 6.43 |
| 3275410 | 7.17 |
| 3279241 | 6.71 |
| 3279857 | 7.17 |
| 3282539 | 3.85 |
| 3398176 | 6.71 |
| 3409331 | 6.71 |
| 3409595 | 7.17 |
| 3415236 | 6.99 |
| 3428795 | 6.99 |
| 3488482 | 6.43 |
| 3559711 | 6.71 |
| 3764087 | 6.43 |
| 3769876 | 6.71 |
| 3781493 | 7.17 |
| 3800224 | 6.71 |

LG22

|  | genetic_pos |
| --- | --- |
| 21434 | 5.24 |
| 22041 | 3.06 |
| 23960 | 3.45 |
| 63865 | 4.86 |
| 76370 | 3.93 |
| 77473 | 5.42 |
| 82454 | 4.57 |
| 102331 | 3.93 |
| 103578 | 5.24 |
| 108787 | 5.24 |
| 118302 | 5.42 |
| 192940 | 4.3 |
| 354120 | 5.61 |
| 358922 | 4.86 |
| 421007 | 4.02 |
| 524095 | 3.93 |
| 530412 | 0 |
| 543082 | 4.57 |
| 543271 | 6.63 |
| 568940 | 3.64 |
| 573066 | 4.76 |
| 577226 | 3.93 |
| 581309 | 3.35 |
| 585971 | 3.93 |
| 592821 | 4.57 |
| 617133 | 4.57 |
| 638623 | 3.93 |
| 741366 | 4.76 |
| 753197 | 4.39 |
| 781309 | 4.39 |
| 800315 | 5.24 |
| 817127 | 4.39 |
| 850945 | 4.39 |
| 858016 | 4.39 |
| 868407 | 5.24 |
| 871809 | 5.24 |
| 879618 | 5.24 |
| 899796 | 4.02 |
| 905981 | 5.42 |
| 919408 | 4.39 |
| 929981 | 5.42 |
| 937565 | 5.42 |
| 947764 | 4.39 |
| 960406 | 5.24 |
| 969156 | 4.39 |
| 977294 | 4.02 |
| 1411868 | 4.57 |
| 1635995 | 5.24 |
| 1649741 | 5.24 |
| 1665813 | 3.35 |
| 1686666 | 5.05 |
| 1693522 | 4.57 |
| 1696704 | 4.86 |
| 1972156 | 3.16 |
| 1981098 | 5.42 |
| 1985074 | 5.05 |
| 2013346 | 4.39 |
| 2014895 | 3.06 |
| 2016180 | 3.93 |
| 2023672 | 2.96 |
| 2029909 | 3.64 |
| 2030801 | 4.57 |
| 2043897 | 5.05 |
| 2044295 | 5.24 |
| 2046706 | 4.39 |
| 2373793 | 5.24 |
| 2401836 | 4.76 |
| 2483399 | 4.57 |
| 2502434 | 4.39 |
| 2523717 | 4.39 |
| 2544335 | 4.39 |
| 2559627 | 4.39 |
| 2564943 | 3.93 |
| 2574940 | 5.24 |
| 2577169 | 5.42 |
| 2584744 | 4.39 |
| 2597879 | 4.02 |
| 2601300 | 4.57 |
| 2607232 | 4.02 |
| 2607925 | 4.76 |
| 2614030 | 5.05 |
| 2625347 | 4.02 |
| 2629719 | 4.76 |
| 2882526 | 4.39 |
| 2936700 | 5.24 |
| 2945400 | 5.24 |
| 3073492 | 5.05 |
| 3074106 | 4.76 |
| 3088796 | 4.39 |
| 3090115 | 5.42 |
| 3095014 | 5.24 |
| 3103321 | 5.05 |
| 3111296 | 4.76 |
| 3135973 | 5.24 |
| 3136213 | 4.86 |
| 3148830 | 4.02 |
| 3157651 | 5.05 |
| 3169712 | 4.39 |
| 3188558 | 5.05 |
| 3197878 | 4.39 |
| 3207553 | 5.24 |
| 3213432 | 3.35 |
| 3224466 | 4.39 |
| 3236621 | 4.57 |
| 3240301 | 4.02 |
| 3243458 | 4.39 |
| 3255738 | 4.39 |
| 3262724 | 4.76 |
| 3270840 | 4.39 |
| 3278744 | 4.39 |
| 3285046 | 4.3 |
| 3300823 | 4.57 |
| 3306401 | 3.45 |
| 3316826 | 5.42 |
| 3318696 | 3.54 |
| 3319026 | 4.57 |
| 3333103 | 4.57 |
| 3345245 | 5.42 |
| 3350510 | 4.39 |
| 3352335 | 4.39 |
| 3386952 | 4.39 |
| 3390050 | 4.57 |
| 3393452 | 5.24 |
| 3401147 | 3.93 |
| 3402530 | 4.39 |
| 3406568 | 4.39 |
| 3409056 | 5.51 |
| 3419959 | 5.51 |
| 3427020 | 4.11 |
| 3429886 | 4.39 |
| 3439572 | 4.76 |
| 3443397 | 5.51 |
| 3467581 | 5.42 |
| 3482387 | 5.61 |
| 3510457 | 5.7 |
| 3512056 | 5.7 |
| 3517866 | 4.39 |
| 3529246 | 5.7 |
| 3544619 | 3.16 |
| 6271290 | 3.93 |
| 6275122 | 4.39 |
| 6276332 | 5.89 |
| 6290793 | 5.89 |
| 6300896 | 5.89 |
| 6315286 | 4.02 |
| 6321180 | 4.39 |
| 6333866 | 5.98 |
| 6335425 | 7.19 |
| 6394706 | 5.89 |
| 6425154 | 5.7 |
| 6427797 | 5.89 |
| 6434999 | 5.8 |
| 6441876 | 5.89 |
| 6442537 | 4.2 |
| 6443711 | 5.61 |
| 6448929 | 5.89 |
| 6465585 | 3.93 |
| 6474711 | 3.16 |
| 6485167 | 4.02 |
| 6487482 | 3.06 |
| 6527780 | 4.39 |
| 6531747 | 4.39 |
| 6539504 | 4.02 |
| 6545199 | 5.89 |
| 6564582 | 5.89 |
| 6586798 | 5.89 |
| 6588532 | 6.92 |
| 6614615 | 5.14 |
| 6615062 | 5.89 |
| 6618172 | 5.89 |
| 6629692 | 7.47 |
| 6634757 | 4.02 |
| 6637982 | 5.42 |
| 6668549 | 5.98 |
| 6673672 | 6.08 |
| 6678288 | 5.98 |
| 6687283 | 6.08 |
| 6692546 | 5.89 |
| 6695672 | 5.89 |
| 6707742 | 5.98 |
| 6715275 | 4.02 |
| 6739019 | 2.59 |
| 6747226 | 7.47 |
| 6751175 | 6.08 |
| 6759892 | 5.98 |
| 6769856 | 6.08 |
| 6785504 | 5.98 |
| 6795655 | 6.17 |
| 6817297 | 7.47 |
| 6833903 | 5.89 |
| 6843384 | 5.8 |
| 6849188 | 4.39 |
| 6859942 | 6.26 |
| 6941154 | 6.08 |
| 6952800 | 5.98 |
| 6960439 | 5.7 |
| 6963424 | 5.51 |
| 6972837 | 4.39 |
| 6975233 | 4.02 |
| 6979861 | 6.26 |

LG23

|  | genetic_pos |
| --- | --- |
| 181167 | 5.15 |
| 233481 | 4.59 |
| 322243 | 5.15 |
| 377369 | 5.15 |
| 397778 | 5.15 |
| 406649 | 5.15 |
| 417462 | 5.15 |
| 500360 | 5.15 |
| 593374 | 4.32 |
| 598420 | 5.06 |
| 706749 | 5.15 |
| 772810 | 5.15 |
| 790025 | 5.15 |
| 790756 | 5.15 |
| 826156 | 3.28 |
| 842439 | 5.15 |
| 865902 | 5.15 |
| 878283 | 5.15 |
| 896406 | 5.15 |
| 946299 | 5.15 |
| 958584 | 5.15 |
| 960717 | 5.15 |
| 976427 | 4.59 |
| 982227 | 5.06 |
| 1092870 | 5.15 |
| 1102570 | 5.15 |
| 1151896 | 5.15 |
| 1158907 | 5.15 |
| 1160926 | 5.06 |
| 1208168 | 5.15 |
| 1225065 | 4.59 |
| 1236107 | 4.59 |
| 1337119 | 5.15 |
| 1347919 | 5.15 |
| 1391403 | 5.15 |
| 1442705 | 4.59 |
| 1454697 | 5.15 |
| 1461023 | 5.15 |
| 1585303 | 2.39 |
| 1604078 | 5.15 |
| 1875384 | 5.15 |
| 1877715 | 5.15 |
| 1908390 | 5.15 |
| 1919699 | 4.59 |
| 1964892 | 5.15 |
| 1975054 | 5.15 |
| 1976050 | 5.15 |
| 1994213 | 5.15 |
| 2006177 | 5.15 |
| 2113556 | 5.15 |
| 2138452 | 4.59 |
| 2145819 | 4.59 |
| 2298970 | 4.23 |
| 2352008 | 5.15 |
| 2353083 | 5.15 |
| 2370925 | 5.15 |
| 2373907 | 5.15 |
| 2374253 | 5.15 |
| 2390873 | 5.15 |
| 2493896 | 4.87 |
| 2605222 | 4.59 |
| 2625593 | 5.06 |
| 2659342 | 5.15 |
| 2665208 | 4.23 |
| 2667611 | 4.32 |
| 2676306 | 5.15 |
| 2679450 | 3.28 |
| 2698648 | 5.06 |
| 2708294 | 5.15 |
| 2716228 | 5.15 |
| 2725208 | 4.32 |
| 2727286 | 4.23 |
| 2767259 | 4.32 |
| 2784773 | 4.32 |
| 2792817 | 5.06 |
| 2804205 | 5.15 |
| 2811141 | 4.59 |
| 2816430 | 5.06 |
| 2816852 | 4.32 |
| 2821580 | 4.23 |
| 2825685 | 5.15 |
| 2828114 | 4.23 |
| 2829196 | 5.06 |
| 2836788 | 4.59 |
| 2851955 | 4.23 |
| 2863803 | 5.15 |
| 2869556 | 4.59 |
| 2886204 | 4.59 |
| 2954546 | 4.23 |
| 2973242 | 5.15 |
| 3113276 | 4.23 |
| 3114270 | 4.23 |
| 3115379 | 4.59 |
| 3121005 | 5.15 |
| 3275866 | 5.15 |
| 3285958 | 4.23 |
| 3301943 | 3.17 |
| 3307489 | 3.28 |
| 3316453 | 4.59 |
| 3324878 | 2.51 |
| 3348828 | 4.5 |
| 3433619 | 3.28 |
| 3434838 | 5.15 |
| 3465811 | 5.15 |
| 3467235 | 5.15 |
| 3483507 | 5.15 |
| 3523460 | 4.59 |
| 3549629 | 5.06 |
| 3567946 | 5.15 |
| 3599187 | 5.06 |
| 3662209 | 4.32 |
| 3684790 | 5.15 |
| 3690110 | 4.59 |
| 3701712 | 5.15 |
| 3731353 | 5.15 |
| 3765024 | 0 |
| 3806661 | 5.06 |
| 3813664 | 5.06 |
| 4854630 | 4.23 |
| 4862343 | 5.15 |
| 4874552 | 4.23 |
| 4883115 | 4.23 |
| 4893664 | 4.32 |
| 4902520 | 3.28 |
| 4921652 | 4.23 |
| 4924329 | 4.32 |
| 4931612 | 5.06 |
| 4943673 | 4.23 |
| 4948471 | 5.06 |
| 4949585 | 0.97 |
| 4959369 | 4.59 |
| 4969864 | 4.32 |
| 5041254 | 5.06 |
| 5050821 | 3.28 |
| 5063995 | 4.59 |
| 5091627 | 3.28 |
| 5123237 | 4.32 |
| 5142178 | 4.23 |
| 5155230 | 3.28 |
| 5170848 | 3.28 |
| 5197788 | 2.96 |
| 5203702 | 4.23 |
| 5219311 | 5.06 |
| 5228566 | 4.32 |
| 5251518 | 4.59 |
| 5272865 | 4.59 |
| 5283414 | 5.06 |
| 5289238 | 5.06 |
| 5514686 | 5.06 |
| 5526389 | 4.32 |
| 5533970 | 5.06 |
| 5606996 | 4.32 |
| 5627118 | 5.15 |
| 5660569 | 5.15 |
| 5678455 | 4.14 |
| 5710579 | 4.32 |
| 5718805 | 5.06 |
| 5728906 | 5.06 |
| 5736849 | 4.05 |
| 5749218 | 4.59 |
| 5750328 | 4.32 |
| 5752488 | 4.32 |
| 5766552 | 4.23 |
| 5779144 | 3.17 |
| 5796484 | 4.23 |
| 5828983 | 4.23 |
| 5842558 | 4.23 |
| 5866595 | 3.96 |
| 5873815 | 4.32 |
| 5898794 | 5.15 |
| 5927406 | 4.23 |
| 5950575 | 4.32 |
| 5968446 | 4.32 |
| 6014090 | 3.77 |
| 6019692 | 3.28 |
| 6068810 | 4.23 |
| 6072076 | 3.87 |
| 6101624 | 4.32 |
| 6106343 | 3.17 |
| 6107198 | 4.32 |
| 6144283 | 2.73 |
| 6157372 | 4.32 |
| 6165945 | 4.41 |
| 6166293 | 4.32 |
| 6217633 | 4.32 |
| 6240699 | 4.23 |
| 6241500 | 5.15 |
| 6253096 | 4.32 |
| 6283824 | 4.23 |
| 6283922 | 5.15 |
| 6285755 | 5.15 |
| 6286730 | 5.15 |
| 6292322 | 4.32 |
| 6297398 | 4.23 |
| 6299674 | 3.77 |
| 6308554 | 4.32 |
| 6310035 | 5.15 |
| 6314541 | 5.15 |
| 6316621 | 4.78 |
| 6323784 | 4.32 |
