## Supplementary figures and images for "Chromosome-level genome sequence of the Genetically Improved Farmed Tilapia (GIFT, *Oreochromis niloticus*) highlights regions of introgression with *O. mossambicus*"

### Supplementary_Figure_2_02.pdf

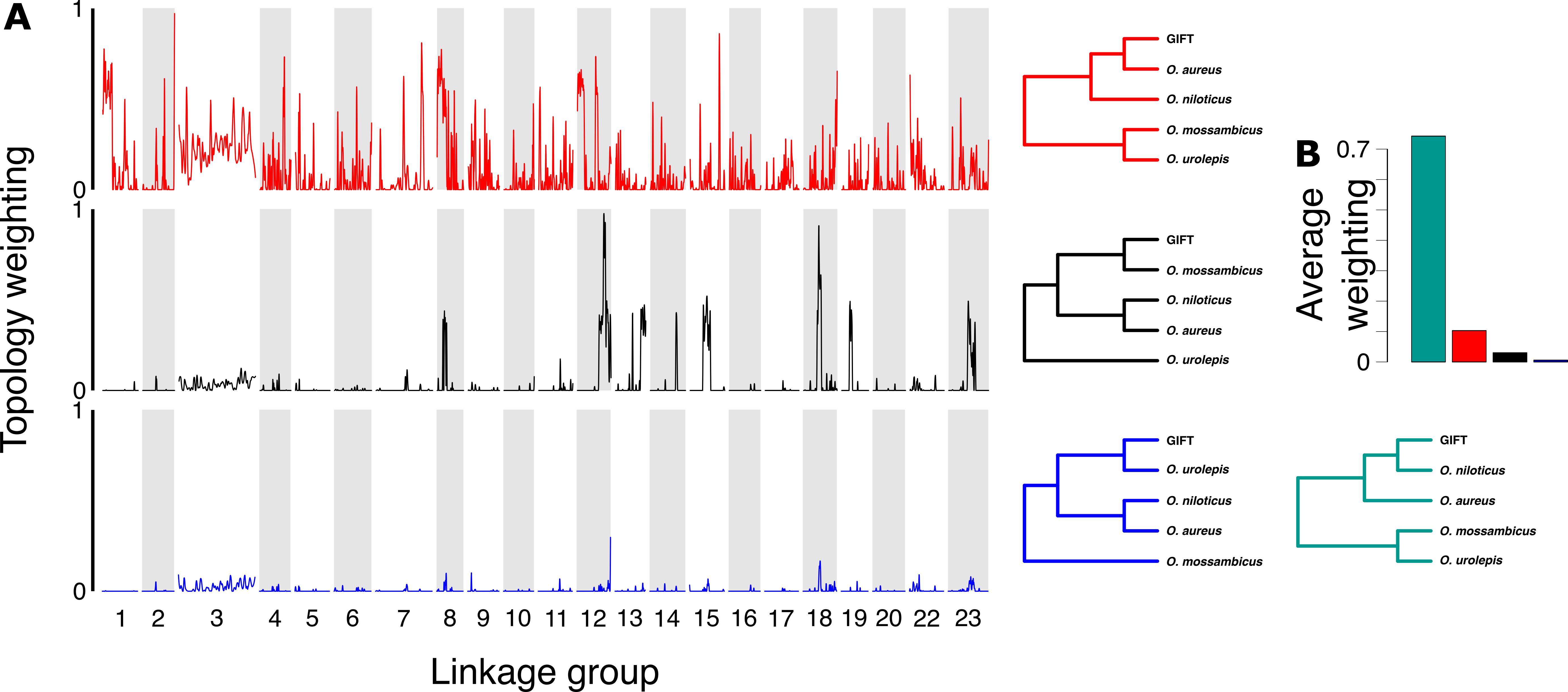

### Supplementary_Figure_3.pdf

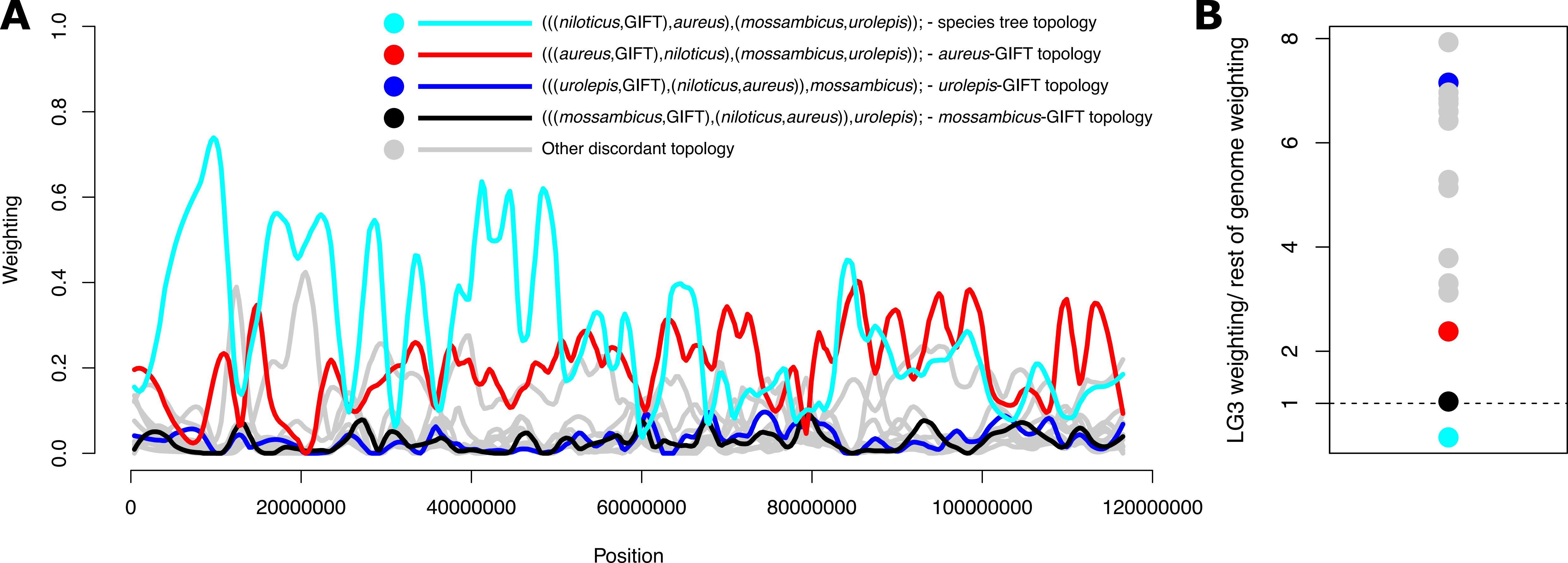

### Supplementary_Figure_4.pdf

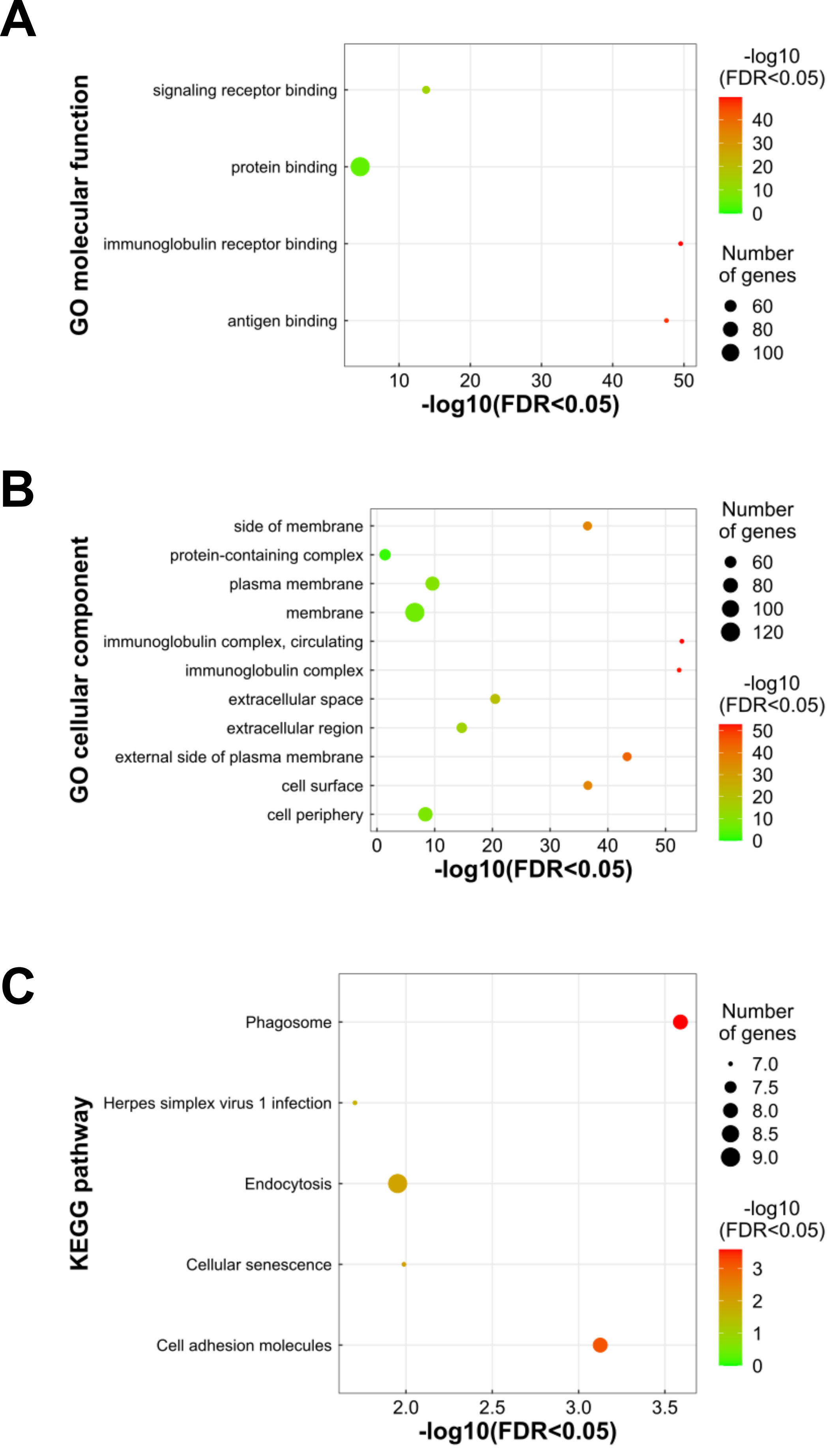
